# Specialized S-type ribosomes of *Plasmodium yoelii* enhance host-to-vector malaria transmission

**DOI:** 10.1101/2024.11.14.623520

**Authors:** James P. McGee, Sylvie Briquet, Olivier Silvie, Scott E. Lindner

## Abstract

Unlike most eukaryotes, *Plasmodium* species only encode 4-5 ribosomal DNA loci, making them an exceptional model system for genetic studies of ribosome specialization. *Plasmodium* ribosomes are characterized as A-type and S-type, defined by differing temporal expression patterns and sequence variation. The two S-type rRNAs (S1 & S2) are most abundant in developing mosquito stages, yet maintain low levels of abundance in blood stages. Two previous, conflicting studies found that one or either S-type rRNA was essential to mosquito-stage development, but these experiments were hampered by technical constraints. Therefore, we used the DiCre recombinase to generate transgenic parasites with clean deletions of the S1, S2, or both S-type rDNAs (S-type null) to characterize their roles and interplay in both mosquito- and blood-stage parasites. Contrary to previous conclusions, the presence of either or both S-type rRNAs was not required for sporozoite development, yet promoted host-to-vector transmission, oocyst maturation, and productive sporogony. Unexpectedly, we found that S-type ribosomes have different, opposing impacts on blood stage development despite being at low abundances. Deletion of S1 rDNA nearly ablated the first wave of parasitemia without impacting transmissible male gametocyte counts. Reciprocally, the deletion of S2 rDNA reduced counts of transmissible male gametocytes without impacting parasitemia, which was phenocopied by the S-type null line. Because the LSU portion of S1 rDNA could not be deleted, we assessed the S-type SSU rDNAs for potentially dominant roles. Introduction of extra plasmid-based copies of either S1 or S2 SSU rDNA resulted in either increased parasitemia or male gametogenesis, respectively. Yet, both decreased transmission and impacted early mosquito-stage development. These results indicate that driving functions of *Plasmodium’s* S-type ribosomes occur during blood stages, with S-type SSUs contributing distinct, separable functions. To the best of our knowledge, this is the first example of specialized ribosomes driving cellular outcomes at low abundances.

**Author Summary:** Malaria parasites (*Plasmodium* spp.) have specialized ribosomes with distinct temporal expression patterns and ribosomal RNA (rRNA) sequence variation. While previous work has demonstrated the importance of the S1 or S2 types of ribosomes to mosquito-stage development, technical limitations led to conflicting conclusions and prevented the investigation of whether the S-type ribosomes were essential for parasite development. Here, we used a DiCre recombinase system to overcome past limitations and generated clean deletions of both S-type rDNA genes, as well as the first *Plasmodium* line with both S-type rDNA genes deleted. Surprisingly, we found that *Plasmodium yoelii* parasites do not require either S-type ribosome for mosquito-stage development, when these rRNAs are most abundant. Instead, we found that S-type ribosomes contribute distinct functions during blood-stage development when these ribosomes make up only a small fraction of the total ribosomes present. In these stages, we found that S1 and S2 ribosomes have different, opposing functions that work cooperatively to promote transmission from the mammalian host to the mosquito vector.

## Introduction

Ribosomes are the fundamental translational machines of the cell, created from highly modified and processed ribosomal RNAs (rRNAs) and a large collection of Ribosomal Proteins (RPs) and Ribosome Associate Proteins (RAPs). In eukaryotes, the 45S pre-rRNA is transcribed from a canonical ribosomal DNA (rDNA) locus, which is then cleaved to form the 18S rRNA for use in the small subunit (SSU) of the ribosome and the 5.8S and 28S rRNAs for use in the large subunit (LSU) along with the 5S rRNA that is transcribed separately (1). Eukaryotes typically code for 100’s to 1000’s of copies of this rDNA locus organized in tandem repeat clusters to meet the translational needs of the cell (2). This repetitive genomic arrangement promotes the widely held idea that all ribosomes are functionally and compositionally uniform. However, some early and more recent studies rightly noted that the individual ribosomes produced from these rDNAs can produce distinctly different ribosomes due to factors including different RP compositions (3, 4, 5, 6) and rRNA sequence variation (7, 8, 9). This heterogeneous nature therefore allows ribosomes to have variable functions and capabilities in translation regulation consistent with the concept of ribosome specialization (reviewed in (10)). For example, the presence of specific RP paralogs in ribosomes has been shown to promote distinct cellular outcomes, including sperm formation in mouse germ cells (11) or human and mice embryonic stem cells differentiating down the mesoderm pathway (12).

However, ribosome specialization is challenging to investigate with reductionistic genetic approaches because of the large number of tandem repeats of rDNAs in most eukaryotes. Notably, a few eukaryotes are tractable for such studies, as they retain only a small number of rDNA loci, yet have two distinct ribosome types characterized by both rRNA sequence variation and temporal abundance pattern. This includes the metazoan *Danio rerio* (zebrafish) which produces maternal and somatic ribosomes (13), and the eukaryotic *Plasmodium* parasites that cause malaria. In both cases, zebrafish and *Plasmodium* position the different rDNA types on separate chromosomes, providing unique genetic systems for studying ribosome specialization (13, 14, 15, 16). Together, these systems provide opportunities to study and compare specialized ribosomes in the context of a metazoan and in a single-celled protozoan eukaryote.

*Plasmodium* expresses minimally two types of ribosomes that are temporally controlled, with the A-type most abundant in the mammalian host during liver and blood stages and the S-type most abundant in late-mosquito stage parasites (15, 17, 18) (reviewed in (19)). Importantly, both ribosome types are expressed throughout the life cycle, with low levels of S-type rRNA present when the A-type predominates and vice versa (20, 21, 22, 23, 24, 25, 26). These ribosome types are further characterized by rRNA sequence variation, where similar trends occur between the human infectious *P. falciparum* and the rodent malaria models *P. berghei* and *P. yoelii* (14, 27, 28). *P. berghei* and *P. yoelii* have two A-type rRNAs (historically called the A and B Units) located on Chr7 and Chr12 that maintain near 100% sequence identity to one another. These A-type rRNAs have sequence variation when compared to the S1 rRNA (historically called the C Unit, found on Chr5) and S2 rRNA (D Unit, found on Chr6). Furthermore, the S-type rRNAs have sequence variation between each other, as the S1 rRNA is a chimera of the A-type and S2 (27, 28).

Several studies have focused on the regulation and importance of these distinct rRNAs. One investigation found that the *Plasmodium falciparum* A and S2 rRNAs can be regulated by temperature and glucose concentrations (20, 21), two factors that differ substantially between mammalian and mosquito hosts. In addition, the essentiality of the S-type ribosomes has been investigated in two related rodent malaria species, although discrepancies exist in their results. In *P. berghei*, neither S-type ribosome type was essential for sporozoite development, leading to the hypothesis that only one S-type ribosome was required in a dose-dependent manner to allow for proper mosquito-stage development (18). In contrast, deletions within the S2 rDNA in *P. yoelii* led to oocysts that did not mature, indicating that the S2 rRNA was essential for full development of mosquito-stage parasites (29). Furthermore, the study involving *P. berghei* noted that the A-type ribosomes could not individually be deleted in blood-stage parasites, but did not provide supporting data for this conclusion (18). Taken together, these data led the authors to conclude that both ribosome types are essential to the life cycle stages in which they are most abundant. However, both studies with *P. berghei* and *P. yoelii* had technical limitations that enabled the deletion of only one rDNA locus per parasite line and that the deletion approaches used did not create clean deletions of the entire targeted rDNA locus.

In this study, we used *P. yoelii* as a genetically tractable model to better understand specialized ribosomes. By incorporating the DiCre recombinase system in *P. yoelii* 17XNL parasites to enable ligand-controlled, sequential genome editing, we generated clean deletions of all individual rDNA loci, as well as the first *Plasmodium* parasite line lacking both S-type rDNAs. Our results show that contrary to previous conclusions, S-type ribosomes are not required for sporozoite development. Instead, we found that the S1 and S2 ribosomes have distinct and opposing functions in blood-stage growth and development that impact parasite transmissibility, even when they are present at low levels compared to the predominant A-type ribosomes. Our results are consistent with S-type ribosomes acting as specialized ribosomes that contribute to the parasite’s response to being introduced into the mammalian host and how it prioritizes rapid increases in parasite numbers versus sexual differentiation to promote onward transmission to the mosquito.

## Results

### *Plasmodium yoelii* S-type rDNA loci are dispensable in blood-stage parasites

Prior investigations into S-type ribosome essentiality in *P. berghei* and *P. yoelii* demonstrated that sequences within S-type rDNA loci could be individually deleted in blood-stage parasites (18, 29). The discrepancy between these studies exists in the phenotypes observed in mosquito-stage parasites with deletions of the S1 or S2 rDNA (also called the C- and D-loci, respectively), which could have arisen due to the different gene targeting strategies used. Moreover, these studies could only target one S-type locus at a time due to technical constraints, and neither study cleanly deleted the entire rDNA locus but instead disrupted the transcription or processing of the pre-rRNA. To address this apparent discrepancy and overcome past technical limitations, we leveraged a dimerizable Cre recombinase (DiCre) system in *Plasmodium yoelii* to first create clean deletions of the entire rDNA of either S-type ribosome. This system allows for the recycling of the selectable marker for pyrimethamine resistance, allowing us to then target the second S-type locus for subsequent deletion.

To delete either the S1 or S2 rDNAs, we used conventional reverse genetics targeting upstream and downstream of the entirety of either S-type rDNA locus. The template provided for recombination contained a deletion cassette that conferred pyrimethamine resistance and constitutive GFP expression flanked by LoxP sites, further flanked by homology arms specific to either S1 or S2 (illustrated in Fig S1). Linearized plasmids were transfected into a *P. yoelii* line that had constitutive DiCre and mCherry expression (30). Similar to previous studies, we were able to readily delete the S-type rDNAs individually in blood-stage parasites. Transgenic parasites were selected for using pyrimethamine and confirmed by genotyping PCR, and enriched for using FACS for GFP+ and mCherry+ parasites. These enriched populations of transgenic parasites were then treated with rapamycin to excise the DNA encoding pyrimethamine resistance and GFP expression to generate parasites with clean deletions of the entire S1 or S2 rDNA locus, hereby called PyΔS1 or PyΔS2 (Fig S1A and B). PyΔS1 and PyΔS2 parasites were cloned via limiting dilution, and upon genotyping confirmation, two clones per line were selected for future experimentation (PyΔS1 clones 1.1 and 2.1; PyΔS2 clones 1.2 and 1.3).

Excision of the deletion cassette in our PyΔS1 and PyΔS2 parasite lines allowed for the reuse of the selectable marker for pyrimethamine for a subsequent genetic modification. PyΔS1 clone 1.1 was transfected with the linearized plasmid containing the deletion cassette with homology arms for S2. An identical workflow as described above was followed and we generated a clean deletion of the S2 rDNA locus in our ΔS1 clone 1.1, thereby generating the first *Plasmodium* parasite line null of either S-type rDNA, hereby called PyΔS1ΔS2 (Fig S1C). Our PyΔS1ΔS2 parasites were cloned via limiting dilution, confirmed via genotyping, and two clones were selected for future experimentation (PyΔS1ΔS2 clones 1.2 and 1.3).

### *Plasmodium yoelii* A-type rDNA loci can also be deleted in blood-stage parasites

It is generally assumed that the highly abundant A-type rRNAs of asexual blood stage parasites are essential, which aligns with a previous unsuccessful attempt to delete either A-type rDNA locus in *P. berghei* (18). To test this assumption, we targeted the two A-type loci in *P. yoelii* independently by using homology arms upstream and downstream specific to the rDNA locus located on either Chr7 or Chr12. Surprisingly, the individual A-type rDNA loci could be deleted. Genotyping revealed that one out of four transfection populations included transgenic parasites with the respective A-type locus deleted (Fig S2); however, these transgenic parasites were rapidly outcompeted by remaining wild-type parasites in the population with continued passaging. This demonstrated that while either A-type locus could be individually deleted, the resulting decreased abundance of the A-type ribosome in blood-stage parasites substantially decreased parasite fitness.

### Neither *Plasmodium yoelii* S-type ribosome is required for sporozoite development

Discrepancies between previous studies of *Plasmodium’s* S-type ribosomes drew different conclusions regarding their essentiality to mosquito-stage parasite development when they are most abundant. Therefore, we investigated S-type ribosome essentiality in mosquito-stage parasites by phenotyping mosquito-stage parasite development of our PyΔS1, PyΔS2, and PyΔS1ΔS2 parasite lines.

Transmission of the DiCre parental or S-type deletion transgenic parasites to *A. stephensi* mosquitoes occurred at the first observed increased wave of male exflagellation counts during mouse blood-stage infection, as identified by phase-contrast live microscopy of wet mounts. Mosquito midguts were dissected seven days post-blood meal to count the percent of mosquitoes infected (prevalence of infection) and the number of oocysts per infected midgut (intensity of infection) by mCherry fluorescence (Fig 1A, Fig S3A). Over three biological replicates, the datasets from transmissions that had outlying, large decreased differences in both prevalence and intensity of infection attributable to technical issues were excluded (Data S1). Therefore, PyΔS1 clone 1.1 and PyΔS1ΔS2 clone 1.3 are represented by only two biological replicates. The prevalences of infection for all S-type deletion lines were comparable to the DiCre control, indicating that an S-type ribosome is not required to transmit to and establish mosquito infection. Both individual S-type deletion lines had decreased intensity of mosquito infection, although the most pronounced decrease was observed in the PyΔS2 parasites. PyΔS1ΔS2 parasites did not have this pronounced decrease, but rather had a similar intensity of infection to the PyΔS1 parasites (Fig 1A). These data indicate that the presence of the S1 ribosome in the absence of the S2 ribosome leads to a larger decrease in either parasite transmission or early mosquito-stage development.

**Figure 1:**
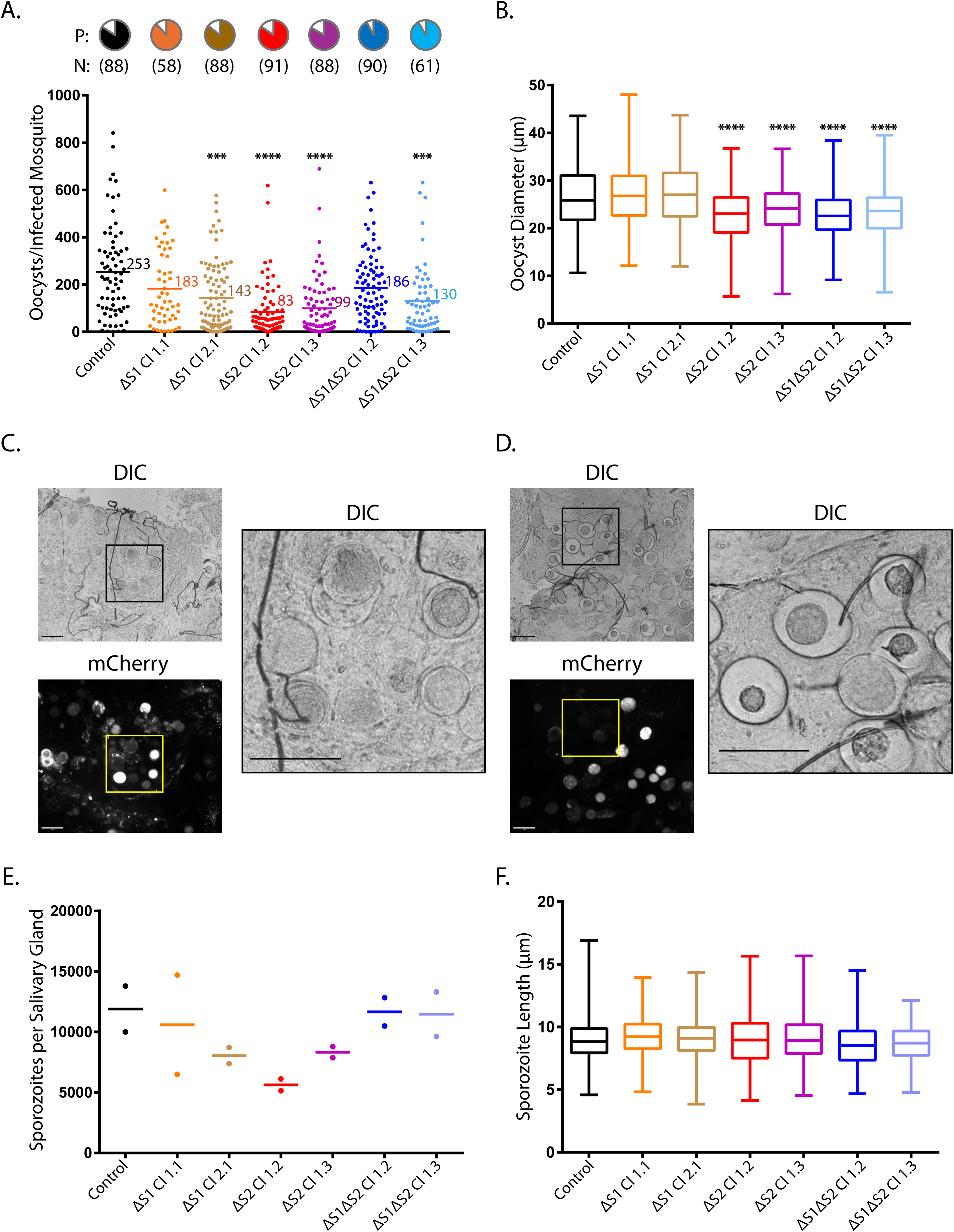
S-type ribosomes differentially impact mosquito-stage development but are not essential for sporozoite development. Mosquitoes took an infectious blood meal from mice infected with clonal transgenic parasites of ΔS1, ΔS2, or ΔS1ΔS2 along with the parental PyDiCre as a control. On day 7 after the blood meal, live fluorescence microscopy of dissected mosquito midguts was used to determine the following: **A)** Prevalence of mosquito infection per line is depicted by pie charts with N indicating the total number of mosquitoes dissected across three biological replicates. Intensity of infection is plotted per line with the average number of oocysts per midgut. Kruskal-Wallis test for multiple comparisons with Dunn’s Correction was used for statistical analysis (***p<0.001; ****p<0.0001). **B)** Day 7 oocyst diameters were measured in ImageJ with n = 100 oocysts measured for each replicate. Kruskal-Wallis test for multiple comparisons with Dunn’s Correction was used for statistical analysis (****p<0.0001). Representative day 14 midguts of **C)** control and **D)** ΔS1ΔS2 clone 1.2 imaged with DIC and live fluorescence microscopy using a 20x objective is shown (scale bar = 50µm). **E)** The sporozoite load per salivary gland for each line on day 14 was measured across two biological replicates (n = 10-20 salivary glands per replicate). **F)** Sporozoite lengths of control and S-type deletion lines were measured in ImageJ using fluorescent microscopy images (n = 50-75 per replicate for three biological replicates).

Both past studies on S-type ribosomes in *P. berghei* and *P. yoelii* reported decreased oocyst diameters when S2 was deleted (18, 29). We similarly measured oocyst diameters of mosquito midguts dissected seven days post-blood meal using DIC and live fluorescence microscopy using a 10x objective (Fig S3A). We observed that the diameters of parasite lines lacking an S2 ribosome (PyΔS2 and PyΔS1ΔS2) were, on average, 3-4µm smaller than control oocysts and that this difference was statistically significant (Fig 1B). We therefore followed oocysts through maturation until day fourteen post-blood meal (Fig S3B). Healthy, matured oocysts that had released sporozoites or were undergoing sporogony were observed in the control line and PyΔS1 (Fig 1C). In both parasite lines lacking the S2 ribosome, PyΔS2 and PyΔS1ΔS2, we observed healthy, mature oocysts but also observed an increased frequency of oocysts that failed to mature. These were identified by dense, circular structures resembling the remnants of sporoblasts that failed to produce sporozoites, which also no longer expressed mCherry, indicating these were non-viable/dead parasites (Fig 1D).

Despite the presence of these failed oocysts, all S-type deletion lines produced sporozoites that migrated to the salivary glands, demonstrating that neither S-type ribosome is essential for sporozoite development (Fig S4). Notably, the S-type deletion lines had no statistically significant decrease in salivary gland sporozoite loads and the small differences observed reflect the intensities of infections (Fig 1A and E). Finally, sporozoite length was then used as an initial indicator of proper sporozoite development and morphology. The lengths of parental and transgenic salivary gland sporozoites were not statistically different, indicating that sporozoite segmentation is not impacted by S-type ribosomes (Fig 1F). These data demonstrate that individual S-type ribosomes differentially impact mosquito stage development, yet neither is required.

### *Plasmodium yoelii* S-type ribosomes have opposing functions in blood-stage parasites

Because the S-type ribosomes are not essential for mosquito-stage parasite development when they are most abundant (15, 17, 18, 22, 23, 24, 25, 26), we sought to determine why *Plasmodium* would have evolved these distinct rRNAs. We hypothesized that the S-type ribosomes would play a functional role at another life cycle stage, even when they are present at low abundances. Therefore, we focused on blood-stage parasites to determine if S-type ribosomes impact transmissible parasites, which could explain the decrease in mosquito infection intensity (Fig 1A). Mice were infected intravenously with 1000 parasites of our S-type deletion lines or PyDiCre as our parental control, and parasites were phenotyped by counting blood-stage parasitemia and male gametogenesis daily throughout infection. Deletion of the S1 rDNA resulted in the near ablation of the first wave of blood-stage parasitemia, as well as significantly lower parasitemia throughout infection (Fig 2A). Despite the loss of the first wave of parasitemia, PyΔS1 parasites maintained the first wave of male gametogenesis and even gained an additional wave compared to the control, although this was only statistically significant for PyΔS1 clone 1.1 (Fig 2B). Contrastingly, deletion of the S2 rDNA did not affect blood-stage parasitemia (Fig 2C) but decreased the first wave of male gametogenesis (Fig 2D). These data reveal that the S-type ribosomes have opposing functions in blood-stage parasites even when the S-type rRNAs are lowly abundant.

**Figure 2:**
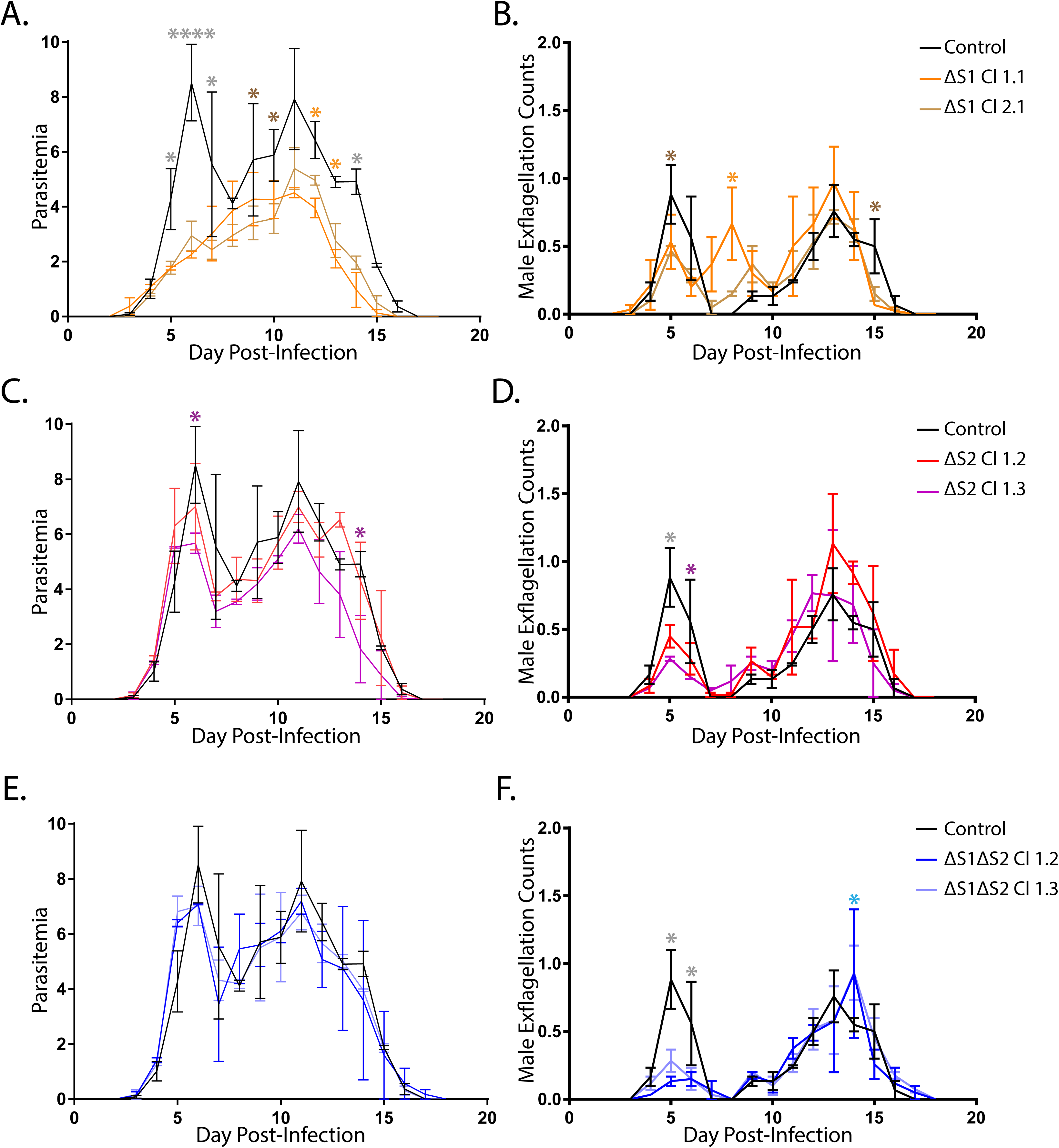
S-type ribosomes have differential, opposing functions in blood-stage parasites. *P. yoelii* blood-stage growth curves were performed by infecting mice with 1000 parasites intravenously. Two biological replicates were performed in technical triplicate. Blood-stage growth and male gametogenesis of ΔS1 **(A, B)**, ΔS2 **(C, D)**, ΔS1ΔS2 **(E, F)** clonal parasites were compared to PyDiCre control. Error bars indicate standard error of the mean across the means of two biological replicates provided in Data S1. A statistical analysis was conducted using Student’s t-test comparing the means per day to the control (*p<0.05; ****p<0.0001).

In addition, we observed that the PyΔS1ΔS2 parasites phenocopied the PyΔS2 parasites, with no effect on blood-stage parasitemia (Fig 2E) and a decreased first wave of male gametogenesis (Fig 2F). Because we generated the PyΔS1ΔS2 parasites by sequentially deleting S2 rDNA in the PyΔS1 parasites (Fig S1C), we concluded that PyΔS1ΔS2 parasites rescued the observed PyΔS1 decreased parasitemia at the cost of decreased male gametogenesis. This indicates the PyΔS1 phenotype is due to the presence of the S2 ribosome in the absence of the S1 ribosome, and by extension we hypothesize that the S1 ribosome works to maintain the increased first wave of blood-stage parasitemia.

### Additional plasmid-based copies of S1 or S2 SSU rDNA produce opposing phenotypes

To determine what rRNA segment of the S1 rRNA contributes to the function producing the first wave of blood-stage parasitemia in wild-type parasites (Fig 2A), we first took a progressive gene truncation approach to remove the 28S, or the 5.8S and 28S sequences of S1 rDNA. Although we could readily delete the entire S1 rDNA locus, we could not delete only these portions (Fig S5). Because these deletions would leave S1 18S expressed and conceptually able to produce a functional small subunit (SSU) of the ribosome, we reasoned that the remaining 18S SSU segment may be playing a functional role that could lead to a defect in asexual parasite growth. To assess this possibility, we took a reciprocal approach using plasmids containing only the promoter and ETS-18S-ITS-5.8S sequences of either S-type rDNA, as was done previously for S2 (29). These S-type SSU rDNA plasmids and a control plasmid without rDNA sequences were transfected into DiCre parasites and were continuously selected with pyrimethamine to promote plasmid retention (Fig S6A). Genotyping PCR and GFP expression via live fluorescence microscopy (Fig S6B and C) confirmed that these parasites harbor these plasmids.

Mice were infected intravenously with 1000 parasites per plasmid-containing parasite line and were phenotyped as described above. Parasites containing the S1 SSU rDNA plasmid had higher trending blood-stage parasitemia and a statistically significant increased and prolonged second wave of infection (Fig 3A) but had a decreased first wave of male gametogenesis (Fig 3B). Parasites containing the S2 SSU rDNA plasmid had earlier waves of blood-stage parasitemia occur (Fig 3A) and had significantly increased waves of male gametogenesis (Fig 3B). Introducing additional copies of either S-type SSU rDNA, therefore, led to opposing phenotypes in blood-stage parasitemia and male gametogenesis.

**Figure 3:**
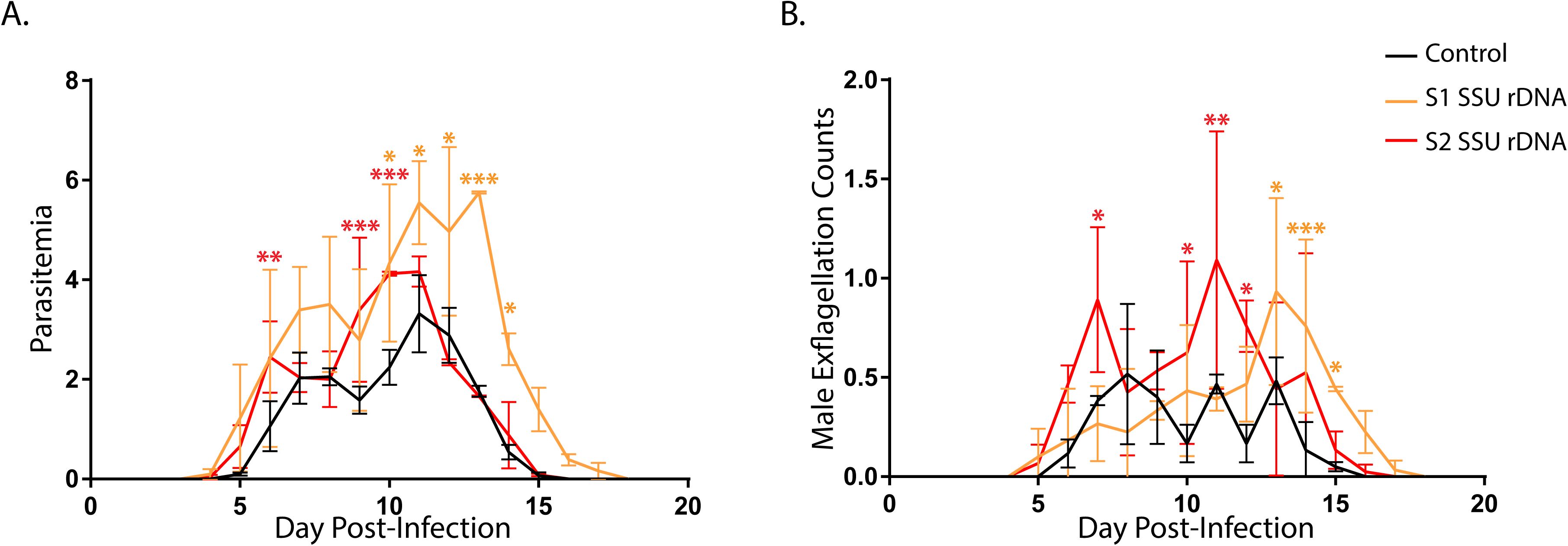
Additional plasmid-based copies of S1 or S2 SSU rDNA produces opposing phenotypes. *P. yoelii* blood-stage growth curves were performed by infecting mice with 1000 parasites intravenously. Two biological replicates were performed in technical triplicate. **A)** Blood-stage growth and **B)** male gametogenesis counts of parasites containing plasmids with S1 or S2 SSU rDNA were compared to the control plasmid (that lacks any rDNA sequences) over the course of mouse infection. Error bars indicate standard error of the mean across the means of two biological replicates provided in Data S1. A statistical analysis was conducted using Student’s t-test comparing means per day to the control (*p<0.05; **p<0.01; ***p<0.001).

### Additional plasmid-based copies of S-type SSU rDNA impact host-to-vector transmission and early mosquito-stage development

Because parasites containing either S-type SSU rDNA plasmid had significant differences in blood-stage parasitemia or male gametogenesis, we hypothesized that host-to-vector transmission would also be impacted. To address this, we performed host-to-vector transmission of plasmid-containing blood-stage parasites to *A. stephensi* mosquitoes. Mosquito midguts were dissected seven days post-blood meal to count both the prevalence and intensity of infection. Transmission of parasites containing either S-type SSU rDNA plasmid resulted in decreased prevalence of infection (not statistically significant) and significantly decreased the intensity of infection, which was more pronounced when transmitted parasites contained the S1 SSU rDNA plasmid (Fig 4A). This indicates that additional copies of either S-type SSU rDNA sequence led to decreased parasite transmissibility.

**Figure 4:**
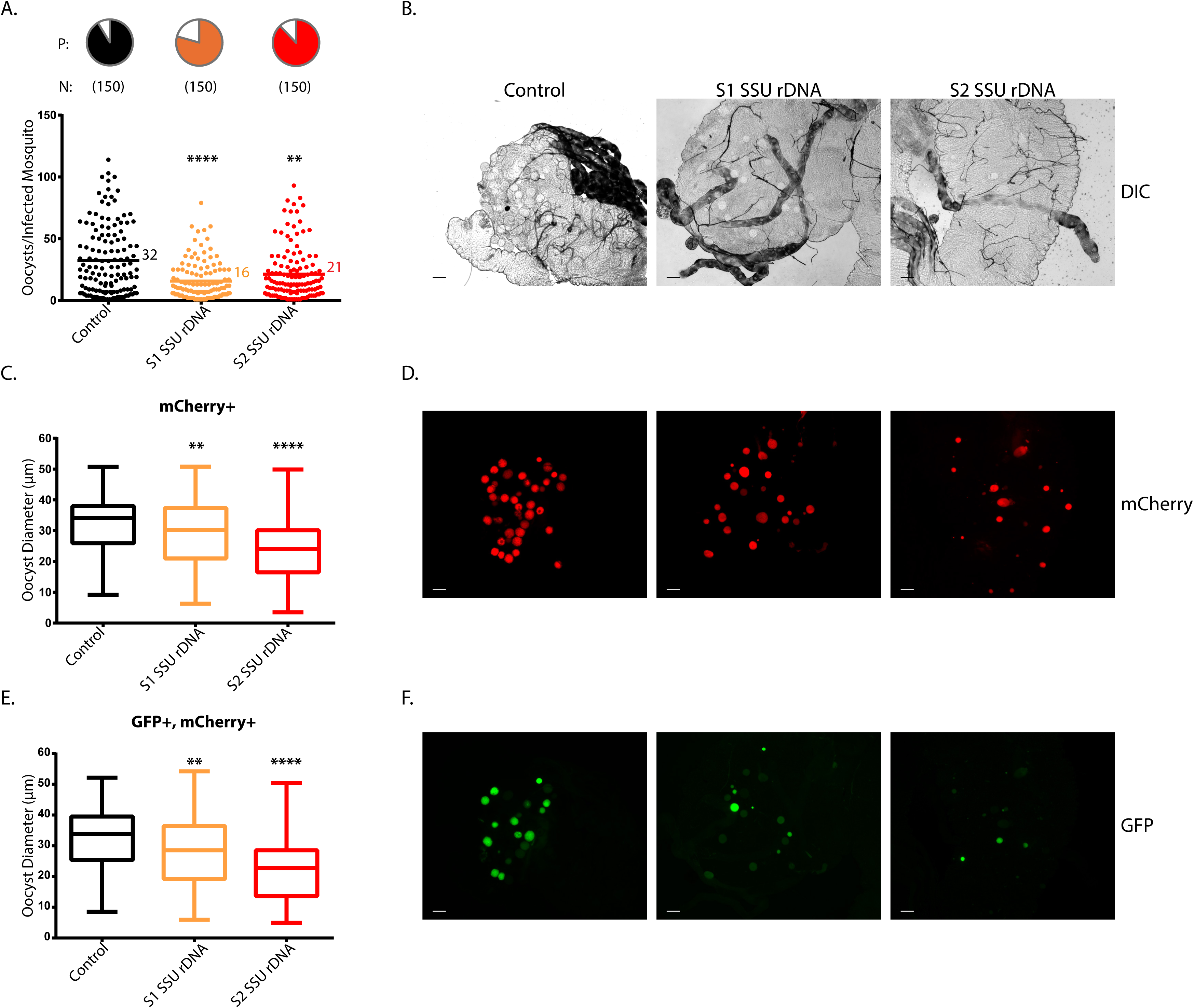
Additional plasmid-based copies of S-type SSU rDNA decreases host-to-vector transmission and early mosquito-stage development. Mosquitoes took an infectious blood meal from mice infected with PyDiCre parasites transfected with a control, S1 SSU rDNA, or S2 SSU rDNA plasmid. On day 7 after a blood meal, live fluorescence microscopy of dissected mosquito midguts was used to determine the following: **A)** Prevalence of mosquito infection per line is depicted by pie charts with N indicating the total number of mosquitoes dissected across three replicates. Intensity of infection is plotted per line with the average number of oocysts per midgut noted. Kruskal-Wallis test for multiple comparisons with Dunn’s correction was used for statistical analysis (**p<0.01; ****p<0.0001). **B)** Representative images of Day 7 midguts imaged with DIC microscopy using a 10X objective are shown (scale bar = 50µm). **C)** Diameters of Day 7 oocysts that were mCherry positive were measured in ImageJ with n = 100 oocysts measured per replicate. GFP expression was not taken into account for these measurements. **D)** Representative images of Day 7 midguts imaged with fluorescence microscopy for mCherry fluorophore excitation using a 10x objective are shown (scale bar = 50µm). **E)** Diameters of Day 7 oocysts that were GFP and mCherry positive were measured in ImageJ with n = 80-100 oocysts measured per replicate. Kruskal-Wallis test for multiple comparisons with Dunn’s correction was used for statistical analysis of oocyst diameters (**p<0.01; ****p<0.0001). **F)** Representative images of Day 7 midguts imaged with fluorescence microscopy for GFP fluorophore excitation using a 10x objective are shown (scale bar = 50µm).

Upon transmission to mosquitoes, parasites were no longer on pyrimethamine pressure to select for the presence of these plasmids. Fluorescence microscopy of midguts dissected seven days post-blood meal demonstrates that while all oocysts express mCherry from its integrated coding sequence in the *p230p* safe harbor locus, only 50-55% of oocysts continue to express GFP from the plasmid (Fig 4D and F, quantified in Fig S7). We first measured day seven post-blood meal oocyst diameters using mCherry fluorescence to sample the entire population of mosquito-stage parasites (Fig 4C and D). Oocysts that developed from transmitted parasites containing either S-type SSU rDNA plasmid had significantly smaller diameters than that of our control with a more pronounced decrease (9-10µm smaller) observed in those that developed from S2 SSU rDNA plasmid-containing parasites (Fig 4C). We subsequently measured the diameters of oocysts that were GFP positive, as these oocysts either still retain or recently had the plasmid. We observed similar results to the diameters measured using mCherry (Fig 4E and F). This indicates additional copies of S-type SSU rDNA affected mosquito-stage development at an earlier time point before plasmid presence was lost.

## Discussion

Mounting evidence indicates that eukaryotes use specialized ribosomes as a way to differentially regulate translation to promote specific developmental and differentiation-based outcomes. *Plasmodium* parasites have long been acknowledged to utilize specialized ribosomes based on their distinct temporal abundance patterns and sequence variations that distinguish the A- and S-type rRNAs (reviewed in (19)). Previous investigations into the S-type ribosomes in mosquito-stage parasites, when they are most abundant, concluded that S-type ribosomes are important or essential for aspects of mosquito-stage parasite development. However, these studies had technical limitations that only allowed for partial rDNA locus deletion for either the S1 or S2 rDNA (18, 29). We overcame these past technical limitations by using a DiCre recombinase system that allowed for the generation of clean deletions of the individual S1 or S2 rDNA locus, as well as both S1 and S2 types in combination to generate the first *Plasmodium* line null of an S-type ribosome. We used these transgenic lines to study S-type ribosome essentiality.

In agreement with these previous studies, we found that the S2 (historically termed the D locus) rRNA contributes to oocyst maturation, as both of our lines lacking the S2 rRNA, PyΔS2 and PyΔS1ΔS2, had statistically significant decreased oocyst diameters (Fig 1B). Furthermore, oocysts in both deletion lines had an increased frequency of oocysts that failed to fully develop and contribute to sporozoite production (Fig 1D). These phenotypes could be due to a specific, specialized function of the S2 ribosome or an overall decreased abundance of ribosomes as proposed by the Waters laboratory (18). We also observed decreased oocyst diameters upon introducing extra copies of either S-type SSU rDNA, with a larger decrease observed with introduction of extra S2 SSU rDNA (Fig 4C and E). This demonstrated that the parasite is highly sensitive to disruptions to its regulation of either S-type ribosome abundance in early mosquito-stage parasites, which results in negative impacts to optimal transmission and development. This is notable because in the previous *P. yoelii* study investigating the S-type ribosomes, only the 18S rDNA sequence was disrupted leaving the 5.8S and 28S rDNA sequences intact, which if transcribed could have impacted this tight regulation resulting in oocysts that failed to mature as was observed (29). This sensitivity to imbalances in ribosome biogenesis is also seen in ribosomopathies of humans and model systems created to study them (reviewed in (31, 32)).

However, in contrast to these previous *Plasmodium* studies, we observed that neither S-type ribosome is required for mosquito-stage parasite development, including the processes of segmentation and the production of sporozoites capable of migrating to and invading the salivary glands (Fig 1E and F, Fig S4). This result was unexpected because these are the stages with the highest abundance of S-type rRNAs. Therefore, we turned to consider why these S-type rRNAs are maintained at low levels of abundance throughout the life cycle. We hypothesized that the S-type rRNA sequence variation and potential differences in composition compared to A-type ribosomes could provide specialized functions that would warrant their continued expression in blood stages. Because previous work concluded that S-type ribosomes were either not expressed or only expressed at very low levels in these stages, little has been done to assess their role in asexual and sexual blood stages. A prior study in *P. berghei* did not identify gross phenotypes when they looked at blood-stage parasites with individual S-type deletion lines (18), and the phenotypes of the S-type deletion lines we report in this study occur at a specific time point during infection only identified through a well-controlled growth-curve assay.

Through multiple lines of investigation, we have identified that S-type ribosomes contribute different, opposing functions in blood-stage parasites despite being present at low abundance and being vastly outnumbered by A-type ribosomes. For example, PyΔS2 parasites had decreased levels of male gametogenesis (Fig 2D), while parasites with extra copies of the S2 SSU rDNA had increased male gametogenesis (Fig 3B). From these results, we conclude that the S2 ribosome functions to promote mature, transmissible male gametocytes. Contrastingly, we observed that the S1 ribosome function impacts blood-stage parasitemia, with decreased parasitemia observed in PyΔS1 (Fig 2A) and a gain in parasitemia observed when parasites were given extra copies of S1 SSU rDNA (Fig 3A). It is likely these impacts on blood-stage parasitemia are due to a change in the ratio of asexual versus sexually differentiated parasites and we therefore interpret this to mean that while the S2 ribosome is promoting sexual differentiation, the S1 ribosome is promoting asexual replication. The presence of both S1 and S2 could reflect a need to balance these activities.

Multiple lines of evidence from this study support a functional interplay between the S1 and S2 ribosomes, which indicate that they might act to compete to promote different outcomes. For instance, the phenotypes of the PyΔS1ΔS2 parasites are not as pronounced as the phenotypes of our individual S-type deletion lines. This is also observed in our transmission data, where while all S-type deletion lines had decreased intensity of mosquito infection, the most pronounced decrease was observed for the PyΔS2 parasites (Fig 1A). Compared to these parasites, the PyΔS1ΔS2 parasites had improved intensities of mosquito infection, similar to that of the PyΔS1 parasites. This indicates that the presence of the S1 ribosome in the absence of the S2 ribosome severely decreases parasite transmissibility. We can speculate this is because of a decrease in the number of mature, male gametocytes due to both the absence of the S2 ribosome and the possible functions of the S1 ribosome that promote asexual replication. In blood-stage parasites, PyΔS1 parasites had decreased blood-stage parasitemia yet maintained male gametogenesis (Fig 2A and B), and this decreased parasitemia is rescued in our PyΔS1ΔS2 parasites (Fig 2E). However, this rescue comes at the cost of decreased male gametogenesis, phenocopying that of the PyΔS2 parasites (Fig 2F). Therefore, the function of the remaining S2 ribosome in promoting male gametocyte differentiation, and potentially sexual differentiation in general, would decrease asexual replication in the absence of the S1 ribosome, resulting in the observed phenotype in the PyΔS1 parasites.

In considering these results, we returned to the question of how the presence of S-type rRNAs could benefit *Plasmodium* specifically, and the broader implications of the range of possible roles of specialized ribosomes more generally. One hypothetical model for *Plasmodium’s* evolution of S-type ribosomes invokes a drive to improve the overall fitness of the parasite during its transmission to and development in the mosquito (Fig 5). In this speculated scenario, an ancestral *Plasmodium* encoded only A-type ribosomes (modeled by PyΔS1ΔS2) that were sufficient to complete the entire life cycle (Fig 1E, Fig S4), but that enabled only suboptimal male gamete production that adversely impacted mammal-to-mosquito transmission (Fig 1A). Based on our gene deletion phenotypes, the evolution of S2 ribosomes (modeled by PyΔS1) increased male gamete production (Fig 2B) and improved oocyst maturation (Fig 1B and C) . However, these benefits would come at the cost of lowering overall asexual blood-stage parasitemia (Fig 2A), thus limiting improvements to host-to-vector transmission (Fig 1A). Therefore, the evolutionary introduction of the S1 ribosome (present day wild-type parasites) would help to balance the costs and benefits of the S2 ribosome by maintaining improved male gamete production while partially restoring the first wave of blood-stage parasite development (Fig 2A and B), giving the parasite evolutionarily improved host-to-vector transmission and initial blood-stage infection of the mammalian host (Fig 1A, Fig 5). Furthermore, the S1 rDNA is a hybrid of the S2 and A-type rDNA (27, 28), an arrangement that could have readily evolved after the introduction of the S2 rDNA.

**Figure 5:**
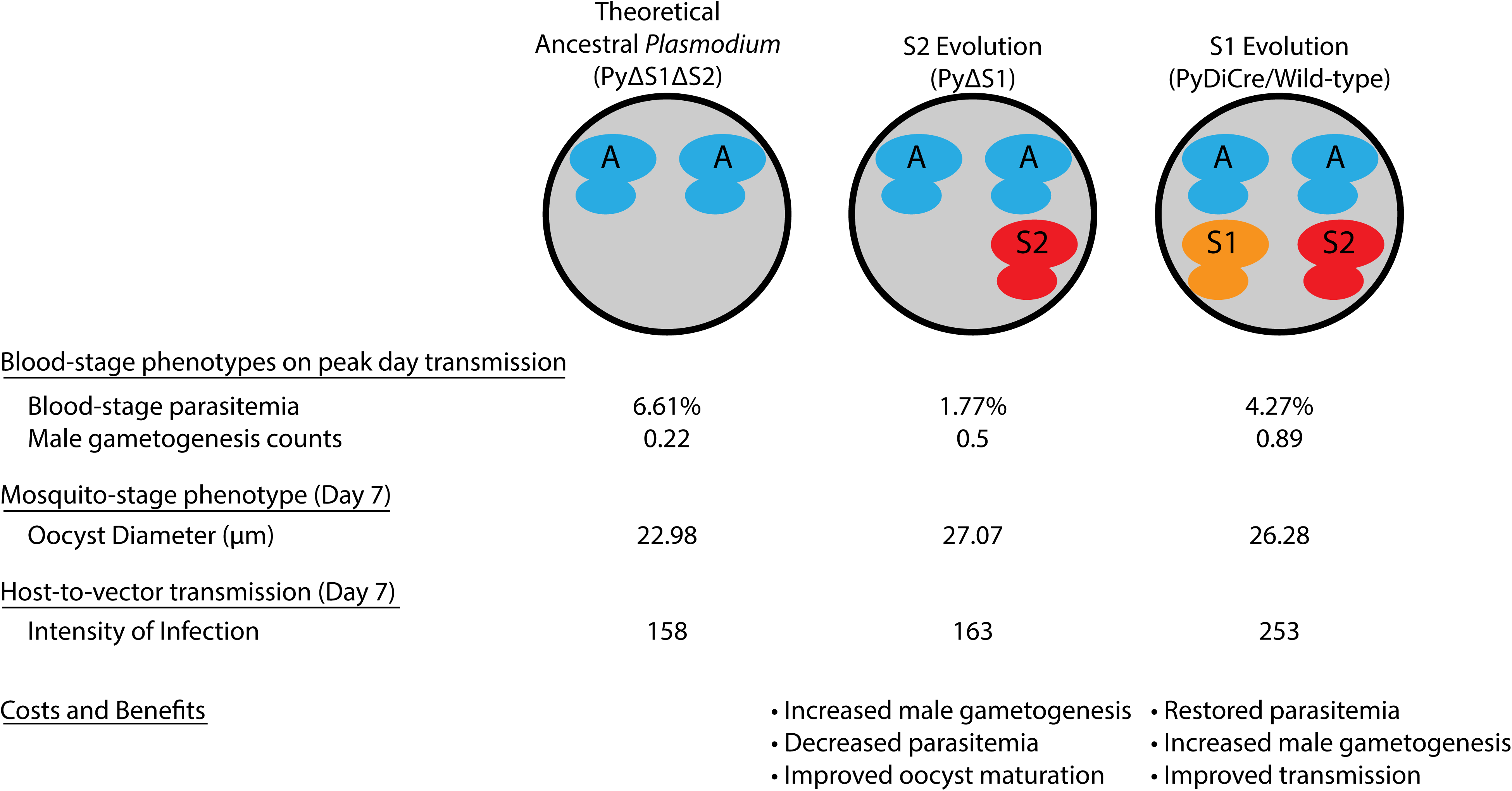
A hypothetical model of the evolutionary introduction of S-type ribosomes to *Plasmodium*. In this hypothetical scenario, an ancestral *Plasmodium* species with only A-type ribosomes (PyΔS1ΔS2) first evolved the S2 ribosomes (PyΔS1), followed by the evolution of the S1 ribosome (present day wild-type parasites). The averaged blood-stage parasitemia or male gametogenesis count (averaged centers of movement per field of view) across clonal transgenic lines or those of the control on the peak day for transmission (D5) during mouse infection are listed (Fig 2, Data S1). The averaged oocyst diameters measured from fluorescent microscopy images of infected midguts dissected 7 days post-blood meal across clonal transgenic lines or that of the control are provided (Fig 1B, Data S1). The intensity of mosquito infection averaged across clonal transgenic lines or that of the control is given (Fig 1A, Data S1). The interpreted costs and benefits of this hypothesized S-type evolution scenario are described.

Finally, it is important to consider how the S-type ribosomes could contribute specific functions and the mechanisms that enable them. The only known differences between the A, S1, and S2 ribosomes are related to their rRNA sequence variation, which exists primarily (but not exclusively) within their expansion segments (ESs). ESs are RNA protrusions from the conserved ribosomal core that have evolved in eukaryotes, vary between species (8, 19), and can have a variety of functions, most of which remain uncharacterized. For instance, specific ESs can recruit specific proteins and enzymes (and their activities) to the ribosome (33, 34) and promote interactions with mRNA (35). The model that these differences in ESs contribute differential functions is consistent with studies of the other excellent model system for studying specialized ribosomes, Zebrafish. Zebrafish have two types of specialized ribosomes expressed either in the germ line or soma, which also contain rRNA sequence variation in the ESs (13, 36). Therefore, we hypothesize the variations in S1 and S2-type ES sequences compared to each other or the A-type ESs will enable specialization of ribosomal functions via recruitment of distinct ribosomal proteins (RPs) or their paralogs, ribosome-associated proteins (RAPs), and/or enzymes that can act on the bound mRNA or the nascent peptide being translated.

Overall, our study demonstrates that the S1 and S2 ribosomes of the rodent malaria parasite *Plasmodium yoelii* are not required for mosquito-stage development. Instead, they play opposing, functional roles in blood-stage parasites to either prioritize initial population size increases or sexual differentiation to promote parasite transmission. Notably, studies on specialized ribosomes indicate that their functions occur when the specialized ribosome is the most abundant ribosome of the cell. To our knowledge, this is the first example of specialized ribosomes having functional roles in driving key cellular outcomes when they are only present at low abundances.

## Materials and Methods

### Ethics Statement

All vertebrate animal care followed the Association for Assessment and Accreditation of Laboratory Animal Care (AAALAC) guidelines and was approved by the Pennsylvania State University Institutional Animal Care and Use Committee (IACUC# PRAMS201342678). All procedures involving vertebrate animals were conducted in strict accordance with the recommendations in the *Guide for the Care and Use of Laboratory Animals* of the National Institutes of Health (37) with approved Office for Laboratory Animal Welfare (OLAW) assurance.

### Use and maintenance of experimental animals

Six-to eight-week-old female Swiss Webster (SW) mice were used for all experiments in this work. *Anopheles stephensi* mosquitoes were reared and maintained at 24°C and 70% humidity under 12-hr light/dark cycles and were fed 0.05% w/v pABA (number 100536-250G; Sigma-Aldrich) supplemented into 8% w/v, sugar water.

### Generation and confirmation of Plasmodium yoelii transgenic parasites

The PyDiCre line was generated using the same construct as for *P. berghei* (30). This construct contains a DiCre cassette, as well as a mCherry cassette under the control of an inactive truncated fragment of HSP70 promoter, and a TgDHFR/TS pyrimethamine resistance cassette flanked by two LoxP sites, to allow Cre-mediated excision and production of drug selectable marker-free parasites. A sequence corresponding to the 3’ UTR of PbDHFR was included at the end of the construct, for homologous recombination at the modified p230p locus of drug selectable marker-free GFP-expressing P. yoelii 17XNL parasites (previously described in (38)). Following transfection of PyGFP parasites, integration of the construct by double homologous recombination resulted in the replacement of GFP by mCherry and reconstitution of a functional HSP70 promoter to drive mCherry expression, along with insertion of the DiCre and TgDHFR/TS cassettes. Transfected parasites were selected with pyrimethamine and mCherry-positive parasites were sorted by FACS. The resulting parasite population was exposed to a single dose of rapamycin that was administered orally to mice. This treatment resulted in excision of the TgDHFR/TS cassette and retention of a single LoxP site. Cloning by limiting dilution resulted in the final drug selectable-marker free mCherry-expressing PyDiCre parasite line.

This PyDiCre line was used for generating *P. yoelii* transgenic parasites in this study. Ribosomal RNA deletion lines were produced (and S1 LSU rDNA sequence deletions attempted) through the conventional reverse genetics approach of using two homology regions (HRs), with one to target upstream and another to target downstream of an entire, specific rDNA locus. Homology regions were PCR amplified from Py17XNL Clone 1.1 genomic DNA (39), combined by SOE PCR, and ligated into pCR-Blunt (Invitrogen) for Sanger sequencing. Upon sequence validation, the merged 3’ and 5’ HRs were inserted into a modified pDEF plasmid containing GFP and HsDHFR expression cassettes flanked by LoxP sites as previously described (40).

PyDiCre blood-stage parasites used for transfections were cultured *ex vivo* to produce schizonts and were then purified by an Accudenz discontinuous gradient as previously described (41). These enriched schizonts were transfected in technical duplicate with 10-20µg linearized plasmid in cytomix with an Amaxa Nucleofector 2b device using Program T-016. Transgenic parasites were selected by drug cycling in a parental and a transfer mouse using 70ug/ml pyrimethamine in drinking water. Parasite populations with genotypic evidence of transgenic parasites were enriched by FACS for GFP+ positive parasites. Excision of targeted DNA was achieved via DiCre recombinase by injecting mice intraperitoneally with 4 mg/ml rapamycin (Fisher Scientific, Cat #AAJ62473MF) in 5% v/v PEG-400 (VWR, Cat # 200002-062), 5% v/v Tween-80 (VWR, Cat # VWRVM126-100ML), and 4% v/v DMSO (Fisher Scientific, Cat #PI85190) at a dosage of 4 mg rapamycin/kg mouse weight (42). Mice were injected with rapamycin when blood-stage parasitemia hit 1-2% and 48 hours later were euthanized by CO_2_ and exsanguinated by cardiac puncture for blood-stage parasite collection. Finally, transgenic parasites were cloned by limiting dilution approaches and clonality was validated by genotyping PCR.

The Small Subunit (SSU) rDNA for plasmids was PCR amplified from Py17XNL clone 1.1 genomic DNA from the specific-type promoter predicted to be ∼1000bp upstream (29) of the specific type 18S gene (S1-PY17X_0522400; S2-PY17X_0625700) through the 5.8S gene to contain Promoter-ETS-18S-ITS1-5.8S rDNA. The control plasmid, pSL0489, contains a HsDHFR and GFP expression cassette as previously described (40), and the S-type SSU rDNA amplicons were ligated into a modified SL0489. Circular plasmids were transfected into PyDiCre-purified schizonts as described above. Transgenic parasites were selected for and maintained on pyrimethamine at all times starting one day post-transfection and confirmed via genotyping PCR for plasmid presence.

All primers used for genomic DNA amplification and genotyping PCRs described are provided in Table S1. The PSU 1kb DNA ladder flanks all experimental lanes of genotyping PCRs (43). Complete plasmid sequences of plasmids described and transfected are provided in File S1.

### Host-to-vector transmission of Plasmodium yoelii

Mice were infected intraperitoneally with experimental lines and the respective control for mosquito transmission trials in biological triplicate. Infections of mice were monitored daily for parasitemia by Giemsa-stained thin blood smears and for male gametogenesis events (visible as discrete exflagellation centers) by wet mount of a drop of blood incubated at room temperature for 8 to 10 min, as described previously (23). On the peak day of exflagellation determined by the control line, the infected mice were anesthetized with a ketamine-xylazine cocktail and mosquitoes were allowed to feed *ad libitum* on the mice for 20 minutes. Mice positions were adjusted every 5 minutes to allow for even feeding.

### Plasmodium yoelii mosquito-stage phenotyping

Infected mosquito midguts were dissected 7 days after taking a blood meal, and the prevalence and intensity of infection were assessed by differential interference contrast (DIC) and live fluorescence microscopy with representative images taken at 10x and 20x objectives. These fluorescent representative images were used to measure oocyst diameters using the line tool in the ImageJ software (44) and, when applicable, the percentage of oocysts that were GFP+ and/or mCherry+. Mosquito salivary glands were dissected 14 days post-blood meal to check for sporozoite presence and to capture representative fluorescence images using a 63x objective. Of these salivary glands, 10-20 were pooled and homogenized, and sporozoites were counted using a hemocytometer. Representative images of salivary gland sporozoites were used to measure the length of sporozoites using the line segment tool in the ImageJ software (44).

### Plasmodium yoelii blood-stage phenotyping

Starter mice were infected intraperitoneally with the control or experimental parasite lines. Upon parasitemia hitting 1-3%, starter mice blood was collected by cardiac puncture and diluted in RPMI to 1,000 parasites per 100 µl. Experimental mice were infected with these 1,000 parasites intravenously, which was performed in technical triplicate for each biological replicate. Infections were monitored daily for parasitemia via Giemsa-stained thin blood smears and male gametogenesis as described above. Mouse infection with *P. yoelii* was allowed to continue until the mouse cleared the infection. DIC and fluorescent microscopy of GFP+ blood-stage parasites were performed using a 63x objective.

### Image acquisition and microscopy analyses

All DIC and fluorescence microscopy described in this study was performed using the noted objectives on the Zeiss Axioscope A1 microscope with representative images taken on the Axiocam 305 mono camera and then processed in Zen Lite version 3.8. The Leica DM750 illumination light microscope was used to count blood-stage parasitemia of Giemsa-stained thin blood smears at 100x objective (brightfield) and male exflagellation centers of wet mounts at 40x objective (phase contrast).

### Statistical analyses

All statistical tests were conducted with GraphPad Prism (v6) with the specific tests noted when used and the p-values defined (*p<0.05; **p<0.01; ***p<0.001; ****p<0.0001).

## Data Availability

All datasets with this study are in the main and supporting files associated with this manuscript.

## Acknowledgements

The authors would like to acknowledge the Huck Institutes’ Core Facility of Penn State at University Park that was instrumental in conducting this work-Genomics: RRID:SCR_023645. All of our work is greatly enabled and impacted by VEuPathDB as an essential resource. We also acknowledge members of the Llinás and Lindner laboratories for critical discussions of this work.

## Author Contributions: (CREDIT Designations)

Conceptualization: JPM, SEL

Data curation: JPM, SEL

Formal analysis: JPM, SEL

Funding acquisition: SEL

Investigation: JPM, SEL

Methodology: JPM, SEL

Project administration: JPM, SEL

Resources: JPM, SB, OS, SEL

Software: N/A

Supervision: SEL

Validation: JPM

Visualization: JPM, SEL

Writing-original draft: JPM, SEL

Writing-reviewing and editing: JPM, SB, OS, SEL

## References

1. Henras AK, Plisson-Chastang C, O’Donohue MF, Chakraborty A, Gleizes PE. An overview of pre-ribosomal RNA processing in eukaryotes. Wiley Interdiscip Rev RNA. 2015;6(2):225–42.

2. Potapova TA, Gerton JL. Ribosomal DNA and the nucleolus in the context of genome organization. Chromosome Res. 2019;27(1-2):109–27.

3. Weijers D, Franke-van Dijk M, Vencken RJ, Quint A, Hooykaas P, Offringa R. An Arabidopsis Minute-like phenotype caused by a semi-dominant mutation in a RIBOSOMAL PROTEIN S5 gene. Development. 2001;128(21):4289–99.

4. Hopes T, Norris K, Agapiou M, McCarthy CGP, Lewis PA, O’Connell MJ, et al. Ribosome heterogeneity in Drosophila melanogaster gonads through paralog-switching. Nucleic Acids Res. 2022;50(4):2240–57.

5. Sugihara Y, Honda H, Iida T, Morinaga T, Hino S, Okajima T, et al. Proteomic analysis of rodent ribosomes revealed heterogeneity including ribosomal proteins L10-like, L22-like 1, and L39-like. J Proteome Res. 2010;9(3):1351–66.

6. Ramagopal S, Ennis HL. Regulation of synthesis of cell-specific ribosomal proteins during differentiation of Dictyostelium discoideum. Proc Natl Acad Sci U S A. 1981;78(5):3083–7.

7. Siekevitz P, Palade GE. A cytochemical study on the pancreas of the guinea pig. II. Functional variations in the enzymatic activity of microsomes. J Biophys Biochem Cytol. 1958;4(3):309–18.

8. Gerbi SA. The evolution of eukaryotic ribosomal DNA. Biosystems. 1986;19(4):247–58.

9. Fan W, Eklund E, Sherman RM, Liu H, Pitts S, Ford B, et al. Widespread genetic heterogeneity of human ribosomal RNA genes. RNA. 2022;28(4):478–92.

10. Genuth NR, Barna M. Heterogeneity and specialized functions of translation machinery: from genes to organisms. Nat Rev Genet. 2018;19(7):431–52.

11. Li H, Huo Y, He X, Yao L, Zhang H, Cui Y, et al. A male germ-cell-specific ribosome controls male fertility. Nature. 2022;612(7941):725-31.

12. Genuth NR, Shi Z, Kunimoto K, Hung V, Xu AF, Kerr CH, et al. A stem cell roadmap of ribosome heterogeneity reveals a function for RPL10A in mesoderm production. Nat Commun. 2022;13(1):5491.

13. Locati MD, Pagano JFB, Girard G, Ensink WA, van Olst M, van Leeuwen S, et al. Expression of distinct maternal and somatic 5.8S, 18S, and 28S rRNA types during zebrafish development. RNA. 2017;23(8):1188-99.

14. Gardner MJ, Hall N, Fung E, White O, Berriman M, Hyman RW, et al. Genome sequence of the human malaria parasite Plasmodium falciparum. Nature. 2002;419(6906):498-511.

15. Gunderson JH, Sogin ML, Wollett G, Hollingdale M, de la Cruz VF, Waters AP, et al. Structurally distinct, stage-specific ribosomes occur in Plasmodium. Science. 1987;238(4829):933-7.

16. Li J, McConkey GA, Rogers MJ, Waters AP, McCutchan TR. Plasmodium: the developmentally regulated ribosome. Exp Parasitol. 1994;78(4):437–41.

17. Rogers MJ, Gutell RR, Damberger SH, Li J, McConkey GA, Waters AP, et al. Structural features of the large subunit rRNA expressed in Plasmodium falciparum sporozoites that distinguish it from the asexually expressed subunit rRNA. RNA. 1996;2(2):134–45.

18. van Spaendonk RM, Ramesar J, van Wigcheren A, Eling W, Beetsma AL, van Gemert GJ, et al. Functional equivalence of structurally distinct ribosomes in the malaria parasite, Plasmodium berghei. J Biol Chem. 2001;276(25):22638–47.

19. McGee JP, Armache JP, Lindner SE. Ribosome heterogeneity and specialization of Plasmodium parasites. PLoS Pathog. 2023;19(4):e1011267.

20. Fang J, Sullivan M, McCutchan TF. The effects of glucose concentration on the reciprocal regulation of rRNA promoters in Plasmodium falciparum. J Biol Chem. 2004;279(1):720–5.

21. Fang J, McCutchan TF. Corrigendum: Malaria: Thermoregulation in a parasite’s life cycle. Nature. 2016;537(7619):254.

22. Lindner SE, Swearingen KE, Shears MJ, Walker MP, Vrana EN, Hart KJ, et al. Transcriptomics and proteomics reveal two waves of translational repression during the maturation of malaria parasite sporozoites. Nature Communications. 2019;10(1).

23. Hart KJ, Oberstaller J, Walker MP, Minns AM, Kennedy MF, Padykula I, et al. Plasmodium male gametocyte development and transmission are critically regulated by the two putative deadenylases of the CAF1/CCR4/NOT complex. PLoS Pathog. 2019;15(1):e1007164.

24. Howick VM, Russell AJC, Andrews T, Heaton H, Reid AJ, Natarajan K, et al. The Malaria Cell Atlas: Single parasite transcriptomes across the complete Plasmodium life cycle. Science. 2019;365(6455).

25. Reid AJ, Talman AM, Bennett HM, Gomes AR, Sanders MJ, Illingworth CJR, et al. Single-cell RNA-seq reveals hidden transcriptional variation in malaria parasites. Elife. 2018;7.

26. Hart KJ, Power BJ, Rios KT, Sebastian A, Lindner SE. The Plasmodium NOT1-G paralogue is an essential regulator of sexual stage maturation and parasite transmission. PLoS Biol. 2021;19(10):e3001434.

27. van Spaendonk RM, Ramesar J, Janse CJ, Waters AP. The rodent malaria parasite Plasmodium berghei does not contain a typical O-type small subunit ribosomal RNA gene. Mol Biochem Parasitol. 2000;105(1):169–74.

28. Amos B, Aurrecoechea C, Barba M, Barreto A, Basenko EY, Bazant W, et al. VEuPathDB: the eukaryotic pathogen, vector and host bioinformatics resource center. Nucleic Acids Res. 2022;50(D1):D898–D911.

29. Qi Y, Zhu F, Eastman RT, Fu Y, Zilversmit M, Pattaradilokrat S, et al. Regulation of Plasmodium yoelii oocyst development by strain- and stage-specific small-subunit rRNA. mBio. 2015;6(2):e00117.

30. Fernandes P, Briquet S, Patarot D, Loubens M, Hoareau-Coudert B, Silvie O. The dimerisable Cre recombinase allows conditional genome editing in the mosquito stages of Plasmodium berghei. PLoS One. 2020;15(10):e0236616.

31. Narla A, Ebert BL. Ribosomopathies: human disorders of ribosome dysfunction. Blood. 2010;115(16):3196–205.

32. Mills EW, Green R. Ribosomopathies: There’s strength in numbers. Science. 2017;358(6363).

33. Fujii K, Susanto TT, Saurabh S, Barna M. Decoding the Function of Expansion Segments in Ribosomes. Mol Cell. 2018;72(6):1013–20 e6.

34. Krauer N, Rauscher R, Polacek N. tRNA Synthetases Are Recruited to Yeast Ribosomes by rRNA Expansion Segment 7L but Do Not Require Association for Functionality. Noncoding RNA. 2021;7(4).

35. Diaz-Lopez I, Toribio R, Berlanga JJ, Ventoso I. An mRNA-binding channel in the ES6S region of the translation 48S-PIC promotes RNA unwinding and scanning. Elife. 2019;8.

36. Shah AN, Leesch F, Lorenzo-Orts L, Grundmann L, Novatchkova M, Haselbach D, et al. A dual ribosomal system in the zebrafish soma and germline. bioRxiv. 2024.

37. Guide for the care and use of laboratory animals. 8th ed. Washington, DC: National Academies Press; 2011.

38. Manzoni G, Briquet S, Risco-Castillo V, Gaultier C, Topcu S, Ivanescu ML, et al. A rapid and robust selection procedure for generating drug-selectable marker-free recombinant malaria parasites. Sci Rep. 2014;4:4760.

39. Godin MJ, Sebastian A, Albert I, Lindner SE. Long-read genome assembly and gene model annotations for the rodent malaria parasite Plasmodium yoelii 17XNL. J Biol Chem. 2023;299(7):104871.

40. Munoz EE, Hart KJ, Walker MP, Kennedy MF, Shipley MM, Lindner SE. ALBA4 modulates its stage-specific interactions and specific mRNA fates during Plasmodium yoelii growth and transmission. Mol Microbiol. 2017;106(2):266–84.

41. Lindner SE, Mikolajczak SA, Vaughan AM, Moon W, Joyce BR, Sullivan WJ, Jr., et al. Perturbations of Plasmodium Puf2 expression and RNA-seq of Puf2-deficient sporozoites reveal a critical role in maintaining RNA homeostasis and parasite transmissibility. Cell Microbiol. 2013;15(7):1266–83.

42. Bitto A, Ito TK, Pineda VV, LeTexier NJ, Huang HZ, Sutlief E, et al. Transient rapamycin treatment can increase lifespan and healthspan in middle-aged mice. Elife. 2016;5.

43. Henrici RC, Pecen TJ, Johnston JL, Tan S. The pPSU Plasmids for Generating DNA Molecular Weight Markers. Sci Rep. 2017;7(1):2438.

44. Schneider CA, Rasband WS, Eliceiri KW. NIH Image to ImageJ: 25 years of image analysis. Nat Methods. 2012;9(7):671–5.

