## Supplementary material for "Specialized S-type ribosomes of *Plasmodium yoelii* enhance host-to-vector malaria transmission": File S1

McGee *et al*. Supplemental File 1- Complete Plasmid Sequences

**pSL0489**

LOCUS	pSL0489__PSd_	7756 bp	DNA	circular	27-JUN-2024

SOURCE

ORGANISM

COMMENT

COMMENT

COMMENT	ApEinfo:methylated:1

FEATURES	Location/Qualifiers

exon	4088..4945

/vntifkey="61"

/locus_tag="AMP"

/label="AMP"

/ApEinfo_label="AMP"

/ApEinfo_fwdcolor="pink"

/ApEinfo_revcolor="pink"

/ApEinfo_graphicformat="arrow_data {{0 1 2 0 0 -1} {} 0}

width 5 offset 0"

rep_origin	5043..5725

/locus_tag="ColE1 origin"

/label="ColE1 origin"

/ApEinfo_label="ColE1 origin"

/ApEinfo_fwdcolor="gray50"

/ApEinfo_revcolor="gray50"

/ApEinfo_graphicformat="arrow_data {{0 1 2 0 0 -1} {} 0}

width 5 offset 0"

exon	6733..7296

/vntifkey="61"

/locus_tag="hDHFR"

/label="hDHFR"

/ApEinfo_label="hDHFR"

/ApEinfo_fwdcolor="pink"

/ApEinfo_revcolor="pink"

/ApEinfo_graphicformat="arrow_data {{0 1 2 0 0 -1} {} 0}

width 5 offset 0"

misc_feature	join(6129..6438,6443..6732)

/vntifkey="21"

/locus_tag="PbEF1a-A\5'UTR"

/label="PbEF1a-A\5'UTR"

/ApEinfo_label="PbEF1a-A\5'UTR"

/ApEinfo_fwdcolor="#eb0214"

/ApEinfo_revcolor="#eb0214"

/ApEinfo_graphicformat="arrow_data {{0 1 2 0 0 -1} {} 0}

width 5 offset 0"

misc_feature	1897..2470

/locus_tag="PbEF1a promoter (from pSL0281)"

/label="PbEF1a promoter (from pSL0281)"

/ApEinfo_label="PbEF1a promoter (from pSL0281)"

/ApEinfo_fwdcolor="cyan"

/ApEinfo_revcolor="green"

/ApEinfo_graphicformat="arrow_data {{0 1 2 0 0 -1} {} 0}

width 5 offset 0"

misc_feature	2515..3193

/locus_tag="GFPmut2"

/label="GFPmut2"

/ApEinfo_label="GFPmut2"

/ApEinfo_fwdcolor="cyan"

/ApEinfo_revcolor="green"

/ApEinfo_graphicformat="arrow_data {{0 1 2 0 0 -1} {} 0}

width 5 offset 0"

misc_feature	7297..7753

/vntifkey="21"

/locus_tag="PbDHFR/TS\3'UTR"

/label="PbDHFR/TS\3'UTR"

/ApEinfo_label="PbDHFR/TS\3'UTR"

/ApEinfo_fwdcolor="#eb0214"

/ApEinfo_revcolor="#eb0214"

/ApEinfo_graphicformat="arrow_data {{0 1 2 0 0 -1} {} 0}

width 5 offset 0"

misc_feature	3200..3646

/locus_tag="PbDHFR/TS\3'UTR(1)"

/label="PbDHFR/TS\3'UTR(1)"

/ApEinfo_label="PbDHFR/TS\3'UTR"

/ApEinfo_fwdcolor="#ff0000"

/ApEinfo_revcolor="green"

/ApEinfo_graphicformat="arrow_data {{0 1 2 0 0 -1} {} 0}

width 5 offset 0"

CDS	4286..4945

/locus_tag="AmpR"

/label="AmpR"

/ApEinfo_label="AmpR"

/ApEinfo_fwdcolor="yellow"

/ApEinfo_revcolor="yellow"

/ApEinfo_graphicformat="arrow_data {{0 1 2 0 0 -1} {} 0}

width 5 offset 0"

misc_feature	7303..7753

/locus_tag="PbDHFR-TS 3'UTR"

/label="PbDHFR-TS 3'UTR"

/ApEinfo_label="PbDHFR-TS 3'UTR"

/ApEinfo_fwdcolor="cyan"

/ApEinfo_revcolor="green"

/ApEinfo_graphicformat="arrow_data {{0 1 2 0 0 -1} {} 0}

width 5 offset 0"

misc_feature	join(6129..6438,6443..6727)

/locus_tag="PbEF1a 5'UTR"

/label="PbEF1a 5'UTR"

/ApEinfo_label="PbEF1a 5'UTR"

/ApEinfo_fwdcolor="#804040"

/ApEinfo_revcolor="#804040"

/ApEinfo_graphicformat="arrow_data {{0 1 2 0 0 -1} {} 0}

width 5 offset 0"

misc_feature	7304..7751

/locus_tag="PbDHFR-TS 3'UTR(1)"

/label="PbDHFR-TS 3'UTR(1)"

/ApEinfo_label="PbDHFR-TS 3'UTR"

/ApEinfo_fwdcolor="cyan"

/ApEinfo_revcolor="green"

/ApEinfo_graphicformat="arrow_data {{0 1 2 0 0 -1} {} 0}

width 5 offset 0"

misc_feature	7..1872

/locus_tag="Py p230p targeting sequences"

/label="Py p230p targeting sequences"

/ApEinfo_label="Py p230p targeting sequences"

/ApEinfo_fwdcolor="cyan"

/ApEinfo_revcolor="green"

/ApEinfo_graphicformat="arrow_data {{0 1 2 0 0 -1} {} 0}

width 5 offset 0"

ORIGIN

1 ggtaccAGCT TTGTGTTTTA TTTGGATGTG CAACAATAGA TTTATGATTA ACCAATGGTA

61 AATCATTATA ATCATATATT TCATCTTCAT TTACATATTT TACTAAAACA GTTTTTACAA

121 TTTGTTCTGT TTCTTTATCT ACACAATGAC ATTCAAATTG ATATGATTTT GTTAATGTAT

181 AATTAAAACC AGCTAATGCA TATTCTGGAT TTTCAAATGA ATAGTTAATA ATATTTTTAG

241 TATTAATTTT ATAGGAATCA TGTAGATAGC CTTTTTCATA TACTTGAGTA AAACATTTTT

301 CTGGCATTTT TTTTGTATTT TCGGGGCATC TAAAACCAAA ACTTGTATTT GATTTGGAAA

361 TTATTATACA TAGATCTCCT TCACTATAAG CATTTAAATA ATTATATAAT TCACCTTTTC

421 TAAAATCACA AAATGAAACT GGTCTATCTT TTAAATCTGT TCGAACATGT AATAAAATAA

481 TTGCTCTTCT ATAATCAGTC CCACCATTTG TTACACCACA TTCTATGGTT AGTGGATATG

541 ATCCATGATT TATAAATGAG GTCATAAAAA ATTTAAAACC TTTAAATTGT TCATTTATTT

601 CTGGTATATC TATATCTTTT GTATTAAAAA ATAAATCTGT AAATTTATTT ATAGGAAATA

661 CATTACTTAA ATATTTTATG GTATTAATAG GTTCTTTACC ATATACGATT CTATTATCTG

721 TTATTTGCTC ACAATTATAC ATGAATAAAC TAAATTCATT TATTGTTTTT CACATATTTG

781 TATTTCACTT AATTTGCTTA AATTATAATC AAAATTACAC ATCTGATAAG GTATATCGTT

841 AAAATATTTA ATCGTTTTAC TATCATCATC AACTATTTTT AAAATGTGAT CAGATGTTGC

901 CATTCCATCA CCAAATGggg cccTATGGAA CTACATCTAT ATAAGAGATT TTTTTATTTT

961 TATATATTGG TTTTGAATCT AGATATTCAT AATTTTTAAT ATATATTGTT CCTATATTTT

1021 TGGTTTTATA ATCTTCACAT GTACAATAAA AATCAAATTC TTTTTTTGTA ATTTTTGAAA

1081 TCATTAAAAA TGTTGATGAA TTATTTATAA AATAATCTGC ATCTATAACT TTTATTTCAT

1141 TAAATATATT TTCGATATTT TCAGTTATTT GTTCATTCAT GTTACTTTTT ATATATACCT

1201 TTCTAAAACA ATTTTTTGGA TTAATTTTTC CAGTTGGACA ATTTAAGCTA ACTACTATAT

1261 TTTCTTTTAA TTCGATTTCA CAAAATTCAT TTTGTTTTTC TAAATCATAA ACATTGGAAT

1321 AGTAAAATTT TTTTTTAGAA TTACCTGAAA AATCACATCC ATATAATTTA TTTTTTGTTG

1381 TATTTTGAGA AACCCTTATC ACTGTAATAC CTTTTTGACC TTTCATATTA TTAATTATTG

1441 TTTTAGTATT ATCACAATAT ATTTTCAATA TTAATACTTT TTCTACAATT AATGGAGTAA

1501 CAAAAGTGAA ATTTCCTTTC TCTTTGAACA TTTTATTTTT ATCTAATATG ATAACACCTG

1561 GTAATATATC ATCTAATTTT TCTTTTTTGT CTAGATACAT TACTTCATCA TTTTCTGGTA

1621 TGTATTCTAT ATTTTTATAT TTTTCTAATT CTTCTGGTTC AACATTTGCA TTTTCTAAAT

1681 AAAAATTATT TTTTTGACTA CTACGTTGTG ATGTTTTCTT ATTATTTATT ATGATATTAC

1741 TATCAAAAAT GTCATCATCA CTTTTTACAA TTTTGGGACA TTTAACTTGA ACCATATTCA

1801 GATTCTGTTC TAGATGTATA TAACATGAAT GAACTGTATC TACTTCATTA ACGGGTTCAA

1861 GAGAATTCGg tagccgcggt ggcggccgcT ctcgagAGCT TAATTCTTTT CGAGCTCTTT

1921 ATGCTTAAGT TTACAATTTA ATATTCATAC TTTAAGTATT TTTTGTAGTA TCCTAGATAT

1981 TGTGCTTTAA ATGCTCACCC CTCAAAGCAC CAGTAATATT TTCATCCACT GAAATACCAT

2041 TAAATTTTCA AAAAAATACT ATGCATATAA TGTTATACAT ATAAACATAA AACGCCATGT

2101 AAATCAAAAA ATATATAAAA ATATGTATAA AAATAAATAT GCACTAAATA TAAGCTAATT

2161 ATGCATAAAA ATTAAAGTGC CCTTTATTAA CTAGctagTC GTAATTATTT ATATTTCTAT

2221 GTTATAAAAA AATCCTCATA TAATAATATA ATTAATATAT GTAATGTTTT TTTTATTTTA

2281 TAATTTTAAT ATAAAATAAT ATGTAAATTA ATTCAAAAAA TAAATATAAT TGTTGTGAAA

2341 CAAAAAACGT AATTTTTTCA TTTGCCTTCA AAATTTAAAT TTATTTTAAT ATTTCCTAAA

2401 ATATATATAC TTTGTGTATA AATATATAAA AATATATATT TGCTTATAAA TAAATAAAAA

2461 TTTTATAAAA atgactagta gtaaaggaga agaacttttc actggagttg tcccAATTCT

2521 TGTTGAATTA GATGGTGATG TTAATGGGCA CAAATTTTCT GTCAGTGGAG AGGGTGAAGG

2581 TGATGCAACA TACGGAAAAC TTACCCTTAA ATTTATTTGC ACTACTGGAA AACTACCTGT

2641 TCCATGGCCA ACACTTGTCA CTACTTTCGC GTATGGTCTT CAATGCTTTG CGAGATACCC

2701 AGATCATATG AAACAGCATG ACTTTTTCAA GAGTGCCATG CCCGAAGGTT ATGTACAGGA

2761 AAGAACTATA TTTTTCAAAG ATGACGGGAA CTACAAGACA CGTGCTGAAG TCAAGTTTGA

2821 AGGTGATACC CTTGTTAATA GAATCGAGTT AAAAGGTATT GATTTTAAAG AAGATGGAAA

2881 CATTCTTGGA CACAAATTGG AATACAACTA TAACTCACAC AATGTATACA TCATGGCAGA

2941 CAAACAAAAG AATGGAATCA AAGTTAACTT CAAAATTAGA CACAACATTG AAGATGGAAG

3001 CGTTCAACTA GCAGACCATT ATCAACAAAA TACTCCAATT GGCGATGGCC CTGTCCTTTT

3061 ACCAGACAAC CATTACCTGT CCACACAATC TGCCCTTTCG AAAGATCCCA ACGAAAAGAG

3121 AGACCACATG GTCCTTCTTG AGTTTGTAAC AGCTGCTGGG ATTACACATG GCATGGATGA

3181 ACTATACAAA TAAggatccG TTTTTCTTAC TTATATATTT ATACCAATTG ATTGTATTTA

3241 TAACTGTAAA AATGTGTATG TTGTGTGCAT ATTTTTTTTT GTGCATGCAC ATGCATGTAA

3301 ATAGCTAAAA TTATGAACAT TTTATTTTTT GTTCAGAAAA AAAAACTTTA CACACATAAA

3361 ATGGCTAGTA TGAATAGCCA TATTTTATAT AAATTAAATC CTATGAATTT ATGACCATAT

3421 TAAAAATTTA GATATTTATG GAACATAATA TGTTTGAAAC AATAAGACAA AATTATTATT

3481 ATTATTATTA TTTTTACTGT TATAATTATG TTGTCTCTTC AATGATTCAT AAATAGTTGG

3541 ACTTGATTTT TAAAATGTTT ATAATATGAT TAGCATAGTT AAATAAAAAA AGTTGAAAAA

3601 TTAAAAAAAA ACATATAAAC ACAAATGATG TTTTTTCCTT CAATTTcggc gcctgatgcg

3661 gtattttctc cttacgcatc tgtgcggtat ttcacaccgc atatggtgca ctctcagtac

3721 aatctgctct gatgccgcat agttaagcca gccccgacac ccgccaacac ccgctgacgc

3781 gccctgacgg gcttgtctgc tcccggcatc cgcttacaga caagctgtga ccgtctccgg

3841 gagctgcatg tgtcagaggt tttcaccgtc atcaccgaaa cgcgcgagac gaaagggcct

3901 cgtgatacgc ctatttttat aggttaatgt catgataata atggtttctt agacgtcagg

3961 tggcactttt cggggaaatg tgcgcggaac ccctatttgt ttatttttct aaatacattc

4021 aaatatgtat ccgctcatga gacaataacc ctgataaatg cttcaataat attgaaaaag

4081 gaagagtatg agtattcaac atttccgtgt cgcccttatt cccttttttg cggcattttg

4141 ccttcctgtt tttgctcacc cagaaacgct ggtgaaagta aaagatgctg aagatcagtt

4201 gggtgcacga gtgggttaca tcgaactgga tctcaacagc ggtaagatcc ttgagagttt

4261 tcgccccgaa gaacgttttc caatgatgag cacttttaaa gttctgctat gtggcgcggt

4321 attatcccgt attgacgccg ggcaagagca actcggtcgc cgcatacact attctcagaa

4381 tgacttggtt gagtactcac cagtcacaga aaagcatctt acggatggca tgacagtaag

4441 agaattatgc agtgctgcca taaccatgag tgataacact gcggccaact tacttctgac

4501 aacgatcgga ggaccgaagg agctaaccgc ttttttgcac aacatggggg atcatgtaac

4561 tcgccttgat cgttgggaac cggagctgaa tgaagccata ccaaacgacg agcgtgacac

4621 cacgatgcct gtagcaatgg caacaacgtt gcgcaaacta ttaactggcg aactacttac

4681 tctagcttcc cggcaacaat taatagactg gatggaggcg gataaagttg caggaccact

4741 tctgcgctcg gcccttccgg ctggctggtt tattgctgat aaatctggag ccggtgagcg

4801 tgggtctcgc ggtatcattg cagcactggg gccagatggt aagccctccc gtatcgtagt

4861 tatctacacg acggggagtc aggcaactat ggatgaacga aatagacaga tcgctgagat

4921 aggtgcctca ctgattaagc attggtaact gtcagaccaa gtttactcat atatacttta

4981 gattgattta aaacttcatt tttaatttaa aaggatctag gtgaagatcc tttttgataa

5041 tctcatgacc aaaatccctt aacgtgagtt ttcgttccac tgagcgtcag accccgtaga

5101 aaagatcaaa ggatcttctt gagatccttt ttttctgcgc gtaatctgct gcttgcaaac

5161 aaaaaaacca ccgctaccag cggtggtttg tttgccggat caagagctac caactctttt

5221 tccgaaggta actggcttca gcagagcgca gataccaaat actgttcttc tagtgtagcc

5281 gtagttaggc caccacttca agaactctgt agcaccgcct acatacctcg ctctgctaat

5341 cctgttacca gtggctgctg ccagtggcga taagtcgtgt cttaccgggt tggactcaag

5401 acgatagtta ccggataagg cgcagcggtc gggctgaacg gggggttcgt gcacacagcc

5461 cagcttggag cgaacgacct acaccgaact gagataccta cagcgtgagc tatgagaaag

5521 cgccacgctt cccgaaggga gaaaggcgga caggtatccg gtaagcggca gggtcggaac

5581 aggagagcgc acgagggagc ttccaggggg aaacgcctgg tatctttata gtcctgtcgg

5641 gtttcgccac ctctgacttg agcgtcgatt tttgtgatgc tcgtcagggg ggcggagcct

5701 atggaaaaac gccagcaacg cggccttttt acggttcctg gccttttgct ggccttttgc

5761 tcacatgttc tttcctgcgt tatcccctga ttctgtggat aaccgtatta ccgcctttga

5821 gtgagctgat accgctcgcc gcagccgaac gaccgagcgc agcgagtcag tgagcgagga

5881 agcggaagag cgcccaatac gcaaaccgcc tctccccgcg cgttggccga ttcattaatg

5941 cagctggcac gacaggtttc ccgactggaa agcgggcagt gagcgcaacg caattaatgt

6001 gagttagctc actcattagg caccccaggc tttacacttt atgcttccgg ctcgtatgtt

6061 gtgtggaatt gtgagcggat aacaatttca cacaggaaac agctatgacc atgattacgc

6121 caagcttgat aattcctgca gcccagctta attcttttcg agctctttat gcttaagttt

6181 acaatttaat attcatactt taagtatttt ttgtagtatc ctagatattg tgctttaaat

6241 gctcacccct caaagcacca gtaatatttt catccactga aataccatta aattttcaaa

6301 aaaatactat gcatataatg ttatacatat aaacataaaa cgccatgtaa atcaaaaaat

6361 atataaaaat atgtataaaa ataaatatgc actaaatata agctaattat gcataaaaat

6421 taaagtgccc tttattaact agctagtcgt aattatttat atttctatgt tataaaaaaa

6481 tcctcatata ataatataat taatatatgt aatgtttttt ttattttata attttaatat

6541 aaaataatat gtaaattaat tcaaaaaata aatataattg ttgtgaaaca aaaaacgtaa

6601 ttttttcatt tgccttcaaa atttaaattt attttaatat ttcctaaaat atatatactt

6661 tgtgtataaa tatataaaaa tatatatttg cttataaata aataaaaaat tttataaaac

6721 atagggggat ccatggttgg ttcgctaaac tgcatcgtcg ctgtgtccca gaacatgggc

6781 atcggcaaga acggggacct gccctggcca ccgctcagga acgaatttag atatttccag

6841 agaatgacca caacctcttc agtagaaggt aaacagaatc tggtgattat gggtaagaag

6901 acctggttct ccattcctga gaagaatcga cctttaaagg gtagaattaa tttagttctc

6961 agcagagaac tcaaggaacc tccacaagga gctcattttc tttccagaag tctagatgat

7021 gccttaaaac ttactgaaca accagaatta gcaaataaag tagacatggt ctggatagtt

7081 ggtggcagtt ctgtttataa ggaagccatg aatcacccag gccatcttaa actatttgtg

7141 acaaggatca tgcaagactt tgaaagtgac acgttttttc cagaaattga tttggagaaa

7201 tataaacttc tgccagaata cccaggtgtt ctctctgatg tccaggagga gaaaggcatt

7261 aagtacaaat ttgaagtata tgagaagaat gattaaggat cccgtttttc ttacttatat

7321 atttatacca attgattgta tttataactg taaaaatgtg tatgttgtgt gcatattttt

7381 ttttgtgcat gcacatgcat gtaaatagct aaaattatga acattttatt ttttgttcag

7441 aaaaaaaaaa ctttacacac ataaaatggc tagtatgaat agccatattt tatataaatt

7501 aaatcctatg aatttatgac catattaaaa atttagatat ttatggaaca taatatgttt

7561 gaaacaataa gacaaaatta ttattattat tattattttt actgttataa ttatgttgtc

7621 tcttcaatga ttcataaata gttggacttg atttttaaaa tgtttataat atgattagca

7681 tagttaaata aaaaaagttg aaaaattaaa aaaaaacata taaacacaaa tgatgttttt

7741 tccttcaatt tcgatg

//

**pSL1555**

LOCUS	pSL1555	7099 bp	DNA	linear	21-OCT-2024

DEFINITION .

ACCESSION

VERSION

SOURCE	.

ORGANISM .

COMMENT

COMMENT

COMMENT	ApEinfo:methylated:1

FEATURES	Location/Qualifiers

exon	6076..6353

/vntifkey="61"

/locus_tag="hDHFR"

/label="hDHFR"

/ApEinfo_label="hDHFR"

/ApEinfo_fwdcolor="pink"

/ApEinfo_revcolor="pink"

/ApEinfo_graphicformat="arrow_data {{0 1 2 0 0 -1} {} 0}

width 5 offset 0"

exon	6354..6639

/vntifkey="61"

/locus_tag="hDHFR(1)"

/label="hDHFR(1)"

/ApEinfo_label="hDHFR"

/ApEinfo_fwdcolor="pink"

/ApEinfo_revcolor="pink"

/ApEinfo_graphicformat="arrow_data {{0 1 2 0 0 -1} {} 0}

width 5 offset 0"

misc_feature	join(5472..5781,5786..6075)

/vntifkey="21"

/locus_tag="PbEF1a-A\5'UTR"

/label="PbEF1a-A\5'UTR"

/ApEinfo_label="PbEF1a-A\5'UTR"

/ApEinfo_fwdcolor="#eb0214"

/ApEinfo_revcolor="#eb0214"

/ApEinfo_graphicformat="arrow_data {{0 1 2 0 0 -1} {} 0}

width 5 offset 0"

misc_feature	1239..1812

/locus_tag="PbEF1a promoter (from pSL0281)"

/label="PbEF1a promoter (from pSL0281)"

/ApEinfo_label="PbEF1a promoter (from pSL0281)"

/ApEinfo_fwdcolor="cyan"

/ApEinfo_revcolor="green"

/ApEinfo_graphicformat="arrow_data {{0 1 2 0 0 -1} {} 0}

width 5 offset 0"

misc_feature	6646..7096

/locus_tag="PbDHFR-TS 3'UTR"

/label="PbDHFR-TS 3'UTR"

/ApEinfo_label="PbDHFR-TS 3'UTR"

/ApEinfo_fwdcolor="cyan"

/ApEinfo_revcolor="green"

/ApEinfo_graphicformat="arrow_data {{0 1 2 0 0 -1} {} 0}

width 5 offset 0"

misc_feature	634..1192

/locus_tag="5' HR for KO"

/label="5' HR for KO"

/ApEinfo_label="5' HR for KO"

/ApEinfo_fwdcolor="#ff0000"

/ApEinfo_revcolor="green"

/ApEinfo_graphicformat="arrow_data {{0 1 2 0 0 -1} {} 0}

width 5 offset 0"

misc_feature	1857..2535

/locus_tag="GFPmut2"

/label="GFPmut2"

/ApEinfo_label="GFPmut2"

/ApEinfo_fwdcolor="cyan"

/ApEinfo_revcolor="green"

/ApEinfo_graphicformat="arrow_data {{0 1 2 0 0 -1} {} 0}

width 5 offset 0"

misc_feature	6647..7094

/locus_tag="PbDHFR-TS 3'UTR(1)"

/label="PbDHFR-TS 3'UTR(1)"

/ApEinfo_label="PbDHFR-TS 3'UTR"

/ApEinfo_fwdcolor="cyan"

/ApEinfo_revcolor="green"

/ApEinfo_graphicformat="arrow_data {{0 1 2 0 0 -1} {} 0}

width 5 offset 0"

misc_feature	2542..2989

/locus_tag="PbDHFR/TS\3'UTR(1)"

/label="PbDHFR/TS\3'UTR(1)"

/ApEinfo_label="PbDHFR/TS\3'UTR"

/ApEinfo_fwdcolor="#ff0000"

/ApEinfo_revcolor="green"

/ApEinfo_graphicformat="arrow_data {{0 1 2 0 0 -1} {} 0}

width 5 offset 0"

rep_origin	4386..5068

/locus_tag="ColE1 origin"

/label="ColE1 origin"

/ApEinfo_label="ColE1 origin"

/ApEinfo_fwdcolor="gray50"

/ApEinfo_revcolor="gray50"

/ApEinfo_graphicformat="arrow_data {{0 1 2 0 0 -1} {} 0}

width 5 offset 0"

CDS	3629..4288

/locus_tag="AmpR"

/label="AmpR"

/ApEinfo_label="AmpR"

/ApEinfo_fwdcolor="yellow"

/ApEinfo_revcolor="yellow"

/ApEinfo_graphicformat="arrow_data {{0 1 2 0 0 -1} {} 0}

width 5 offset 0"

misc_feature	join(5472..5781,5786..6070)

/locus_tag="PbEF1a 5'UTR"

/label="PbEF1a 5'UTR"

/ApEinfo_label="PbEF1a 5'UTR"

/ApEinfo_fwdcolor="#804040"

/ApEinfo_revcolor="#804040"

/ApEinfo_graphicformat="arrow_data {{0 1 2 0 0 -1} {} 0}

width 5 offset 0"

misc_feature	1..34

/locus_tag="loxP Site"

/label="loxP Site"

/ApEinfo_label="loxP Site"

/ApEinfo_fwdcolor="#ffff00"

/ApEinfo_revcolor="green"

/ApEinfo_graphicformat="arrow_data {{0 1 2 0 0 -1} {} 0}

width 5 offset 0"

misc_feature	1199..1237

/locus_tag="loxP Site(1)"

/label="loxP Site(1)"

/ApEinfo_label="loxP Site"

/ApEinfo_fwdcolor="#ffff00"

/ApEinfo_revcolor="green"

/ApEinfo_graphicformat="arrow_data {{0 1 2 0 0 -1} {} 0}

width 5 offset 0"

misc_feature	41..609

/locus_tag="3' HR"

/label="3' HR"

/ApEinfo_label="3' HR"

/ApEinfo_fwdcolor="#8080c0"

/ApEinfo_revcolor="green"

/ApEinfo_graphicformat="arrow_data {{0 0.5 0 1 2 0 0 -1 0

-0.5} {} 0} width 5 offset 0"

ORIGIN

1 ATAACTTCGT ATAGCATACA TTATACGAAG TTATGGTACC GTTCTTCTTT GACTTACTGC

61 TGATTTAAAA GCGAAACTAT ATGTATAATA CTTGTTATAT ACAATTTATA CAATTAATCC

121 TATTGCCATT TGGAGGAGGA GTCATGCCTC CTTTTGGTTC TTGTAATTAA AAACAAAGAT

181 GAATTAATAC AAACTGTAGA CGACTTTTAG GCCTCGGGGT GCTGTAAACA TGAAAGTAAA

241 CTTAGTTTTA CGATCTGTTA AGGCTTATCC TCTGTGGTAA AGTATTTTTT TAAACTTAAA

301 TAGTATTTTT TTAAGCTTAA ATATAAATAA AAAAAATACA TATTATAATA TATTATAAAT

361 TTATATTGCA TAAGTTTGCA AATATAAATT TATGATATAT TATATATATT AACAATTTTT

421 AATATCAATT TTTTACATTT TTGATTATGC AATTATTTGT ATAAAAAAAA TATATATTAT

481 AATATATTAT AAATTTATAT TGCAAACTTA TGCAGTATAA ATTTATGATA TATTATATAT

541 ATTAACAATT TTTAATATCA ATTTTTTACA TTATTTATGC TCGTGATTGC ATGGATTTCC

601 TACTTAGGGG ATCGTCGACG ATCACCGGTG ATCCAGGCCA CCACATGTTA AGTAGTATAT

661 ATACTGTTGA AGTAGATACA AATATTAGTG TATGCTTCAG CAGTGGTGTA TTATTTACGC

721 CAAGTAAATG TATTGATGAA ATTCGAAGCA TCTTTGTGTT TATTTGTAAT TTAATTTTAG

781 AAAGTGTTTT TTATATGTAA TTGATGTAAA TATCATTTCC GAACTCGTAT GGGAAAGAAG

841 TATCAAGTCA ATTGCATATA AAAATAAAAT TGGACCAATG ATTGAAAAAT CGAAATTGTT

901 AATCATTATA TGATATTATA TAATAGTTGA TAATAAGTAG TTTAGTTAGT TTTAGTTATT

961 ATATGTAAAT TAATAATTTA TGTATGTTTC TGAAAGGTAC GAGGTTTTAA ATAATACTTT

1021 CAATTTTAAT TTTTTAATAT TCGAATGTAT AATGAAACGA CGACCAACCA ATGTATATAA

1081 TTATTTTAAA TAATTATATA TGCAATATGT AATAAATATG TTTTAGGATA TATTATAGCA

1141 TTTGAAAAAA TGTGTGTAGT GTGTTTGTAA AGTTATAGAA AACATCGTGG TGCTCGAGAT

1201 AACTTCGTAT AGCATACATT ATACGAAGTT ATGTCGAGAG CTTAATTCTT TTCGAGCTCT

1261 TTATGCTTAA GTTTACAATT TAATATTCAT ACTTTAAGTA TTTTTTGTAG TATCCTAGAT

1321 ATTGTGCTTT AAATGCTCAC CCCTCAAAGC ACCAGTAATA TTTTCATCCA CTGAAATACC

1381 ATTAAATTTT CAAAAAAATA CTATGCATAT AATGTTATAC ATATAAACAT AAAACGCCAT

1441 GTAAATCAAA AAATATATAA AAATATGTAT AAAAATAAAT ATGCACTAAA TATAAGCTAA

1501 TTATGCATAA AAATTAAAGT GCCCTTTATT AACTAGCTAG TCGTAATTAT TTATATTTCT

1561 ATGTTATAAA AAAATCCTCA TATAATAATA TAATTAATAT ATGTAATGTT TTTTTTATTT

1621 TATAATTTTA ATATAAAATA ATATGTAAAT TAATTCAAAA AATAAATATA ATTGTTGTGA

1681 AACAAAAAAC GTAATTTTTT CATTTGCCTT CAAAATTTAA ATTTATTTTA ATATTTCCTA

1741 AAATATATAT ACTTTGTGTA TAAATATATA AAAATATATA TTTGCTTATA AATAAATAAA

1801 AATTTTATAA AAATGACTAG TAGTAAAGGA GAAGAACTTT TCACTGGAGT TGTCCCAATT

1861 CTTGTTGAAT TAGATGGTGA TGTTAATGGG CACAAATTTT CTGTCAGTGG AGAGGGTGAA

1921 GGTGATGCAA CATACGGAAA ACTTACCCTT AAATTTATTT GCACTACTGG AAAACTACCT

1981 GTTCCATGGC CAACACTTGT CACTACTTTC GCGTATGGTC TTCAATGCTT TGCGAGATAC

2041 CCAGATCATA TGAAACAGCA TGACTTTTTC AAGAGTGCCA TGCCCGAAGG TTATGTACAG

2101 GAAAGAACTA TATTTTTCAA AGATGACGGG AACTACAAGA CACGTGCTGA AGTCAAGTTT

2161 GAAGGTGATA CCCTTGTTAA TAGAATCGAG TTAAAAGGTA TTGATTTTAA AGAAGATGGA

2221 AACATTCTTG GACACAAATT GGAATACAAC TATAACTCAC ACAATGTATA CATCATGGCA

2281 GACAAACAAA AGAATGGAAT CAAAGTTAAC TTCAAAATTA GACACAACAT TGAAGATGGA

2341 AGCGTTCAAC TAGCAGACCA TTATCAACAA AATACTCCAA TTGGCGATGG CCCTGTCCTT

2401 TTACCAGACA ACCATTACCT GTCCACACAA TCTGCCCTTT CGAAAGATCC CAACGAAAAG

2461 AGAGACCACA TGGTCCTTCT TGAGTTTGTA ACAGCTGCTG GGATTACACA TGGCATGGAT

2521 GAACTATACA AATAAGGATC CGTTTTTCTT ACTTATATAT TTATACCAAT TGATTGTATT

2581 TATAACTGTA AAAATGTGTA TGTTGTGTGC ATATTTTTTT TTGTGCATGC ACATGCATGT

2641 AAATAGCTAA AATTATGAAC ATTTTATTTT TTGTTCAGAA AAAAAAAACT TTACACACAT

2701 AAAATGGCTA GTATGAATAG CCATATTTTA TATAAATTAA ATCCTATGAA TTTATGACCA

2761 TATTAAAAAT TTAGATATTT ATGGAACATA ATATGTTTGA AACAATAAGA CAAAATTATT

2821 ATTATTATTA TTATTTTTAC TGTTATAATT ATGTTGTCTC TTCAATGATT CATAAATAGT

2881 TGGACTTGAT TTTTAAAATG TTTATAATAT GATTAGCATA GTTAAATAAA AAAAGTTGAA

2941 AAATTAAAAA AAAACATATA AACACAAATG ATGTTTTTTC CTTCAATTTC GGCGCCTGAT

3001 GCGGTATTTT CTCCTTACGC ATCTGTGCGG TATTTCACAC CGCATATGGT GCACTCTCAG

3061 TACAATCTGC TCTGATGCCG CATAGTTAAG CCAGCCCCGA CACCCGCCAA CACCCGCTGA

3121 CGCGCCCTGA CGGGCTTGTC TGCTCCCGGC ATCCGCTTAC AGACAAGCTG TGACCGTCTC

3181 CGGGAGCTGC ATGTGTCAGA GGTTTTCACC GTCATCACCG AAACGCGCGA GACGAAAGGG

3241 CCTCGTGATA CGCCTATTTT TATAGGTTAA TGTCATGATA ATAATGGTTT CTTAGACGTC

3301 AGGTGGCACT TTTCGGGGAA ATGTGCGCGG AACCCCTATT TGTTTATTTT TCTAAATACA

3361 TTCAAATATG TATCCGCTCA TGAGACAATA ACCCTGATAA ATGCTTCAAT AATATTGAAA

3421 AAGGAAGAGT ATGAGTATTC AACATTTCCG TGTCGCCCTT ATTCCCTTTT TTGCGGCATT

3481 TTGCCTTCCT GTTTTTGCTC ACCCAGAAAC GCTGGTGAAA GTAAAAGATG CTGAAGATCA

3541 GTTGGGTGCA CGAGTGGGTT ACATCGAACT GGATCTCAAC AGCGGTAAGA TCCTTGAGAG

3601 TTTTCGCCCC GAAGAACGTT TTCCAATGAT GAGCACTTTT AAAGTTCTGC TATGTGGCGC

3661 GGTATTATCC CGTATTGACG CCGGGCAAGA GCAACTCGGT CGCCGCATAC ACTATTCTCA

3721 GAATGACTTG GTTGAGTACT CACCAGTCAC AGAAAAGCAT CTTACGGATG GCATGACAGT

3781 AAGAGAATTA TGCAGTGCTG CCATAACCAT GAGTGATAAC ACTGCGGCCA ACTTACTTCT

3841 GACAACGATC GGAGGACCGA AGGAGCTAAC CGCTTTTTTG CACAACATGG GGGATCATGT

3901 AACTCGCCTT GATCGTTGGG AACCGGAGCT GAATGAAGCC ATACCAAACG ACGAGCGTGA

3961 CACCACGATG CCTGTAGCAA TGGCAACAAC GTTGCGCAAA CTATTAACTG GCGAACTACT

4021 TACTCTAGCT TCCCGGCAAC AATTAATAGA CTGGATGGAG GCGGATAAAG TTGCAGGACC

4081 ACTTCTGCGC TCGGCCCTTC CGGCTGGCTG GTTTATTGCT GATAAATCTG GAGCCGGTGA

4141 GCGTGGGTCT CGCGGTATCA TTGCAGCACT GGGGCCAGAT GGTAAGCCCT CCCGTATCGT

4201 AGTTATCTAC ACGACGGGGA GTCAGGCAAC TATGGATGAA CGAAATAGAC AGATCGCTGA

4261 GATAGGTGCC TCACTGATTA AGCATTGGTA ACTGTCAGAC CAAGTTTACT CATATATACT

4321 TTAGATTGAT TTAAAACTTC ATTTTTAATT TAAAAGGATC TAGGTGAAGA TCCTTTTTGA

4381 TAATCTCATG ACCAAAATCC CTTAACGTGA GTTTTCGTTC CACTGAGCGT CAGACCCCGT

4441 AGAAAAGATC AAAGGATCTT CTTGAGATCC TTTTTTTCTG CGCGTAATCT GCTGCTTGCA

4501 AACAAAAAAA CCACCGCTAC CAGCGGTGGT TTGTTTGCCG GATCAAGAGC TACCAACTCT

4561 TTTTCCGAAG GTAACTGGCT TCAGCAGAGC GCAGATACCA AATACTGTCC TTCTAGTGTA

4621 GCCGTAGTTA GGCCACCACT TCAAGAACTC TGTAGCACCG CCTACATACC TCGCTCTGCT

4681 AATCCTGTTA CCAGTGGCTG CTGCCAGTGG CGATAAGTCG TGTCTTACCG GGTTGGACTC

4741 AAGACGATAG TTACCGGATA AGGCGCAGCG GTCGGGCTGA ACGGGGGGTT CGTGCACACA

4801 GCCCAGCTTG GAGCGAACGA CCTACACCGA ACTGAGATAC CTACAGCGTG AGCATTGAGA

4861 AAGCGCCACG CTTCCCGAAG GGAGAAAGGC GGACAGGTAT CCGGTAAGCG GCAGGGTCGG

4921 AACAGGAGAG CGCACGAGGG AGCTTCCAGG GGGAAACGCC TGGTATCTTT ATAGTCCTGT

4981 CGGGTTTCGC CACCTCTGAC TTGAGCGTCG ATTTTTGTGA TGCTCGTCAG GGGGGCGGAG

5041 CCTATGGAAA AACGCCAGCA ACGCGGCCTT TTTACGGTTC CTGGCCTTTT GCTGGCCTTT

5101 TGCTCACATG TTCTTTCCTG CGTTATCCCC TGATTCTGTG GATAACCGTA TTACCGCCTT

5161 TGAGTGAGCT GATACCGCTC GCCGCAGCCG AACGACCGAG CGCAGCGAGT CAGTGAGCGA

5221 GGAAGCGGAA GAGCGCCCAA TACGCAAACC GCCTCTCCCC GCGCGTTGGC CGATTCATTA

5281 ATGCAGCTGG CACGACAGGT TTCCCGACTG GAAAGCGGGC AGTGAGCGCA ACGCAATTAA

5341 TGTGAGTTAG CTCACTCATT AGGCACCCCA GGCTTTACAC TTTATGCTTC CGGCTCGTAT

5401 GTTGTGTGGA ATTGTGAGCG GATAACAATT TCACACAGGA AACAGCTATG ACCATGATTA

5461 CGCCAAGCTT GATAATTCCT GCAGCCCAGC TTAATTCTTT TCGAGCTCTT TATGCTTAAG

5521 TTTACAATTT AATATTCATA CTTTAAGTAT TTTTTGTAGT ATCCTAGATA TTGTGCTTTA

5581 AATGCTCACC CCTCAAAGCA CCAGTAATAT TTTCATCCAC TGAAATACCA TTAAATTTTC

5641 AAAAAAATAC TATGCATATA ATGTTATACA TATAAACATA AAACGCCATG TAAATCAAAA

5701 AATATATAAA AATATGTATA AAAATAAATA TGCACTAAAT ATAAGCTAAT TATGCATAAA

5761 AATTAAAGTG CCCTTTATTA ACTAGCTAGT CGTAATTATT TATATTTCTA TGTTATAAAA

5821 AAATCCTCAT ATAATAATAT AATTAATATA TGTAATGTTT TTTTTATTTT ATAATTTTAA

5881 TATAAAATAA TATGTAAATT AATTCAAAAA ATAAATATAA TTGTTGTGAA ACAAAAAACG

5941 TAATTTTTTC ATTTGCCTTC AAAATTTAAA TTTATTTTAA TATTTCCTAA AATATATATA

6001 CTTTGTGTAT AAATATATAA AAATATATAT TTGCTTATAA ATAAATAAAA AATTTTATAA

6061 AACATAGGGG GATCCATGGT TGGTTCGCTA AACTGCATCG TCGCTGTGTC CCAGAACATG

6121 GGCATCGGCA AGAACGGGGA CCTGCCCTGG CCACCGCTCA GGAACGAATT TAGATATTTC

6181 CAGAGAATGA CCACAACCTC TTCAGTAGAA GGTAAACAGA ATCTGGTGAT TATGGGTAAG

6241 AAGACCTGGT TCTCCATTCC TGAGAAGAAT CGACCTTTAA AGGGTAGAAT TAATTTAGTT

6301 CTCAGCAGAG AACTCAAGGA ACCTCCACAA GGAGCTCATT TTCTTTCCAG AAGTCTAGAT

6361 GATGCCTTAA AACTTACTGA ACAACCAGAA TTAGCAAATA AAGTAGACAT GGTCTGGATA

6421 GTTGGTGGCA GTTCTGTTTA TAAGGAAGCC ATGAATCACC CAGGCCATCT TAAACTATTT

6481 GTGACAAGGA TCATGCAAGA CTTTGAAAGT GACACGTTTT TTCCAGAAAT TGATTTGGAG

6541 AAATATAAAC TTCTGCCAGA ATACCCAGGT GTTCTCTCTG ATGTCCAGGA GGAGAAAGGC

6601 ATTAAGTACA AATTTGAAGT ATATGAGAAG AATGATTAAG GATCCCGTTT TTCTTACTTA

6661 TATATTTATA CCAATTGATT GTATTTATAA CTGTAAAAAT GTGTATGTTG TGTGCATATT

6721 TTTTTTTGTG CATGCACATG CATGTAAATA GCTAAAATTA TGAACATTTT ATTTTTTGTT

6781 CAGAAAAAAA AAACTTTACA CACATAAAAT GGCTAGTATG AATAGCCATA TTTTATATAA

6841 ATTAAATCCT ATGAATTTAT GACCATATTA AAAATTTAGA TATTTATGGA ACATAATATG

6901 TTTGAAACAA TAAGACAAAA TTATTATTAT TATTATTATT TTTACTGTTA TAATTATGTT

6961 GTCTCTTCAA TGATTCATAA ATAGTTGGAC TTGATTTTTA AAATGTTTAT AATATGATTA

7021 GCATAGTTAA ATAAAAAAAG TTGAAAAATT AAAAAAAAAC ATATAAACAC AAATGATGTT

7081 TTTTCCTTCA ATTTCGATG

//

**pSL1556**

LOCUS	pSL1556__Chr6_S_	7258 bp	DNA	linear	07-JUN-2021

DEFINITION .

ACCESSION

VERSION

SOURCE	.

ORGANISM .

COMMENT

COMMENT

COMMENT	ApEinfo:methylated:1

FEATURES	Location/Qualifiers

exon	6235..6512

/vntifkey="61"

/locus_tag="hDHFR"

/label="hDHFR"

/ApEinfo_label="hDHFR"

/ApEinfo_fwdcolor="pink"

/ApEinfo_revcolor="pink"

/ApEinfo_graphicformat="arrow_data {{0 1 2 0 0 -1} {} 0}

width 5 offset 0"

exon	6513..6798

/vntifkey="61"

/locus_tag="hDHFR(1)"

/label="hDHFR(1)"

/ApEinfo_label="hDHFR"

/ApEinfo_fwdcolor="pink"

/ApEinfo_revcolor="pink"

/ApEinfo_graphicformat="arrow_data {{0 1 2 0 0 -1} {} 0}

width 5 offset 0"

misc_feature	join(5631..5940,5945..6234)

/vntifkey="21"

/locus_tag="PbEF1a-A\5'UTR"

/label="PbEF1a-A\5'UTR"

/ApEinfo_label="PbEF1a-A\5'UTR"

/ApEinfo_fwdcolor="#eb0214"

/ApEinfo_revcolor="#eb0214"

/ApEinfo_graphicformat="arrow_data {{0 1 2 0 0 -1} {} 0}

width 5 offset 0"

misc_feature	6799..7255

/vntifkey="21"

/locus_tag="PbDHFR/TS\3'UTR"

/label="PbDHFR/TS\3'UTR"

/ApEinfo_label="PbDHFR/TS\3'UTR"

/ApEinfo_fwdcolor="#eb0214"

/ApEinfo_revcolor="#eb0214"

/ApEinfo_graphicformat="arrow_data {{0 1 2 0 0 -1} {} 0}

width 5 offset 0"

misc_feature	1398..1971

/locus_tag="PbEF1a promoter (from pSL0281)"

/label="PbEF1a promoter (from pSL0281)"

/ApEinfo_label="PbEF1a promoter (from pSL0281)"

/ApEinfo_fwdcolor="cyan"

/ApEinfo_revcolor="green"

/ApEinfo_graphicformat="arrow_data {{0 1 2 0 0 -1} {} 0}

width 5 offset 0"

misc_feature	6805..7255

/locus_tag="PbDHFR-TS 3'UTR"

/label="PbDHFR-TS 3'UTR"

/ApEinfo_label="PbDHFR-TS 3'UTR"

/ApEinfo_fwdcolor="cyan"

/ApEinfo_revcolor="green"

/ApEinfo_graphicformat="arrow_data {{0 1 2 0 0 -1} {} 0}

width 5 offset 0"

misc_feature	2016..2694

/locus_tag="GFPmut2"

/label="GFPmut2"

/ApEinfo_label="GFPmut2"

/ApEinfo_fwdcolor="cyan"

/ApEinfo_revcolor="green"

/ApEinfo_graphicformat="arrow_data {{0 1 2 0 0 -1} {} 0}

width 5 offset 0"

misc_feature	6806..7253

/locus_tag="PbDHFR-TS 3'UTR(1)"

/label="PbDHFR-TS 3'UTR(1)"

/ApEinfo_label="PbDHFR-TS 3'UTR"

/ApEinfo_fwdcolor="cyan"

/ApEinfo_revcolor="green"

/ApEinfo_graphicformat="arrow_data {{0 1 2 0 0 -1} {} 0}

width 5 offset 0"

misc_feature	812..1351

/locus_tag="5' HR for KO"

/label="5' HR for KO"

/ApEinfo_label="5' HR for KO"

/ApEinfo_fwdcolor="#f5010a"

/ApEinfo_revcolor="green"

/ApEinfo_graphicformat="arrow_data {{0 1 2 0 0 -1} {} 0}

width 5 offset 0"

misc_feature	2701..3148

/locus_tag="PbDHFR/TS\3'UTR(1)"

/label="PbDHFR/TS\3'UTR(1)"

/ApEinfo_label="PbDHFR/TS\3'UTR"

/ApEinfo_fwdcolor="#ff0000"

/ApEinfo_revcolor="green"

/ApEinfo_graphicformat="arrow_data {{0 1 2 0 0 -1} {} 0}

width 5 offset 0"

rep_origin	4545..5227

/locus_tag="ColE1 origin"

/label="ColE1 origin"

/ApEinfo_label="ColE1 origin"

/ApEinfo_fwdcolor="gray50"

/ApEinfo_revcolor="gray50"

/ApEinfo_graphicformat="arrow_data {{0 1 2 0 0 -1} {} 0}

width 5 offset 0"

CDS	3788..4447

/locus_tag="AmpR"

/label="AmpR"

/ApEinfo_label="AmpR"

/ApEinfo_fwdcolor="yellow"

/ApEinfo_revcolor="yellow"

/ApEinfo_graphicformat="arrow_data {{0 1 2 0 0 -1} {} 0}

width 5 offset 0"

misc_feature	join(5631..5940,5945..6229)

/locus_tag="PbEF1a 5'UTR"

/label="PbEF1a 5'UTR"

/ApEinfo_label="PbEF1a 5'UTR"

/ApEinfo_fwdcolor="#804040"

/ApEinfo_revcolor="#804040"

/ApEinfo_graphicformat="arrow_data {{0 1 2 0 0 -1} {} 0}

width 5 offset 0"

misc_feature	1..34

/locus_tag="loxP Site"

/label="loxP Site"

/ApEinfo_label="loxP Site"

/ApEinfo_fwdcolor="#ffff00"

/ApEinfo_revcolor="green"

/ApEinfo_graphicformat="arrow_data {{0 1 2 0 0 -1} {} 0}

width 5 offset 0"

misc_feature	1358..1396

/locus_tag="loxP Site(1)"

/label="loxP Site(1)"

/ApEinfo_label="loxP Site"

/ApEinfo_fwdcolor="#ffff00"

/ApEinfo_revcolor="green"

/ApEinfo_graphicformat="arrow_data {{0 1 2 0 0 -1} {} 0}

width 5 offset 0"

misc_feature	41..787

/locus_tag="3' HR KO"

/label="3' HR KO"

/ApEinfo_label="3' HR KO"

/ApEinfo_fwdcolor="#9b0a64"

/ApEinfo_revcolor="green"

/ApEinfo_graphicformat="arrow_data {{0 1 2 0 0 -1} {} 0}

width 5 offset 0"

ORIGIN

1 ATAACTTCGT ATAGCATACA TTATACGAAG TTATggtacc GAGTGAACAA GTAATAAGTA

61 TATATTTGGG AAGGTGGTAT ATTGTTTTAA TAGTGGGAGG GATGATATTA TTTAAAAATA

121 TGTTTAAGTT ATTATGTTTT TTTATAAGTA CAAAATTGTC ATAAAAAGAT AGTATGTATA

181 GACACTTTAT GTTAAAATAT AAATTATAAT ATATTATGAG TGAGGTAATT ATATATGTAA

241 ATATTATATA ATATAGTATT TTTTACATCT ATAAAAAATG GAAGGGAATG TGTAAAAATA

301 ATGGTAAAAA AACAATGTTG TATAGTAAGT AATAGTGTAT ATAAATATGT TAATTCTTTT

361 ATATGGTATA ATTTTCTTTA TTGTTCGGTG ATTGACTTTT TATATTTTTT ATTGAAATTT

421 TATTAAGTCA ATATTTGAAA TTTAGTATTA CTAAATGAGC ATATTTTATA AGGGTTATAA

481 TTCAAGTTCT AATATTCCTA TAAATATATA TGATGGTGAT TGGGTTGATT TTTTAATAAG

541 TAATTATACA TTTGGGGTTG TAATAATAAT AATAAACTAA TATTGATATT TATATATATA

601 TTATTCTCAA TTATAATATA ATAAATATGT AAAGATTGCT TATATTTATA GCCAGAATTT

661 ATTACAAAAT TATAAGTTGA ATTACAAAAT TATATGTTGA GTTTACTTTT TTTAAATATA

721 TAAGTTCAAT ATATTCAATA TATATTTTTG GATAGACAAA TGAAAATGGT TAAGTAGTAG

781 AAAAGGGgat cgtcgacgat caccggtgat cCATCCATAT ATACTCAATC GTCGGGAGGT

841 TGTAAAAGGG TACTTATTAA TTTCAAAAAA ATATATGGTT ATAAATCCAA TTATGTATAT

901 TTTATATATT AATATAAATA GTAAAAAATA AATGAATAAT AGTTTTTTGG GGACTCCACA

961 TGTTGAGCTA TAATAATAGT ATCAGTAGTA GAAGTAATAT AAGTAGTAAA AAGAAAATAT

1021 AAAAGTTAGA AATTTATATT TTTTAATATT GTTATTGTTT ATAAATTTAT ATTATTTAAT

1081 TATTTTTAAT TATTATTATT TTTAAAGATA AATAAAATAA ATAAGTTTTT TATGGGCTTG

1141 GCAATATGGA TGTTGTAATT TTCCAAAACT TGTTTAATTA CAAGTTCCTA CTATTGATAT

1201 TACATTACTA AAGTTAATTT TTAACATTAA TAATTTGTAT TAAATAAATA CAATTTATTT

1261 AAATCCTCAT TATTTTATAA TTTAATTAAT ATAAGATAGT GTTTTTATAT ATATTAGCAT

1321 TTCCGAATTT CCGTTTGGTA TTTATTGTGT GctcgagATA ACTTCGTATA GCATACATTA

1381 TACGAAGTTA TgtcgagAGC TTAATTCTTT TCGAGCTCTT TATGCTTAAG TTTACAATTT

1441 AATATTCATA CTTTAAGTAT TTTTTGTAGT ATCCTAGATA TTGTGCTTTA AATGCTCACC

1501 CCTCAAAGCA CCAGTAATAT TTTCATCCAC TGAAATACCA TTAAATTTTC AAAAAAATAC

1561 TATGCATATA ATGTTATACA TATAAACATA AAACGCCATG TAAATCAAAA AATATATAAA

1621 AATATGTATA AAAATAAATA TGCACTAAAT ATAAGCTAAT TATGCATAAA AATTAAAGTG

1681 CCCTTTATTA ACTAGctagT CGTAATTATT TATATTTCTA TGTTATAAAA AAATCCTCAT

1741 ATAATAATAT AATTAATATA TGTAATGTTT TTTTTATTTT ATAATTTTAA TATAAAATAA

1801 TATGTAAATT AATTCAAAAA ATAAATATAA TTGTTGTGAA ACAAAAAACG TAATTTTTTC

1861 ATTTGCCTTC AAAATTTAAA TTTATTTTAA TATTTCCTAA AATATATATA CTTTGTGTAT

1921 AAATATATAA AAATATATAT TTGCTTATAA ATAAATAAAA ATTTTATAAA Aatgactagt

1981 agtaaaggag aagaactttt cactggagtt gtcccAATTC TTGTTGAATT AGATGGTGAT

2041 GTTAATGGGC ACAAATTTTC TGTCAGTGGA GAGGGTGAAG GTGATGCAAC ATACGGAAAA

2101 CTTACCCTTA AATTTATTTG CACTACTGGA AAACTACCTG TTCCATGGCC AACACTTGTC

2161 ACTACTTTCG CGTATGGTCT TCAATGCTTT GCGAGATACC CAGATCATAT GAAACAGCAT

2221 GACTTTTTCA AGAGTGCCAT GCCCGAAGGT TATGTACAGG AAAGAACTAT ATTTTTCAAA

2281 GATGACGGGA ACTACAAGAC ACGTGCTGAA GTCAAGTTTG AAGGTGATAC CCTTGTTAAT

2341 AGAATCGAGT TAAAAGGTAT TGATTTTAAA GAAGATGGAA ACATTCTTGG ACACAAATTG

2401 GAATACAACT ATAACTCACA CAATGTATAC ATCATGGCAG ACAAACAAAA GAATGGAATC

2461 AAAGTTAACT TCAAAATTAG ACACAACATT GAAGATGGAA GCGTTCAACT AGCAGACCAT

2521 TATCAACAAA ATACTCCAAT TGGCGATGGC CCTGTCCTTT TACCAGACAA CCATTACCTG

2581 TCCACACAAT CTGCCCTTTC GAAAGATCCC AACGAAAAGA GAGACCACAT GGTCCTTCTT

2641 GAGTTTGTAA CAGCTGCTGG GATTACACAT GGCATGGATG AACTATACAA ATAAggatcc

2701 GTTTTTCTTA CTTATATATT TATACCAATT GATTGTATTT ATAACTGTAA AAATGTGTAT

2761 GTTGTGTGCA TATTTTTTTT TGTGCATGCA CATGCATGTA AATAGCTAAA ATTATGAACA

2821 TTTTATTTTT TGTTCAGAAA AAAAAAACTT TACACACATA AAATGGCTAG TATGAATAGC

2881 CATATTTTAT ATAAATTAAA TCCTATGAAT TTATGACCAT ATTAAAAATT TAGATATTTA

2941 TGGAACATAA TATGTTTGAA ACAATAAGAC AAAATTATTA TTATTATTAT TATTTTTACT

3001 GTTATAATTA TGTTGTCTCT TCAATGATTC ATAAATAGTT GGACTTGATT TTTAAAATGT

3061 TTATAATATG ATTAGCATAG TTAAATAAAA AAAGTTGAAA AATTAAAAAA AAACATATAA

3121 ACACAAATGA TGTTTTTTCC TTCAATTTcg gcgcctgatg cggtattttc tccttacgca

3181 tctgtgcggt atttcacacc gcatatggtg cactctcagt acaatctgct ctgatgccgc

3241 atagttaagc cagccccgac acccgccaac acccgctgac gcgccctgac gggcttgtct

3301 gctcccggca tccgcttaca gacaagctgt gaccgtctcc gggagctgca tgtgtcagag

3361 gttttcaccg tcatcaccga aacgcgcgag acgaaagggc ctcgtgatac gcctattttt

3421 ataggttaat gtcatgataa taatggtttc ttagacgtca ggtggcactt ttcggggaaa

3481 tgtgcgcgga acccctattt gtttattttt ctaaatacat tcaaatatgt atccgctcat

3541 gagacaataa ccctgataaa tgcttcaata atattgaaaa aggaagagta tgagtattca

3601 acatttccgt gtcgccctta ttcccttttt tgcggcattt tgccttcctg tttttgctca

3661 cccagaaacg ctggtgaaag taaaagatgc tgaagatcag ttgggtgcac gagtgggtta

3721 catcgaactg gatctcaaca gcggtaagat ccttgagagt tttcgccccg aagaacgttt

3781 tccaatgatg agcactttta aagttctgct atgtggcgcg gtattatccc gtattgacgc

3841 cgggcaagag caactcggtc gccgcataca ctattctcag aatgacttgg ttgagtactc

3901 accagtcaca gaaaagcatc ttacggatgg catgacagta agagaattat gcagtgctgc

3961 cataaccatg agtgataaca ctgcggccaa cttacttctg acaacgatcg gaggaccgaa

4021 ggagctaacc gcttttttgc acaacatggg ggatcatgta actcgccttg atcgttggga

4081 accggagctg aatgaagcca taccaaacga cgagcgtgac accacgatgc ctgtagcaat

4141 ggcaacaacg ttgcgcaaac tattaactgg cgaactactt actctagctt cccggcaaca

4201 attaatagac tggatggagg cggataaagt tgcaggacca cttctgcgct cggcccttcc

4261 ggctggctgg tttattgctg ataaatctgg agccggtgag cgtgggtctc gcggtatcat

4321 tgcagcactg gggccagatg gtaagccctc ccgtatcgta gttatctaca cgacggggag

4381 tcaggcaact atggatgaac gaaatagaca gatcgctgag ataggtgcct cactgattaa

4441 gcattggtaa ctgtcagacc aagtttactc atatatactt tagattgatt taaaacttca

4501 tttttaattt aaaaggatct aggtgaagat cctttttgat aatctcatga ccaaaatccc

4561 ttaacgtgag ttttcgttcc actgagcgtc agaccccgta gaaaagatca aaggatcttc

4621 ttgagatcct ttttttctgc gcgtaatctg ctgcttgcaa acaaaaaaac caccgctacc

4681 agcggtggtt tgtttgccgg atcaagagct accaactctt tttccgaagg taactggctt

4741 cagcagagcg cagataccaa atactgtcct tctagtgtag ccgtagttag gccaccactt

4801 caagaactct gtagcaccgc ctacatacct cgctctgcta atcctgttac cagtggctgc

4861 tgccagtggc gataagtcgt gtcttaccgg gttggactca agacgatagt taccggataa

4921 ggcgcagcgg tcgggctgaa cggggggttc gtgcacacag cccagcttgg agcgaacgac

4981 ctacaccgaa ctgagatacc tacagcgtga gcattgagaa agcgccacgc ttcccgaagg

5041 gagaaaggcg gacaggtatc cggtaagcgg cagggtcgga acaggagagc gcacgaggga

5101 gcttccaggg ggaaacgcct ggtatcttta tagtcctgtc gggtttcgcc acctctgact

5161 tgagcgtcga tttttgtgat gctcgtcagg ggggcggagc ctatggaaaa acgccagcaa

5221 cgcggccttt ttacggttcc tggccttttg ctggcctttt gctcacatgt tctttcctgc

5281 gttatcccct gattctgtgg ataaccgtat taccgccttt gagtgagctg ataccgctcg

5341 ccgcagccga acgaccgagc gcagcgagtc agtgagcgag gaagcggaag agcgcccaat

5401 acgcaaaccg cctctccccg cgcgttggcc gattcattaa tgcagctggc acgacaggtt

5461 tcccgactgg aaagcgggca gtgagcgcaa cgcaattaat gtgagttagc tcactcatta

5521 ggcaccccag gctttacact ttatgcttcc ggctcgtatg ttgtgtggaa ttgtgagcgg

5581 ataacaattt cacacaggaa acagctatga ccatgattac gccaagcttg ataattcctg

5641 cagcccagct taattctttt cgagctcttt atgcttaagt ttacaattta atattcatac

5701 tttaagtatt ttttgtagta tcctagatat tgtgctttaa atgctcaccc ctcaaagcac

5761 cagtaatatt ttcatccact gaaataccat taaattttca aaaaaatact atgcatataa

5821 tgttatacat ataaacataa aacgccatgt aaatcaaaaa atatataaaa atatgtataa

5881 aaataaatat gcactaaata taagctaatt atgcataaaa attaaagtgc cctttattaa

5941 ctagctagtc gtaattattt atatttctat gttataaaaa aatcctcata taataatata

6001 attaatatat gtaatgtttt ttttatttta taattttaat ataaaataat atgtaaatta

6061 attcaaaaaa taaatataat tgttgtgaaa caaaaaacgt aattttttca tttgccttca

6121 aaatttaaat ttattttaat atttcctaaa atatatatac tttgtgtata aatatataaa

6181 aatatatatt tgcttataaa taaataaaaa attttataaa acataggggg atccatggtt

6241 ggttcgctaa actgcatcgt cgctgtgtcc cagaacatgg gcatcggcaa gaacggggac

6301 ctgccctggc caccgctcag gaacgaattt agatatttcc agagaatgac cacaacctct

6361 tcagtagaag gtaaacagaa tctggtgatt atgggtaaga agacctggtt ctccattcct

6421 gagaagaatc gacctttaaa gggtagaatt aatttagttc tcagcagaga actcaaggaa

6481 cctccacaag gagctcattt tctttccaga agtctagatg atgccttaaa acttactgaa

6541 caaccagaat tagcaaataa agtagacatg gtctggatag ttggtggcag ttctgtttat

6601 aaggaagcca tgaatcaccc aggccatctt aaactatttg tgacaaggat catgcaagac

6661 tttgaaagtg acacgttttt tccagaaatt gatttggaga aatataaact tctgccagaa

6721 tacccaggtg ttctctctga tgtccaggag gagaaaggca ttaagtacaa atttgaagta

6781 tatgagaaga atgattaagg atcccgtttt tcttacttat atatttatac caattgattg

6841 tatttataac tgtaaaaatg tgtatgttgt gtgcatattt ttttttgtgc atgcacatgc

6901 atgtaaatag ctaaaattat gaacatttta ttttttgttc agaaaaaaaa aactttacac

6961 acataaaatg gctagtatga atagccatat tttatataaa ttaaatccta tgaatttatg

7021 accatattaa aaatttagat atttatggaa cataatatgt ttgaaacaat aagacaaaat

7081 tattattatt attattattt ttactgttat aattatgttg tctcttcaat gattcataaa

7141 tagttggact tgatttttaa aatgtttata atatgattag catagttaaa taaaaaaagt

7201 tgaaaaatta aaaaaaaaca tataaacaca aatgatgttt tttccttcaa tttcgatg

//

**pSL1593**

LOCUS	pSL1593__Chr12_A	7107 bp	DNA	linear	14-OCT-2024

DEFINITION .

ACCESSION

VERSION

SOURCE	.

ORGANISM .

COMMENT

COMMENT	ApEinfo:methylated:1

FEATURES	Location/Qualifiers

misc_feature	1..34

/locus_tag="loxP Site"

/label="loxP Site"

/ApEinfo_label="loxP Site"

/ApEinfo_fwdcolor="#ffff00"

/ApEinfo_revcolor="green"

/ApEinfo_graphicformat="arrow_data {{0 1 2 0 0 -1} {} 0}

width 5 offset 0"

exon	6084..6361

/vntifkey="61"

/locus_tag="hDHFR"

/label="hDHFR"

/ApEinfo_label="hDHFR"

/ApEinfo_fwdcolor="pink"

/ApEinfo_revcolor="pink"

/ApEinfo_graphicformat="arrow_data {{0 1 2 0 0 -1} {} 0}

width 5 offset 0"

misc_feature	41..550

/locus_tag="3' HR"

/label="3' HR"

/ApEinfo_label="3' HR 2"

/ApEinfo_fwdcolor="#80ff00"

/ApEinfo_revcolor="green"

/ApEinfo_graphicformat="arrow_data {{0 1 2 0 0 -1} {} 0}

width 5 offset 0"

exon	6362..6647

/vntifkey="61"

/locus_tag="hDHFR(1)"

/label="hDHFR(1)"

/ApEinfo_label="hDHFR"

/ApEinfo_fwdcolor="pink"

/ApEinfo_revcolor="pink"

/ApEinfo_graphicformat="arrow_data {{0 1 2 0 0 -1} {} 0}

width 5 offset 0"

misc_feature	2550..2997

/locus_tag="PbDHFR/TS\3'UTR"

/label="PbDHFR/TS\3'UTR"

/ApEinfo_label="PbDHFR/TS\3'UTR"

/ApEinfo_fwdcolor="#ff0000"

/ApEinfo_revcolor="green"

/ApEinfo_graphicformat="arrow_data {{0 1 2 0 0 -1} {} 0}

width 5 offset 0"

misc_feature	1207..1245

/locus_tag="loxP Site(1)"

/label="loxP Site(1)"

/ApEinfo_label="loxP Site"

/ApEinfo_fwdcolor="#ffff00"

/ApEinfo_revcolor="green"

/ApEinfo_graphicformat="arrow_data {{0 1 2 0 0 -1} {} 0}

width 5 offset 0"

misc_feature	join(5480..5789,5794..6083)

/vntifkey="21"

/locus_tag="PbEF1a-A\5'UTR"

/label="PbEF1a-A\5'UTR"

/ApEinfo_label="PbEF1a-A\5'UTR"

/ApEinfo_fwdcolor="#eb0214"

/ApEinfo_revcolor="#eb0214"

/ApEinfo_graphicformat="arrow_data {{0 1 2 0 0 -1} {} 0}

width 5 offset 0"

rep_origin	4394..5076

/locus_tag="ColE1 origin"

/label="ColE1 origin"

/ApEinfo_label="ColE1 origin"

/ApEinfo_fwdcolor="gray50"

/ApEinfo_revcolor="gray50"

/ApEinfo_graphicformat="arrow_data {{0 1 2 0 0 -1} {} 0}

width 5 offset 0"

misc_feature	6648..7104

/vntifkey="21"

/locus_tag="PbDHFR/TS\3'UTR(1)"

/label="PbDHFR/TS\3'UTR(1)"

/ApEinfo_label="PbDHFR/TS\3'UTR"

/ApEinfo_fwdcolor="#eb0214"

/ApEinfo_revcolor="#eb0214"

/ApEinfo_graphicformat="arrow_data {{0 1 2 0 0 -1} {} 0}

width 5 offset 0"

misc_feature	1247..1820

/locus_tag="PbEF1a promoter (from pSL0281)"

/label="PbEF1a promoter (from pSL0281)"

/ApEinfo_label="PbEF1a promoter (from pSL0281)"

/ApEinfo_fwdcolor="cyan"

/ApEinfo_revcolor="green"

/ApEinfo_graphicformat="arrow_data {{0 1 2 0 0 -1} {} 0}

width 5 offset 0"

CDS	3637..4296

/locus_tag="AmpR"

/label="AmpR"

/ApEinfo_label="AmpR"

/ApEinfo_fwdcolor="yellow"

/ApEinfo_revcolor="yellow"

/ApEinfo_graphicformat="arrow_data {{0 1 2 0 0 -1} {} 0}

width 5 offset 0"

misc_feature	575..1200

/locus_tag="5' HR"

/label="5' HR"

/ApEinfo_label="5' HR 2"

/ApEinfo_fwdcolor="#80ff00"

/ApEinfo_revcolor="green"

/ApEinfo_graphicformat="arrow_data {{0 1 2 0 0 -1} {} 0}

width 5 offset 0"

misc_feature	6654..7104

/locus_tag="PbDHFR-TS 3'UTR"

/label="PbDHFR-TS 3'UTR"

/ApEinfo_label="PbDHFR-TS 3'UTR"

/ApEinfo_fwdcolor="cyan"

/ApEinfo_revcolor="green"

/ApEinfo_graphicformat="arrow_data {{0 1 2 0 0 -1} {} 0}

width 5 offset 0"

misc_feature	join(5480..5789,5794..6078)

/locus_tag="PbEF1a 5'UTR"

/label="PbEF1a 5'UTR"

/ApEinfo_label="PbEF1a 5'UTR"

/ApEinfo_fwdcolor="#804040"

/ApEinfo_revcolor="#804040"

/ApEinfo_graphicformat="arrow_data {{0 1 2 0 0 -1} {} 0}

width 5 offset 0"

misc_feature	1865..2543

/locus_tag="GFPmut2"

/label="GFPmut2"

/ApEinfo_label="GFPmut2"

/ApEinfo_fwdcolor="cyan"

/ApEinfo_revcolor="green"

/ApEinfo_graphicformat="arrow_data {{0 1 2 0 0 -1} {} 0}

width 5 offset 0"

misc_feature	6655..7102

/locus_tag="PbDHFR-TS 3'UTR(1)"

/label="PbDHFR-TS 3'UTR(1)"

/ApEinfo_label="PbDHFR-TS 3'UTR"

/ApEinfo_fwdcolor="cyan"

/ApEinfo_revcolor="green"

/ApEinfo_graphicformat="arrow_data {{0 1 2 0 0 -1} {} 0}

width 5 offset 0"

ORIGIN

1 ATAACTTCGT ATAGCATACA TTATACGAAG TTATggtacc CTCATAAACA AGTTTATATA

61 TTTTATGAGG AGGGTATATA TATATCCATA GCGTGTGTGG AAGCTCTTAT ATAATTTTTA

121 AAAAATATAA GGAATATATT ATTGTAGTAT ATATATTATG TATATAACTA AACTATATTT

181 CGTTTAAAAG CATAAAAGAT TGTTTTGGTG ATCACTAAAT TTTTTGTTTT TTTTAACAAT

241 GAAGTGTTAT GTTTATTTGA ACATGAGTTC TAAGGTATAT TCAGGATATT TGCATTATAT

301 ATATACTCGG ATAAATGAAC CAAATTCCGC AATGCATTAG ATAATACCAT ATATTGACAG

361 GTTTAGTAAT ACACGGTTCC TTTTATAAGA CCAAATGGTG TGTGTTATTA ATAACAAATC

421 TCTGCCCCTG TTAATATTAA TATTGGAATT ACCATTTTAA CGAAAAAAAT AATAGTTGAT

481 TATCGGTTTA TTAAATAAAA ATTATATTTA TATGTTTTAC ACACTAGGGA ACTTTTACAC

541 TATTTTATCG gatcgtcgac gatcaccggt gatcCATGGT TAAATTGAAG GAAATATAAT

601 GGTTGTCTTA TTTTATGTTA AAAATAAAAT ACAATGGAGC AAAAAAATAT AAACGATCAT

661 TTAAAATTAA TTACTATGAT ATGCTTTGGG TATATAAATA CTTAAAAATA TATTCATTAT

721 TTGTAGTATT TTATATTAAA ATATTGTAAG TAATATATAT GATGTTAAAG TTAACATGTT

781 TCATGAATAT TATGTTATAT TCGATGATTA ACAACTGTTT ATAAAACACA AAAAGGTCTT

841 AAAACATAAA AGGTCTTAAA ACATAAAAGG TCTTAAAACA TAAAAGGTCT TAAAACATAA

901 AAGGTCTTAA AACATAAAAG GTCTTAAAAC ATAAAAGGTC TTAAAACATA AAAGGTCTTA

961 AAACACAAAA GGTCTTAAAA CATAAAAGGT CTTAAAACAC AAAAGGTCTT AAAACACAAA

1021 AGGTCTTAAA ACATAAAAAG GGATTTGCTA CATTATAAAA CAAGATTGAT GATGTAGTAA

1081 TAATTATGAA TATATTATTT AGTACGAATT TAACAATAAA GATGTTAAAT TGAATTTGAA

1141 TGTTTTTTAA GTGTAAGTTT TGTTTACGCC TTGGATAATT TAATATTGTA GCGCCTTTGG

1201 ctcgagATAA CTTCGTATAG CATACATTAT ACGAAGTTAT gtcgagAGCT TAATTCTTTT

1261 CGAGCTCTTT ATGCTTAAGT TTACAATTTA ATATTCATAC TTTAAGTATT TTTTGTAGTA

1321 TCCTAGATAT TGTGCTTTAA ATGCTCACCC CTCAAAGCAC CAGTAATATT TTCATCCACT

1381 GAAATACCAT TAAATTTTCA AAAAAATACT ATGCATATAA TGTTATACAT ATAAACATAA

1441 AACGCCATGT AAATCAAAAA ATATATAAAA ATATGTATAA AAATAAATAT GCACTAAATA

1501 TAAGCTAATT ATGCATAAAA ATTAAAGTGC CCTTTATTAA CTAGctagTC GTAATTATTT

1561 ATATTTCTAT GTTATAAAAA AATCCTCATA TAATAATATA ATTAATATAT GTAATGTTTT

1621 TTTTATTTTA TAATTTTAAT ATAAAATAAT ATGTAAATTA ATTCAAAAAA TAAATATAAT

1681 TGTTGTGAAA CAAAAAACGT AATTTTTTCA TTTGCCTTCA AAATTTAAAT TTATTTTAAT

1741 ATTTCCTAAA ATATATATAC TTTGTGTATA AATATATAAA AATATATATT TGCTTATAAA

1801 TAAATAAAAA TTTTATAAAA atgactagta gtaaaggaga agaacttttc actggagttg

1861 tcccAATTCT TGTTGAATTA GATGGTGATG TTAATGGGCA CAAATTTTCT GTCAGTGGAG

1921 AGGGTGAAGG TGATGCAACA TACGGAAAAC TTACCCTTAA ATTTATTTGC ACTACTGGAA

1981 AACTACCTGT TCCATGGCCA ACACTTGTCA CTACTTTCGC GTATGGTCTT CAATGCTTTG

2041 CGAGATACCC AGATCATATG AAACAGCATG ACTTTTTCAA GAGTGCCATG CCCGAAGGTT

2101 ATGTACAGGA AAGAACTATA TTTTTCAAAG ATGACGGGAA CTACAAGACA CGTGCTGAAG

2161 TCAAGTTTGA AGGTGATACC CTTGTTAATA GAATCGAGTT AAAAGGTATT GATTTTAAAG

2221 AAGATGGAAA CATTCTTGGA CACAAATTGG AATACAACTA TAACTCACAC AATGTATACA

2281 TCATGGCAGA CAAACAAAAG AATGGAATCA AAGTTAACTT CAAAATTAGA CACAACATTG

2341 AAGATGGAAG CGTTCAACTA GCAGACCATT ATCAACAAAA TACTCCAATT GGCGATGGCC

2401 CTGTCCTTTT ACCAGACAAC CATTACCTGT CCACACAATC TGCCCTTTCG AAAGATCCCA

2461 ACGAAAAGAG AGACCACATG GTCCTTCTTG AGTTTGTAAC AGCTGCTGGG ATTACACATG

2521 GCATGGATGA ACTATACAAA TAAggatccG TTTTTCTTAC TTATATATTT ATACCAATTG

2581 ATTGTATTTA TAACTGTAAA AATGTGTATG TTGTGTGCAT ATTTTTTTTT GTGCATGCAC

2641 ATGCATGTAA ATAGCTAAAA TTATGAACAT TTTATTTTTT GTTCAGAAAA AAAAAACTTT

2701 ACACACATAA AATGGCTAGT ATGAATAGCC ATATTTTATA TAAATTAAAT CCTATGAATT

2761 TATGACCATA TTAAAAATTT AGATATTTAT GGAACATAAT ATGTTTGAAA CAATAAGACA

2821 AAATTATTAT TATTATTATT ATTTTTACTG TTATAATTAT GTTGTCTCTT CAATGATTCA

2881 TAAATAGTTG GACTTGATTT TTAAAATGTT TATAATATGA TTAGCATAGT TAAATAAAAA

2941 AAGTTGAAAA ATTAAAAAAA AACATATAAA CACAAATGAT GTTTTTTCCT TCAATTTcgg

3001 cgcctgatgc ggtattttct ccttacgcat ctgtgcggta tttcacaccg catatggtgc

3061 actctcagta caatctgctc tgatgccgca tagttaagcc agccccgaca cccgccaaca

3121 cccgctgacg cgccctgacg ggcttgtctg ctcccggcat ccgcttacag acaagctgtg

3181 accgtctccg ggagctgcat gtgtcagagg ttttcaccgt catcaccgaa acgcgcgaga

3241 cgaaagggcc tcgtgatacg cctattttta taggttaatg tcatgataat aatggtttct

3301 tagacgtcag gtggcacttt tcggggaaat gtgcgcggaa cccctatttg tttatttttc

3361 taaatacatt caaatatgta tccgctcatg agacaataac cctgataaat gcttcaataa

3421 tattgaaaaa ggaagagtat gagtattcaa catttccgtg tcgcccttat tccctttttt

3481 gcggcatttt gccttcctgt ttttgctcac ccagaaacgc tggtgaaagt aaaagatgct

3541 gaagatcagt tgggtgcacg agtgggttac atcgaactgg atctcaacag cggtaagatc

3601 cttgagagtt ttcgccccga agaacgtttt ccaatgatga gcacttttaa agttctgcta

3661 tgtggcgcgg tattatcccg tattgacgcc gggcaagagc aactcggtcg ccgcatacac

3721 tattctcaga atgacttggt tgagtactca ccagtcacag aaaagcatct tacggatggc

3781 atgacagtaa gagaattatg cagtgctgcc ataaccatga gtgataacac tgcggccaac

3841 ttacttctga caacgatcgg aggaccgaag gagctaaccg cttttttgca caacatgggg

3901 gatcatgtaa ctcgccttga tcgttgggaa ccggagctga atgaagccat accaaacgac

3961 gagcgtgaca ccacgatgcc tgtagcaatg gcaacaacgt tgcgcaaact attaactggc

4021 gaactactta ctctagcttc ccggcaacaa ttaatagact ggatggaggc ggataaagtt

4081 gcaggaccac ttctgcgctc ggcccttccg gctggctggt ttattgctga taaatctgga

4141 gccggtgagc gtgggtctcg cggtatcatt gcagcactgg ggccagatgg taagccctcc

4201 cgtatcgtag ttatctacac gacggggagt caggcaacta tggatgaacg aaatagacag

4261 atcgctgaga taggtgcctc actgattaag cattggtaac tgtcagacca agtttactca

4321 tatatacttt agattgattt aaaacttcat ttttaattta aaaggatcta ggtgaagatc

4381 ctttttgata atctcatgac caaaatccct taacgtgagt tttcgttcca ctgagcgtca

4441 gaccccgtag aaaagatcaa aggatcttct tgagatcctt tttttctgcg cgtaatctgc

4501 tgcttgcaaa caaaaaaacc accgctacca gcggtggttt gtttgccgga tcaagagcta

4561 ccaactcttt ttccgaaggt aactggcttc agcagagcgc agataccaaa tactgtcctt

4621 ctagtgtagc cgtagttagg ccaccacttc aagaactctg tagcaccgcc tacatacctc

4681 gctctgctaa tcctgttacc agtggctgct gccagtggcg ataagtcgtg tcttaccggg

4741 ttggactcaa gacgatagtt accggataag gcgcagcggt cgggctgaac ggggggttcg

4801 tgcacacagc ccagcttgga gcgaacgacc tacaccgaac tgagatacct acagcgtgag

4861 cattgagaaa gcgccacgct tcccgaaggg agaaaggcgg acaggtatcc ggtaagcggc

4921 agggtcggaa caggagagcg cacgagggag cttccagggg gaaacgcctg gtatctttat

4981 agtcctgtcg ggtttcgcca cctctgactt gagcgtcgat ttttgtgatg ctcgtcaggg

5041 gggcggagcc tatggaaaaa cgccagcaac gcggcctttt tacggttcct ggccttttgc

5101 tggccttttg ctcacatgtt ctttcctgcg ttatcccctg attctgtgga taaccgtatt

5161 accgcctttg agtgagctga taccgctcgc cgcagccgaa cgaccgagcg cagcgagtca

5221 gtgagcgagg aagcggaaga gcgcccaata cgcaaaccgc ctctccccgc gcgttggccg

5281 attcattaat gcagctggca cgacaggttt cccgactgga aagcgggcag tgagcgcaac

5341 gcaattaatg tgagttagct cactcattag gcaccccagg ctttacactt tatgcttccg

5401 gctcgtatgt tgtgtggaat tgtgagcgga taacaatttc acacaggaaa cagctatgac

5461 catgattacg ccaagcttga taattcctgc agcccagctt aattcttttc gagctcttta

5521 tgcttaagtt tacaatttaa tattcatact ttaagtattt tttgtagtat cctagatatt

5581 gtgctttaaa tgctcacccc tcaaagcacc agtaatattt tcatccactg aaataccatt

5641 aaattttcaa aaaaatacta tgcatataat gttatacata taaacataaa acgccatgta

5701 aatcaaaaaa tatataaaaa tatgtataaa aataaatatg cactaaatat aagctaatta

5761 tgcataaaaa ttaaagtgcc ctttattaac tagctagtcg taattattta tatttctatg

5821 ttataaaaaa atcctcatat aataatataa ttaatatatg taatgttttt tttattttat

5881 aattttaata taaaataata tgtaaattaa ttcaaaaaat aaatataatt gttgtgaaac

5941 aaaaaacgta attttttcat ttgccttcaa aatttaaatt tattttaata tttcctaaaa

6001 tatatatact ttgtgtataa atatataaaa atatatattt gcttataaat aaataaaaaa

6061 ttttataaaa cataggggga tccatggttg gttcgctaaa ctgcatcgtc gctgtgtccc

6121 agaacatggg catcggcaag aacggggacc tgccctggcc accgctcagg aacgaattta

6181 gatatttcca gagaatgacc acaacctctt cagtagaagg taaacagaat ctggtgatta

6241 tgggtaagaa gacctggttc tccattcctg agaagaatcg acctttaaag ggtagaatta

6301 atttagttct cagcagagaa ctcaaggaac ctccacaagg agctcatttt ctttccagaa

6361 gtctagatga tgccttaaaa cttactgaac aaccagaatt agcaaataaa gtagacatgg

6421 tctggatagt tggtggcagt tctgtttata aggaagccat gaatcaccca ggccatctta

6481 aactatttgt gacaaggatc atgcaagact ttgaaagtga cacgtttttt ccagaaattg

6541 atttggagaa atataaactt ctgccagaat acccaggtgt tctctctgat gtccaggagg

6601 agaaaggcat taagtacaaa tttgaagtat atgagaagaa tgattaagga tcccgttttt

6661 cttacttata tatttatacc aattgattgt atttataact gtaaaaatgt gtatgttgtg

6721 tgcatatttt tttttgtgca tgcacatgca tgtaaatagc taaaattatg aacattttat

6781 tttttgttca gaaaaaaaaa actttacaca cataaaatgg ctagtatgaa tagccatatt

6841 ttatataaat taaatcctat gaatttatga ccatattaaa aatttagata tttatggaac

6901 ataatatgtt tgaaacaata agacaaaatt attattatta ttattatttt tactgttata

6961 attatgttgt ctcttcaatg attcataaat agttggactt gatttttaaa atgtttataa

7021 tatgattagc atagttaaat aaaaaaagtt gaaaaattaa aaaaaaacat ataaacacaa

7081 atgatgtttt ttccttcaat ttcgatg

//

**pSL1597**

LOCUS	pSL1597__Chr7_A_	7091 bp	DNA	circular	14-OCT-2024

DEFINITION .

ACCESSION

VERSION

SOURCE	.

ORGANISM .

COMMENT

COMMENT	ApEinfo:methylated:1

FEATURES	Location/Qualifiers

misc_feature	1..34

/locus_tag="loxP Site"

/label="loxP Site"

/ApEinfo_label="loxP Site"

/ApEinfo_fwdcolor="#ffff00"

/ApEinfo_revcolor="green"

/ApEinfo_graphicformat="arrow_data {{0 1 2 0 0 -1} {} 0}

width 5 offset 0"

misc_feature	41..617

/locus_tag="3' HR"

/label="3' HR"

/ApEinfo_label="3' HR"

/ApEinfo_fwdcolor="#ff80ff"

/ApEinfo_revcolor="green"

/ApEinfo_graphicformat="arrow_data {{0 0.5 0 1 2 0 0 -1 0

-0.5} {} 0} width 5 offset 0"

exon	6068..6345

/vntifkey="61"

/locus_tag="hDHFR"

/label="hDHFR"

/ApEinfo_label="hDHFR"

/ApEinfo_fwdcolor="pink"

/ApEinfo_revcolor="pink"

/ApEinfo_graphicformat="arrow_data {{0 1 2 0 0 -1} {} 0}

width 5 offset 0"

misc_feature	642..1184

/locus_tag="5' HR"

/label="5' HR"

/ApEinfo_label="5' HR 2"

/ApEinfo_fwdcolor="#00ff00"

/ApEinfo_revcolor="green"

/ApEinfo_graphicformat="arrow_data {{0 1 2 0 0 -1} {} 0}

width 5 offset 0"

exon	6346..6631

/vntifkey="61"

/locus_tag="hDHFR(1)"

/label="hDHFR(1)"

/ApEinfo_label="hDHFR"

/ApEinfo_fwdcolor="pink"

/ApEinfo_revcolor="pink"

/ApEinfo_graphicformat="arrow_data {{0 1 2 0 0 -1} {} 0}

width 5 offset 0"

misc_feature	1231..1804

/locus_tag="PbEF1a promoter (from pSL0281)"

/label="PbEF1a promoter (from pSL0281)"

/ApEinfo_label="PbEF1a promoter (from pSL0281)"

/ApEinfo_fwdcolor="cyan"

/ApEinfo_revcolor="green"

/ApEinfo_graphicformat="arrow_data {{0 1 2 0 0 -1} {} 0}

width 5 offset 0"

misc_feature	2534..2981

/locus_tag="PbDHFR/TS\3'UTR"

/label="PbDHFR/TS\3'UTR"

/ApEinfo_label="PbDHFR/TS\3'UTR"

/ApEinfo_fwdcolor="#ff0000"

/ApEinfo_revcolor="green"

/ApEinfo_graphicformat="arrow_data {{0 1 2 0 0 -1} {} 0}

width 5 offset 0"

CDS	3621..4280

/locus_tag="AmpR"

/label="AmpR"

/ApEinfo_label="AmpR"

/ApEinfo_fwdcolor="yellow"

/ApEinfo_revcolor="yellow"

/ApEinfo_graphicformat="arrow_data {{0 1 2 0 0 -1} {} 0}

width 5 offset 0"

misc_feature	1191..1229

/locus_tag="loxP Site(1)"

/label="loxP Site(1)"

/ApEinfo_label="loxP Site"

/ApEinfo_fwdcolor="#ffff00"

/ApEinfo_revcolor="green"

/ApEinfo_graphicformat="arrow_data {{0 1 2 0 0 -1} {} 0}

width 5 offset 0"

misc_feature	1849..2527

/locus_tag="GFPmut2"

/label="GFPmut2"

/ApEinfo_label="GFPmut2"

/ApEinfo_fwdcolor="cyan"

/ApEinfo_revcolor="green"

/ApEinfo_graphicformat="arrow_data {{0 1 2 0 0 -1} {} 0}

width 5 offset 0"

misc_feature	join(5464..5773,5778..6067)

/vntifkey="21"

/locus_tag="PbEF1a-A\5'UTR"

/label="PbEF1a-A\5'UTR"

/ApEinfo_label="PbEF1a-A\5'UTR"

/ApEinfo_fwdcolor="#eb0214"

/ApEinfo_revcolor="#eb0214"

/ApEinfo_graphicformat="arrow_data {{0 1 2 0 0 -1} {} 0}

width 5 offset 0"

rep_origin	4378..5060

/locus_tag="ColE1 origin"

/label="ColE1 origin"

/ApEinfo_label="ColE1 origin"

/ApEinfo_fwdcolor="gray50"

/ApEinfo_revcolor="gray50"

/ApEinfo_graphicformat="arrow_data {{0 1 2 0 0 -1} {} 0}

width 5 offset 0"

misc_feature	6638..7088

/locus_tag="PbDHFR-TS 3'UTR"

/label="PbDHFR-TS 3'UTR"

/ApEinfo_label="PbDHFR-TS 3'UTR"

/ApEinfo_fwdcolor="cyan"

/ApEinfo_revcolor="green"

/ApEinfo_graphicformat="arrow_data {{0 1 2 0 0 -1} {} 0}

width 5 offset 0"

misc_feature	6639..7086

/locus_tag="PbDHFR-TS 3'UTR(1)"

/label="PbDHFR-TS 3'UTR(1)"

/ApEinfo_label="PbDHFR-TS 3'UTR"

/ApEinfo_fwdcolor="cyan"

/ApEinfo_revcolor="green"

/ApEinfo_graphicformat="arrow_data {{0 1 2 0 0 -1} {} 0}

width 5 offset 0"

misc_feature	join(5464..5773,5778..6062)

/locus_tag="PbEF1a 5'UTR"

/label="PbEF1a 5'UTR"

/ApEinfo_label="PbEF1a 5'UTR"

/ApEinfo_fwdcolor="#804040"

/ApEinfo_revcolor="#804040"

/ApEinfo_graphicformat="arrow_data {{0 1 2 0 0 -1} {} 0}

width 5 offset 0"

misc_feature	6632..7088

/vntifkey="21"

/locus_tag="PbDHFR/TS\3'UTR(1)"

/label="PbDHFR/TS\3'UTR(1)"

/ApEinfo_label="PbDHFR/TS\3'UTR"

/ApEinfo_fwdcolor="#eb0214"

/ApEinfo_revcolor="#eb0214"

/ApEinfo_graphicformat="arrow_data {{0 1 2 0 0 -1} {} 0}

width 5 offset 0"

ORIGIN

1 ATAACTTCGT ATAGCATACA TTATACGAAG TTATggtacc GTTAAAGATT ATGTAATGTA

61 TTTCCAACAT GTATTTATAA TATTTTTTAT GTTATGATAT TTCCTATATT ATGTTAAAAC

121 TTATCTAATA ATTTTCCAAC TATTTTTATC GTATATTTTT AAGTTAGATA TTATAGATTA

181 TGTTAAAATT TATTATATTA AAATTATTTT TTTTATATTT TACACTAAGT TATTTTATTA

241 TTATATATAA AAGTAAAAAT CTTTTGTATA TACTATTAAT AATATTTATT AATACTTGTA

301 TTTATAATTT TTTTTTATAA TTAACTCTAA ATAATATACT CCTTATTTTT TGTTTAATCA

361 AAACTTGCAA TATTAAGTGC TCCTTAAAAT TTTTTATTTC CATAAAATCT TTTAGGTTTT

421 TTATATTTTT ATATTTTGGT GTGCTATAAA ATCTATGGGC TTTTACAAAA ATTAAAAAGT

481 AAAGGTTTTA TTATTTATAA AGTTCCTTTT TATTACATAT TATTATAAAA AATTGTTTTT

541 ATTTTTTATA GAAGTATTGA TTTAATAAGA TATTAATAAA TGAAAAGTAA GAGCGTTTAA

601 AAATGTATAA ATACAACgat cgtcgacgat caccggtgat cGTGTTTACG CATTAAAACT

661 TACTATGAAT AATCAAATAT ATAAATTAAT AAAATAAAAA TAGAGAAGCA TATGCATTTC

721 TTATAATAAT TGTTTGTGTT TTTCTAAACA CGCTATTGTT TTTTTTGCAT GTACCCAAAT

781 TTATCAACGT ATAATGCAAA TTATTCTATT TTTGTTCTTA AAAGGTGTTG AGTTCAATAA

841 ATAAACAATA TACATATATA GAAGAGATAG AAACTAATAT AATATATTTA CGTAATAATA

901 CAAAACTCAC AATATTATAA AATATTCCCA AATTATTATG TATTTCTCAT AATTAACAAG

961 CCCTTTTTTA TTTATAAAAG GGTTGTTCCT TTGTTTAAAC GTCTATACCC AAAGTTGAGT

1021 AATTACTCAA TTCATAAAAT AATCTTTAAC ATTTTTATAT TCATGATTAT AATACACATT

1081 TGTATAATTC ATTATAGGTA ACTTATATAA GATATATTCT TTGCCTTTTG TAATTTGCAT

1141 AACTTTGGTA AACAAGCATG CTGAACACAT TAAACTATGA AGTGctcgag ATAACTTCGT

1201 ATAGCATACA TTATACGAAG TTATgtcgag AGCTTAATTC TTTTCGAGCT CTTTATGCTT

1261 AAGTTTACAA TTTAATATTC ATACTTTAAG TATTTTTTGT AGTATCCTAG ATATTGTGCT

1321 TTAAATGCTC ACCCCTCAAA GCACCAGTAA TATTTTCATC CACTGAAATA CCATTAAATT

1381 TTCAAAAAAA TACTATGCAT ATAATGTTAT ACATATAAAC ATAAAACGCC ATGTAAATCA

1441 AAAAATATAT AAAAATATGT ATAAAAATAA ATATGCACTA AATATAAGCT AATTATGCAT

1501 AAAAATTAAA GTGCCCTTTA TTAACTAGct agTCGTAATT ATTTATATTT CTATGTTATA

1561 AAAAAATCCT CATATAATAA TATAATTAAT ATATGTAATG TTTTTTTTAT TTTATAATTT

1621 TAATATAAAA TAATATGTAA ATTAATTCAA AAAATAAATA TAATTGTTGT GAAACAAAAA

1681 ACGTAATTTT TTCATTTGCC TTCAAAATTT AAATTTATTT TAATATTTCC TAAAATATAT

1741 ATACTTTGTG TATAAATATA TAAAAATATA TATTTGCTTA TAAATAAATA AAAATTTTAT

1801 AAAAatgact agtagtaaag gagaagaact tttcactgga gttgtcccAA TTCTTGTTGA

1861 ATTAGATGGT GATGTTAATG GGCACAAATT TTCTGTCAGT GGAGAGGGTG AAGGTGATGC

1921 AACATACGGA AAACTTACCC TTAAATTTAT TTGCACTACT GGAAAACTAC CTGTTCCATG

1981 GCCAACACTT GTCACTACTT TCGCGTATGG TCTTCAATGC TTTGCGAGAT ACCCAGATCA

2041 TATGAAACAG CATGACTTTT TCAAGAGTGC CATGCCCGAA GGTTATGTAC AGGAAAGAAC

2101 TATATTTTTC AAAGATGACG GGAACTACAA GACACGTGCT GAAGTCAAGT TTGAAGGTGA

2161 TACCCTTGTT AATAGAATCG AGTTAAAAGG TATTGATTTT AAAGAAGATG GAAACATTCT

2221 TGGACACAAA TTGGAATACA ACTATAACTC ACACAATGTA TACATCATGG CAGACAAACA

2281 AAAGAATGGA ATCAAAGTTA ACTTCAAAAT TAGACACAAC ATTGAAGATG GAAGCGTTCA

2341 ACTAGCAGAC CATTATCAAC AAAATACTCC AATTGGCGAT GGCCCTGTCC TTTTACCAGA

2401 CAACCATTAC CTGTCCACAC AATCTGCCCT TTCGAAAGAT CCCAACGAAA AGAGAGACCA

2461 CATGGTCCTT CTTGAGTTTG TAACAGCTGC TGGGATTACA CATGGCATGG ATGAACTATA

2521 CAAATAAgga tccGTTTTTC TTACTTATAT ATTTATACCA ATTGATTGTA TTTATAACTG

2581 TAAAAATGTG TATGTTGTGT GCATATTTTT TTTTGTGCAT GCACATGCAT GTAAATAGCT

2641 AAAATTATGA ACATTTTATT TTTTGTTCAG AAAAAAAAAA CTTTACACAC ATAAAATGGC

2701 TAGTATGAAT AGCCATATTT TATATAAATT AAATCCTATG AATTTATGAC CATATTAAAA

2761 ATTTAGATAT TTATGGAACA TAATATGTTT GAAACAATAA GACAAAATTA TTATTATTAT

2821 TATTATTTTT ACTGTTATAA TTATGTTGTC TCTTCAATGA TTCATAAATA GTTGGACTTG

2881 ATTTTTAAAA TGTTTATAAT ATGATTAGCA TAGTTAAATA AAAAAAGTTG AAAAATTAAA

2941 AAAAAACATA TAAACACAAA TGATGTTTTT TCCTTCAATT Tcggcgcctg atgcggtatt

3001 ttctccttac gcatctgtgc ggtatttcac accgcatatg gtgcactctc agtacaatct

3061 gctctgatgc cgcatagtta agccagcccc gacacccgcc aacacccgct gacgcgccct

3121 gacgggcttg tctgctcccg gcatccgctt acagacaagc tgtgaccgtc tccgggagct

3181 gcatgtgtca gaggttttca ccgtcatcac cgaaacgcgc gagacgaaag ggcctcgtga

3241 tacgcctatt tttataggtt aatgtcatga taataatggt ttcttagacg tcaggtggca

3301 cttttcgggg aaatgtgcgc ggaaccccta tttgtttatt tttctaaata cattcaaata

3361 tgtatccgct catgagacaa taaccctgat aaatgcttca ataatattga aaaaggaaga

3421 gtatgagtat tcaacatttc cgtgtcgccc ttattccctt ttttgcggca ttttgccttc

3481 ctgtttttgc tcacccagaa acgctggtga aagtaaaaga tgctgaagat cagttgggtg

3541 cacgagtggg ttacatcgaa ctggatctca acagcggtaa gatccttgag agttttcgcc

3601 ccgaagaacg ttttccaatg atgagcactt ttaaagttct gctatgtggc gcggtattat

3661 cccgtattga cgccgggcaa gagcaactcg gtcgccgcat acactattct cagaatgact

3721 tggttgagta ctcaccagtc acagaaaagc atcttacgga tggcatgaca gtaagagaat

3781 tatgcagtgc tgccataacc atgagtgata acactgcggc caacttactt ctgacaacga

3841 tcggaggacc gaaggagcta accgcttttt tgcacaacat gggggatcat gtaactcgcc

3901 ttgatcgttg ggaaccggag ctgaatgaag ccataccaaa cgacgagcgt gacaccacga

3961 tgcctgtagc aatggcaaca acgttgcgca aactattaac tggcgaacta cttactctag

4021 cttcccggca acaattaata gactggatgg aggcggataa agttgcagga ccacttctgc

4081 gctcggccct tccggctggc tggtttattg ctgataaatc tggagccggt gagcgtgggt

4141 ctcgcggtat cattgcagca ctggggccag atggtaagcc ctcccgtatc gtagttatct

4201 acacgacggg gagtcaggca actatggatg aacgaaatag acagatcgct gagataggtg

4261 cctcactgat taagcattgg taactgtcag accaagttta ctcatatata ctttagattg

4321 atttaaaact tcatttttaa tttaaaagga tctaggtgaa gatccttttt gataatctca

4381 tgaccaaaat cccttaacgt gagttttcgt tccactgagc gtcagacccc gtagaaaaga

4441 tcaaaggatc ttcttgagat cctttttttc tgcgcgtaat ctgctgcttg caaacaaaaa

4501 aaccaccgct accagcggtg gtttgtttgc cggatcaaga gctaccaact ctttttccga

4561 aggtaactgg cttcagcaga gcgcagatac caaatactgt ccttctagtg tagccgtagt

4621 taggccacca cttcaagaac tctgtagcac cgcctacata cctcgctctg ctaatcctgt

4681 taccagtggc tgctgccagt ggcgataagt cgtgtcttac cgggttggac tcaagacgat

4741 agttaccgga taaggcgcag cggtcgggct gaacgggggg ttcgtgcaca cagcccagct

4801 tggagcgaac gacctacacc gaactgagat acctacagcg tgagcattga gaaagcgcca

4861 cgcttcccga agggagaaag gcggacaggt atccggtaag cggcagggtc ggaacaggag

4921 agcgcacgag ggagcttcca gggggaaacg cctggtatct ttatagtcct gtcgggtttc

4981 gccacctctg acttgagcgt cgatttttgt gatgctcgtc aggggggcgg agcctatgga

5041 aaaacgccag caacgcggcc tttttacggt tcctggcctt ttgctggcct tttgctcaca

5101 tgttctttcc tgcgttatcc cctgattctg tggataaccg tattaccgcc tttgagtgag

5161 ctgataccgc tcgccgcagc cgaacgaccg agcgcagcga gtcagtgagc gaggaagcgg

5221 aagagcgccc aatacgcaaa ccgcctctcc ccgcgcgttg gccgattcat taatgcagct

5281 ggcacgacag gtttcccgac tggaaagcgg gcagtgagcg caacgcaatt aatgtgagtt

5341 agctcactca ttaggcaccc caggctttac actttatgct tccggctcgt atgttgtgtg

5401 gaattgtgag cggataacaa tttcacacag gaaacagcta tgaccatgat tacgccaagc

5461 ttgataattc ctgcagccca gcttaattct tttcgagctc tttatgctta agtttacaat

5521 ttaatattca tactttaagt attttttgta gtatcctaga tattgtgctt taaatgctca

5581 cccctcaaag caccagtaat attttcatcc actgaaatac cattaaattt tcaaaaaaat

5641 actatgcata taatgttata catataaaca taaaacgcca tgtaaatcaa aaaatatata

5701 aaaatatgta taaaaataaa tatgcactaa atataagcta attatgcata aaaattaaag

5761 tgccctttat taactagcta gtcgtaatta tttatatttc tatgttataa aaaaatcctc

5821 atataataat ataattaata tatgtaatgt tttttttatt ttataatttt aatataaaat

5881 aatatgtaaa ttaattcaaa aaataaatat aattgttgtg aaacaaaaaa cgtaattttt

5941 tcatttgcct tcaaaattta aatttatttt aatatttcct aaaatatata tactttgtgt

6001 ataaatatat aaaaatatat atttgcttat aaataaataa aaaattttat aaaacatagg

6061 gggatccatg gttggttcgc taaactgcat cgtcgctgtg tcccagaaca tgggcatcgg

6121 caagaacggg gacctgccct ggccaccgct caggaacgaa tttagatatt tccagagaat

6181 gaccacaacc tcttcagtag aaggtaaaca gaatctggtg attatgggta agaagacctg

6241 gttctccatt cctgagaaga atcgaccttt aaagggtaga attaatttag ttctcagcag

6301 agaactcaag gaacctccac aaggagctca ttttctttcc agaagtctag atgatgcctt

6361 aaaacttact gaacaaccag aattagcaaa taaagtagac atggtctgga tagttggtgg

6421 cagttctgtt tataaggaag ccatgaatca cccaggccat cttaaactat ttgtgacaag

6481 gatcatgcaa gactttgaaa gtgacacgtt ttttccagaa attgatttgg agaaatataa

6541 acttctgcca gaatacccag gtgttctctc tgatgtccag gaggagaaag gcattaagta

6601 caaatttgaa gtatatgaga agaatgatta aggatcccgt ttttcttact tatatattta

6661 taccaattga ttgtatttat aactgtaaaa atgtgtatgt tgtgtgcata tttttttttg

6721 tgcatgcaca tgcatgtaaa tagctaaaat tatgaacatt ttattttttg ttcagaaaaa

6781 aaaaacttta cacacataaa atggctagta tgaatagcca tattttatat aaattaaatc

6841 ctatgaattt atgaccatat taaaaattta gatatttatg gaacataata tgtttgaaac

6901 aataagacaa aattattatt attattatta tttttactgt tataattatg ttgtctcttc

6961 aatgattcat aaatagttgg acttgatttt taaaatgttt ataatatgat tagcatagtt

7021 aaataaaaaa agttgaaaaa ttaaaaaaaa acatataaac acaaatgatg ttttttcctt

7081 caatttcgat g

//

**pSL1700**

LOCUS	pSL1700	7310 bp	DNA	circular	21-OCT-2024

DEFINITION .

ACCESSION

VERSION

SOURCE	.

ORGANISM .

COMMENT

COMMENT	ApEinfo:methylated:1

FEATURES	Location/Qualifiers

exon	6287..6564

/vntifkey="61"

/locus_tag="hDHFR"

/label="hDHFR"

/ApEinfo_label="hDHFR"

/ApEinfo_fwdcolor="pink"

/ApEinfo_revcolor="pink"

/ApEinfo_graphicformat="arrow_data {{0 1 2 0 0 -1} {} 0}

width 5 offset 0"

exon	6565..6850

/vntifkey="61"

/locus_tag="hDHFR(1)"

/label="hDHFR(1)"

/ApEinfo_label="hDHFR"

/ApEinfo_fwdcolor="pink"

/ApEinfo_revcolor="pink"

/ApEinfo_graphicformat="arrow_data {{0 1 2 0 0 -1} {} 0}

width 5 offset 0"

misc_feature	join(5683..5992,5997..6286)

/vntifkey="21"

/locus_tag="PbEF1a-A\5'UTR"

/label="PbEF1a-A\5'UTR"

/ApEinfo_label="PbEF1a-A\5'UTR"

/ApEinfo_fwdcolor="#eb0214"

/ApEinfo_revcolor="#eb0214"

/ApEinfo_graphicformat="arrow_data {{0 1 2 0 0 -1} {} 0}

width 5 offset 0"

misc_feature	6851..7307

/vntifkey="21"

/locus_tag="PbDHFR/TS\3'UTR"

/label="PbDHFR/TS\3'UTR"

/ApEinfo_label="PbDHFR/TS\3'UTR"

/ApEinfo_fwdcolor="#eb0214"

/ApEinfo_revcolor="#eb0214"

/ApEinfo_graphicformat="arrow_data {{0 1 2 0 0 -1} {} 0}

width 5 offset 0"

misc_feature	1450..2023

/locus_tag="PbEF1a promoter (from pSL0281)"

/label="PbEF1a promoter (from pSL0281)"

/ApEinfo_label="PbEF1a promoter (from pSL0281)"

/ApEinfo_fwdcolor="cyan"

/ApEinfo_revcolor="green"

/ApEinfo_graphicformat="arrow_data {{0 1 2 0 0 -1} {} 0}

width 5 offset 0"

misc_feature	6857..7307

/locus_tag="PbDHFR-TS 3'UTR"

/label="PbDHFR-TS 3'UTR"

/ApEinfo_label="PbDHFR-TS 3'UTR"

/ApEinfo_fwdcolor="cyan"

/ApEinfo_revcolor="green"

/ApEinfo_graphicformat="arrow_data {{0 1 2 0 0 -1} {} 0}

width 5 offset 0"

misc_feature	677..1403

/locus_tag="LSU Del A 5' HR"

/label="LSU Del A 5' HR"

/ApEinfo_label="LSU Del A 5' HR"

/ApEinfo_fwdcolor="#0080ff"

/ApEinfo_revcolor="green"

/ApEinfo_graphicformat="arrow_data {{0 0.5 0 1 2 0 0 -1 0

-0.5} {} 0} width 5 offset 0"

misc_feature	2068..2746

/locus_tag="GFPmut2"

/label="GFPmut2"

/ApEinfo_label="GFPmut2"

/ApEinfo_fwdcolor="cyan"

/ApEinfo_revcolor="green"

/ApEinfo_graphicformat="arrow_data {{0 1 2 0 0 -1} {} 0}

width 5 offset 0"

misc_feature	6858..7305

/locus_tag="PbDHFR-TS 3'UTR(1)"

/label="PbDHFR-TS 3'UTR(1)"

/ApEinfo_label="PbDHFR-TS 3'UTR"

/ApEinfo_fwdcolor="cyan"

/ApEinfo_revcolor="green"

/ApEinfo_graphicformat="arrow_data {{0 1 2 0 0 -1} {} 0}

width 5 offset 0"

misc_feature	2753..3200

/locus_tag="PbDHFR/TS\3'UTR(1)"

/label="PbDHFR/TS\3'UTR(1)"

/ApEinfo_label="PbDHFR/TS\3'UTR"

/ApEinfo_fwdcolor="#ff0000"

/ApEinfo_revcolor="green"

/ApEinfo_graphicformat="arrow_data {{0 1 2 0 0 -1} {} 0}

width 5 offset 0"

rep_origin	4597..5279

/locus_tag="ColE1 origin"

/label="ColE1 origin"

/ApEinfo_label="ColE1 origin"

/ApEinfo_fwdcolor="gray50"

/ApEinfo_revcolor="gray50"

/ApEinfo_graphicformat="arrow_data {{0 1 2 0 0 -1} {} 0}

width 5 offset 0"

CDS	3840..4499

/locus_tag="AmpR"

/label="AmpR"

/ApEinfo_label="AmpR"

/ApEinfo_fwdcolor="yellow"

/ApEinfo_revcolor="yellow"

/ApEinfo_graphicformat="arrow_data {{0 1 2 0 0 -1} {} 0}

width 5 offset 0"

misc_feature	join(5683..5992,5997..6281)

/locus_tag="PbEF1a 5'UTR"

/label="PbEF1a 5'UTR"

/ApEinfo_label="PbEF1a 5'UTR"

/ApEinfo_fwdcolor="#804040"

/ApEinfo_revcolor="#804040"

/ApEinfo_graphicformat="arrow_data {{0 1 2 0 0 -1} {} 0}

width 5 offset 0"

misc_feature	1..34

/locus_tag="loxP Site"

/label="loxP Site"

/ApEinfo_label="loxP Site"

/ApEinfo_fwdcolor="#ffff00"

/ApEinfo_revcolor="green"

/ApEinfo_graphicformat="arrow_data {{0 1 2 0 0 -1} {} 0}

width 5 offset 0"

misc_feature	1410..1448

/locus_tag="loxP Site(1)"

/label="loxP Site(1)"

/ApEinfo_label="loxP Site"

/ApEinfo_fwdcolor="#ffff00"

/ApEinfo_revcolor="green"

/ApEinfo_graphicformat="arrow_data {{0 1 2 0 0 -1} {} 0}

width 5 offset 0"

misc_feature	41..638

/locus_tag="3' HR"

/label="3' HR"

/ApEinfo_label="3' HR"

/ApEinfo_fwdcolor="#00ff00"

/ApEinfo_revcolor="green"

/ApEinfo_graphicformat="arrow_data {{0 0.5 0 1 2 0 0 -1 0

-0.5} {} 0} width 5 offset 0"

ORIGIN

1 ATAACTTCGT ATAGCATACA TTATACGAAG TTATggtacc GTTCTTCTTT GACTTACTGC

61 TGATTTAAAA GCGAAACTAT ATGTATAATA CTTGTTATAT ACAATTTATA CAATTAATCC

121 TATTGCCATT TGGAGGAGGA GTCATGCCTC CTTTTGGTTC TTGTAATTAA AAACAAAGAT

181 GAATTAATAC AAACTGTAGA CGACTTTTAG GCCTCGGGGT GCTGTAAACA TGAAAGTAAA

241 CTTAGTTTTA CGATCTGTTA AGGCTTATCC TCTGTGGTAA AGTATTTTTT TAAACTTAAA

301 TAGTATTTTT TTAAGCTTAA ATATAAATAA AAAAAATACA TATTATAATA TATTATAAAT

361 TTATATTGCA TAAGTTTGCA AATATAAATT TATGATATAT TATATATATT AACAATTTTT

421 AATATCAATT TTTTACATTT TTGATTATGC AATTATTTGT ATAAAAAAAA TATATATTAT

481 AATATATTAT AAATTTATAT TGCAAACTTA TGCAGTATAA ATTTATGATA TATTATATAT

541 ATTAACAATT TTTAATATCA ATTTTTTACA TTATTTATGC TCGTGATTGC ATGGATTTCC

601 TACTTAGGGT GATATTATCG ATCATATGTG TCGtgtcgga tcgtcgacga tcaccggtGA

661 GTATTTAAAA TTATATCTGC GCGTGGGTGT CAAAGCCTAC ACGTCAGCAT ATATTTTTCC

721 TCCACTGAAA AGTGTAGGTA ATCTTTATCA ATACATATCG TGATGGGGAT AGATTATTGC

781 AATTATTAAT CTTGAACGAG GAATGCCTAG TAAGCATGAT TCATCAGATT GTGCTGACTA

841 CGTCCCTGCC CTTTGTACAC ACCGCCCGTC GCTCCTACCG ATTGAAAGAT ATGATGAATT

901 GTTTGGACAA GAAAATAGAA ATTTTATTTT TATTTTTTTT GGAAGGACCG TAAATCCTAT

961 CTTTTAAAGG AAGGAGAAGT CGTAACAAGG TTTCCGTAGG TGAACCTGCG GAAGGATCAT

1021 TATTAATACG ATTTAAAGAT TTGTATACTA ATATTTAATG GATAATAATT TTGTATTTAT

1081 ATGAAATTGT TATTAATTTC ATTAAATAAT AGTATATTAA TCTTTGTATA TATTTTTATT

1141 ATTGTATTAA ATATATATAT ATTGATTTGC TATATATACT TGTATATGTA ATGAATTGAT

1201 TTATATATGT TTAGTATATA TAATAAAAAT ATATACATTA CAAGTCTTAA CTATCAATTT

1261 TTGCATAACT TTTATTTTAA TTATAAATTT TGTGTTTATT TTTTTATTAT AAATCGATTT

1321 AATATATTAT CTGTATTTAA CATATAAAAT TGTATTATAT AAAAATCTAA ATTTATCACA

1381 ATCTTATTAG CACAATCTTA ACGctcgagA TAACTTCGTA TAGCATACAT TATACGAAGT

1441 TATgtcgagA GCTTAATTCT TTTCGAGCTC TTTATGCTTA AGTTTACAAT TTAATATTCA

1501 TACTTTAAGT ATTTTTTGTA GTATCCTAGA TATTGTGCTT TAAATGCTCA CCCCTCAAAG

1561 CACCAGTAAT ATTTTCATCC ACTGAAATAC CATTAAATTT TCAAAAAAAT ACTATGCATA

1621 TAATGTTATA CATATAAACA TAAAACGCCA TGTAAATCAA AAAATATATA AAAATATGTA

1681 TAAAAATAAA TATGCACTAA ATATAAGCTA ATTATGCATA AAAATTAAAG TGCCCTTTAT

1741 TAACTAGcta gTCGTAATTA TTTATATTTC TATGTTATAA AAAAATCCTC ATATAATAAT

1801 ATAATTAATA TATGTAATGT TTTTTTTATT TTATAATTTT AATATAAAAT AATATGTAAA

1861 TTAATTCAAA AAATAAATAT AATTGTTGTG AAACAAAAAA CGTAATTTTT TCATTTGCCT

1921 TCAAAATTTA AATTTATTTT AATATTTCCT AAAATATATA TACTTTGTGT ATAAATATAT

1981 AAAAATATAT ATTTGCTTAT AAATAAATAA AAATTTTATA AAAatgacta gtagtaaagg

2041 agaagaactt ttcactggag ttgtcccAAT TCTTGTTGAA TTAGATGGTG ATGTTAATGG

2101 GCACAAATTT TCTGTCAGTG GAGAGGGTGA AGGTGATGCA ACATACGGAA AACTTACCCT

2161 TAAATTTATT TGCACTACTG GAAAACTACC TGTTCCATGG CCAACACTTG TCACTACTTT

2221 CGCGTATGGT CTTCAATGCT TTGCGAGATA CCCAGATCAT ATGAAACAGC ATGACTTTTT

2281 CAAGAGTGCC ATGCCCGAAG GTTATGTACA GGAAAGAACT ATATTTTTCA AAGATGACGG

2341 GAACTACAAG ACACGTGCTG AAGTCAAGTT TGAAGGTGAT ACCCTTGTTA ATAGAATCGA

2401 GTTAAAAGGT ATTGATTTTA AAGAAGATGG AAACATTCTT GGACACAAAT TGGAATACAA

2461 CTATAACTCA CACAATGTAT ACATCATGGC AGACAAACAA AAGAATGGAA TCAAAGTTAA

2521 CTTCAAAATT AGACACAACA TTGAAGATGG AAGCGTTCAA CTAGCAGACC ATTATCAACA

2581 AAATACTCCA ATTGGCGATG GCCCTGTCCT TTTACCAGAC AACCATTACC TGTCCACACA

2641 ATCTGCCCTT TCGAAAGATC CCAACGAAAA GAGAGACCAC ATGGTCCTTC TTGAGTTTGT

2701 AACAGCTGCT GGGATTACAC ATGGCATGGA TGAACTATAC AAATAAggat ccGTTTTTCT

2761 TACTTATATA TTTATACCAA TTGATTGTAT TTATAACTGT AAAAATGTGT ATGTTGTGTG

2821 CATATTTTTT TTTGTGCATG CACATGCATG TAAATAGCTA AAATTATGAA CATTTTATTT

2881 TTTGTTCAGA AAAAAAAAAC TTTACACACA TAAAATGGCT AGTATGAATA GCCATATTTT

2941 ATATAAATTA AATCCTATGA ATTTATGACC ATATTAAAAA TTTAGATATT TATGGAACAT

3001 AATATGTTTG AAACAATAAG ACAAAATTAT TATTATTATT ATTATTTTTA CTGTTATAAT

3061 TATGTTGTCT CTTCAATGAT TCATAAATAG TTGGACTTGA TTTTTAAAAT GTTTATAATA

3121 TGATTAGCAT AGTTAAATAA AAAAAGTTGA AAAATTAAAA AAAAACATAT AAACACAAAT

3181 GATGTTTTTT CCTTCAATTT cggcgcctga tgcggtattt tctccttacg catctgtgcg

3241 gtatttcaca ccgcatatgg tgcactctca gtacaatctg ctctgatgcc gcatagttaa

3301 gccagccccg acacccgcca acacccgctg acgcgccctg acgggcttgt ctgctcccgg

3361 catccgctta cagacaagct gtgaccgtct ccgggagctg catgtgtcag aggttttcac

3421 cgtcatcacc gaaacgcgcg agacgaaagg gcctcgtgat acgcctattt ttataggtta

3481 atgtcatgat aataatggtt tcttagacgt caggtggcac ttttcgggga aatgtgcgcg

3541 gaacccctat ttgtttattt ttctaaatac attcaaatat gtatccgctc atgagacaat

3601 aaccctgata aatgcttcaa taatattgaa aaaggaagag tatgagtatt caacatttcc

3661 gtgtcgccct tattcccttt tttgcggcat tttgccttcc tgtttttgct cacccagaaa

3721 cgctggtgaa agtaaaagat gctgaagatc agttgggtgc acgagtgggt tacatcgaac

3781 tggatctcaa cagcggtaag atccttgaga gttttcgccc cgaagaacgt tttccaatga

3841 tgagcacttt taaagttctg ctatgtggcg cggtattatc ccgtattgac gccgggcaag

3901 agcaactcgg tcgccgcata cactattctc agaatgactt ggttgagtac tcaccagtca

3961 cagaaaagca tcttacggat ggcatgacag taagagaatt atgcagtgct gccataacca

4021 tgagtgataa cactgcggcc aacttacttc tgacaacgat cggaggaccg aaggagctaa

4081 ccgctttttt gcacaacatg ggggatcatg taactcgcct tgatcgttgg gaaccggagc

4141 tgaatgaagc cataccaaac gacgagcgtg acaccacgat gcctgtagca atggcaacaa

4201 cgttgcgcaa actattaact ggcgaactac ttactctagc ttcccggcaa caattaatag

4261 actggatgga ggcggataaa gttgcaggac cacttctgcg ctcggccctt ccggctggct

4321 ggtttattgc tgataaatct ggagccggtg agcgtgggtc tcgcggtatc attgcagcac

4381 tggggccaga tggtaagccc tcccgtatcg tagttatcta cacgacgggg agtcaggcaa

4441 ctatggatga acgaaataga cagatcgctg agataggtgc ctcactgatt aagcattggt

4501 aactgtcaga ccaagtttac tcatatatac tttagattga tttaaaactt catttttaat

4561 ttaaaaggat ctaggtgaag atcctttttg ataatctcat gaccaaaatc ccttaacgtg

4621 agttttcgtt ccactgagcg tcagaccccg tagaaaagat caaaggatct tcttgagatc

4681 ctttttttct gcgcgtaatc tgctgcttgc aaacaaaaaa accaccgcta ccagcggtgg

4741 tttgtttgcc ggatcaagag ctaccaactc tttttccgaa ggtaactggc ttcagcagag

4801 cgcagatacc aaatactgtc cttctagtgt agccgtagtt aggccaccac ttcaagaact

4861 ctgtagcacc gcctacatac ctcgctctgc taatcctgtt accagtggct gctgccagtg

4921 gcgataagtc gtgtcttacc gggttggact caagacgata gttaccggat aaggcgcagc

4981 ggtcgggctg aacggggggt tcgtgcacac agcccagctt ggagcgaacg acctacaccg

5041 aactgagata cctacagcgt gagcattgag aaagcgccac gcttcccgaa gggagaaagg

5101 cggacaggta tccggtaagc ggcagggtcg gaacaggaga gcgcacgagg gagcttccag

5161 ggggaaacgc ctggtatctt tatagtcctg tcgggtttcg ccacctctga cttgagcgtc

5221 gatttttgtg atgctcgtca ggggggcgga gcctatggaa aaacgccagc aacgcggcct

5281 ttttacggtt cctggccttt tgctggcctt ttgctcacat gttctttcct gcgttatccc

5341 ctgattctgt ggataaccgt attaccgcct ttgagtgagc tgataccgct cgccgcagcc

5401 gaacgaccga gcgcagcgag tcagtgagcg aggaagcgga agagcgccca atacgcaaac

5461 cgcctctccc cgcgcgttgg ccgattcatt aatgcagctg gcacgacagg tttcccgact

5521 ggaaagcggg cagtgagcgc aacgcaatta atgtgagtta gctcactcat taggcacccc

5581 aggctttaca ctttatgctt ccggctcgta tgttgtgtgg aattgtgagc ggataacaat

5641 ttcacacagg aaacagctat gaccatgatt acgccaagct tgataattcc tgcagcccag

5701 cttaattctt ttcgagctct ttatgcttaa gtttacaatt taatattcat actttaagta

5761 ttttttgtag tatcctagat attgtgcttt aaatgctcac ccctcaaagc accagtaata

5821 ttttcatcca ctgaaatacc attaaatttt caaaaaaata ctatgcatat aatgttatac

5881 atataaacat aaaacgccat gtaaatcaaa aaatatataa aaatatgtat aaaaataaat

5941 atgcactaaa tataagctaa ttatgcataa aaattaaagt gccctttatt aactagctag

6001 tcgtaattat ttatatttct atgttataaa aaaatcctca tataataata taattaatat

6061 atgtaatgtt ttttttattt tataatttta atataaaata atatgtaaat taattcaaaa

6121 aataaatata attgttgtga aacaaaaaac gtaatttttt catttgcctt caaaatttaa

6181 atttatttta atatttccta aaatatatat actttgtgta taaatatata aaaatatata

6241 tttgcttata aataaataaa aaattttata aaacataggg ggatccatgg ttggttcgct

6301 aaactgcatc gtcgctgtgt cccagaacat gggcatcggc aagaacgggg acctgccctg

6361 gccaccgctc aggaacgaat ttagatattt ccagagaatg accacaacct cttcagtaga

6421 aggtaaacag aatctggtga ttatgggtaa gaagacctgg ttctccattc ctgagaagaa

6481 tcgaccttta aagggtagaa ttaatttagt tctcagcaga gaactcaagg aacctccaca

6541 aggagctcat tttctttcca gaagtctaga tgatgcctta aaacttactg aacaaccaga

6601 attagcaaat aaagtagaca tggtctggat agttggtggc agttctgttt ataaggaagc

6661 catgaatcac ccaggccatc ttaaactatt tgtgacaagg atcatgcaag actttgaaag

6721 tgacacgttt tttccagaaa ttgatttgga gaaatataaa cttctgccag aatacccagg

6781 tgttctctct gatgtccagg aggagaaagg cattaagtac aaatttgaag tatatgagaa

6841 gaatgattaa ggatcccgtt tttcttactt atatatttat accaattgat tgtatttata

6901 actgtaaaaa tgtgtatgtt gtgtgcatat ttttttttgt gcatgcacat gcatgtaaat

6961 agctaaaatt atgaacattt tattttttgt tcagaaaaaa aaaactttac acacataaaa

7021 tggctagtat gaatagccat attttatata aattaaatcc tatgaattta tgaccatatt

7081 aaaaatttag atatttatgg aacataatat gtttgaaaca ataagacaaa attattatta

7141 ttattattat ttttactgtt ataattatgt tgtctcttca atgattcata aatagttgga

7201 cttgattttt aaaatgttta taatatgatt agcatagtta aataaaaaaa gttgaaaaat

7261 taaaaaaaaa catataaaca caaatgatgt tttttccttc aatttcgatg

//

**pSL1701**

LOCUS	pSL1701	6929 bp	DNA	linear	21-OCT-2024

DEFINITION .

ACCESSION

VERSION

SOURCE	.

ORGANISM .

COMMENT

COMMENT

COMMENT	ApEinfo:methylated:1

FEATURES	Location/Qualifiers

exon	5906..6183

/vntifkey="61"

/locus_tag="hDHFR"

/label="hDHFR"

/ApEinfo_label="hDHFR"

/ApEinfo_fwdcolor="pink"

/ApEinfo_revcolor="pink"

/ApEinfo_graphicformat="arrow_data {{0 1 2 0 0 -1} {} 0}

width 5 offset 0"

exon	6184..6469

/vntifkey="61"

/locus_tag="hDHFR(1)"

/label="hDHFR(1)"

/ApEinfo_label="hDHFR"

/ApEinfo_fwdcolor="pink"

/ApEinfo_revcolor="pink"

/ApEinfo_graphicformat="arrow_data {{0 1 2 0 0 -1} {} 0}

width 5 offset 0"

misc_feature	join(5302..5611,5616..5905)

/vntifkey="21"

/locus_tag="PbEF1a-A\5'UTR"

/label="PbEF1a-A\5'UTR"

/ApEinfo_label="PbEF1a-A\5'UTR"

/ApEinfo_fwdcolor="#eb0214"

/ApEinfo_revcolor="#eb0214"

/ApEinfo_graphicformat="arrow_data {{0 1 2 0 0 -1} {} 0}

width 5 offset 0"

misc_feature	6470..6926

/vntifkey="21"

/locus_tag="PbDHFR/TS\3'UTR"

/label="PbDHFR/TS\3'UTR"

/ApEinfo_label="PbDHFR/TS\3'UTR"

/ApEinfo_fwdcolor="#eb0214"

/ApEinfo_revcolor="#eb0214"

/ApEinfo_graphicformat="arrow_data {{0 1 2 0 0 -1} {} 0}

width 5 offset 0"

misc_feature	1069..1642

/locus_tag="PbEF1a promoter (from pSL0281)"

/label="PbEF1a promoter (from pSL0281)"

/ApEinfo_label="PbEF1a promoter (from pSL0281)"

/ApEinfo_fwdcolor="cyan"

/ApEinfo_revcolor="green"

/ApEinfo_graphicformat="arrow_data {{0 1 2 0 0 -1} {} 0}

width 5 offset 0"

misc_feature	6476..6926

/locus_tag="PbDHFR-TS 3'UTR"

/label="PbDHFR-TS 3'UTR"

/ApEinfo_label="PbDHFR-TS 3'UTR"

/ApEinfo_fwdcolor="cyan"

/ApEinfo_revcolor="green"

/ApEinfo_graphicformat="arrow_data {{0 1 2 0 0 -1} {} 0}

width 5 offset 0"

misc_feature	1687..2365

/locus_tag="GFPmut2"

/label="GFPmut2"

/ApEinfo_label="GFPmut2"

/ApEinfo_fwdcolor="cyan"

/ApEinfo_revcolor="green"

/ApEinfo_graphicformat="arrow_data {{0 1 2 0 0 -1} {} 0}

width 5 offset 0"

misc_feature	6477..6924

/locus_tag="PbDHFR-TS 3'UTR(1)"

/label="PbDHFR-TS 3'UTR(1)"

/ApEinfo_label="PbDHFR-TS 3'UTR"

/ApEinfo_fwdcolor="cyan"

/ApEinfo_revcolor="green"

/ApEinfo_graphicformat="arrow_data {{0 1 2 0 0 -1} {} 0}

width 5 offset 0"

misc_feature	2372..2819

/locus_tag="PbDHFR/TS\3'UTR(1)"

/label="PbDHFR/TS\3'UTR(1)"

/ApEinfo_label="PbDHFR/TS\3'UTR"

/ApEinfo_fwdcolor="#ff0000"

/ApEinfo_revcolor="green"

/ApEinfo_graphicformat="arrow_data {{0 1 2 0 0 -1} {} 0}

width 5 offset 0"

rep_origin	4216..4898

/locus_tag="ColE1 origin"

/label="ColE1 origin"

/ApEinfo_label="ColE1 origin"

/ApEinfo_fwdcolor="gray50"

/ApEinfo_revcolor="gray50"

/ApEinfo_graphicformat="arrow_data {{0 1 2 0 0 -1} {} 0}

width 5 offset 0"

misc_feature	635..1022

/locus_tag="PyChr5 LSU Del 5'HR B"

/label="PyChr5 LSU Del 5'HR B"

/ApEinfo_label="PyChr5 LSU Del 5'HR B"

/ApEinfo_fwdcolor="#ff8000"

/ApEinfo_revcolor="green"

/ApEinfo_graphicformat="arrow_data {{0 0.5 0 1 2 0 0 -1 0

-0.5} {} 0} width 5 offset 0"

CDS	3459..4118

/locus_tag="AmpR"

/label="AmpR"

/ApEinfo_label="AmpR"

/ApEinfo_fwdcolor="yellow"

/ApEinfo_revcolor="yellow"

/ApEinfo_graphicformat="arrow_data {{0 1 2 0 0 -1} {} 0}

width 5 offset 0"

misc_feature	join(5302..5611,5616..5900)

/locus_tag="PbEF1a 5'UTR"

/label="PbEF1a 5'UTR"

/ApEinfo_label="PbEF1a 5'UTR"

/ApEinfo_fwdcolor="#804040"

/ApEinfo_revcolor="#804040"

/ApEinfo_graphicformat="arrow_data {{0 1 2 0 0 -1} {} 0}

width 5 offset 0"

misc_feature	1..34

/locus_tag="loxP Site"

/label="loxP Site"

/ApEinfo_label="loxP Site"

/ApEinfo_fwdcolor="#ffff00"

/ApEinfo_revcolor="green"

/ApEinfo_graphicformat="arrow_data {{0 1 2 0 0 -1} {} 0}

width 5 offset 0"

misc_feature	1029..1067

/locus_tag="loxP Site(1)"

/label="loxP Site(1)"

/ApEinfo_label="loxP Site"

/ApEinfo_fwdcolor="#ffff00"

/ApEinfo_revcolor="green"

/ApEinfo_graphicformat="arrow_data {{0 1 2 0 0 -1} {} 0}

width 5 offset 0"

misc_feature	41..614

/locus_tag="3' HR"

/label="3' HR"

/ApEinfo_label="3' HR"

/ApEinfo_fwdcolor="#ffff00"

/ApEinfo_revcolor="green"

/ApEinfo_graphicformat="arrow_data {{0 0.5 0 1 2 0 0 -1 0

-0.5} {} 0} width 5 offset 0"

ORIGIN

1 ATAACTTCGT ATAGCATACA TTATACGAAG TTATggtacc GTTCTTCTTT GACTTACTGC

61 TGATTTAAAA GCGAAACTAT ATGTATAATA CTTGTTATAT ACAATTTATA CAATTAATCC

121 TATTGCCATT TGGAGGAGGA GTCATGCCTC CTTTTGGTTC TTGTAATTAA AAACAAAGAT

181 GAATTAATAC AAACTGTAGA CGACTTTTAG GCCTCGGGGT GCTGTAAACA TGAAAGTAAA

241 CTTAGTTTTA CGATCTGTTA AGGCTTATCC TCTGTGGTAA AGTATTTTTT TAAACTTAAA

301 TAGTATTTTT TTAAGCTTAA ATATAAATAA AAAAAATACA TATTATAATA TATTATAAAT

361 TTATATTGCA TAAGTTTGCA AATATAAATT TATGATATAT TATATATATT AACAATTTTT

421 AATATCAATT TTTTACATTT TTGATTATGC AATTATTTGT ATAAAAAAAA TATATATTAT

481 AATATATTAT AAATTTATAT TGCAAACTTA TGCAGTATAA ATTTATGATA TATTATATAT

541 ATTAACAATT TTTAATATCA ATTTTTTACA TTATTTATGC TCGTGATTGC ATGGATTTCC

601 TACTTAGGGt gtcggatcgt cgacgatcac cggtCGATTT AAAGATTTGT ATACTAATAT

661 TTAATGGATA ATAATTTTGT ATTTATATGA AATTGTTATT AATTTCATTA AATAATAGTA

721 TATTAATCTT TGTATATATT TTTATTATTG TATTAAATAT ATATATATTG ATTTGCTATA

781 TATACTTGTA TATGTAATGA ATTGATTTAT ATATGTTTAG TATATATAAT AAAAATATAT

841 ACATTACAAG TCTTAACTAT CAATTTTTGC ATAACTTTTA TTTTAATTAT AAATTTTGTG

901 TTTATTTTTT TATTATAAAT CGATTTAATA TATTATCTGT ATTTAACATA TAAAATTGTA

961 TTATATAAAA ATCTAAATTT ATCACAATCT TATTAGCACA ATCTTAACGA TGGATGTCTT

1021 GGctcgagAT AACTTCGTAT AGCATACATT ATACGAAGTT ATgtcgagAG CTTAATTCTT

1081 TTCGAGCTCT TTATGCTTAA GTTTACAATT TAATATTCAT ACTTTAAGTA TTTTTTGTAG

1141 TATCCTAGAT ATTGTGCTTT AAATGCTCAC CCCTCAAAGC ACCAGTAATA TTTTCATCCA

1201 CTGAAATACC ATTAAATTTT CAAAAAAATA CTATGCATAT AATGTTATAC ATATAAACAT

1261 AAAACGCCAT GTAAATCAAA AAATATATAA AAATATGTAT AAAAATAAAT ATGCACTAAA

1321 TATAAGCTAA TTATGCATAA AAATTAAAGT GCCCTTTATT AACTAGctag TCGTAATTAT

1381 TTATATTTCT ATGTTATAAA AAAATCCTCA TATAATAATA TAATTAATAT ATGTAATGTT

1441 TTTTTTATTT TATAATTTTA ATATAAAATA ATATGTAAAT TAATTCAAAA AATAAATATA

1501 ATTGTTGTGA AACAAAAAAC GTAATTTTTT CATTTGCCTT CAAAATTTAA ATTTATTTTA

1561 ATATTTCCTA AAATATATAT ACTTTGTGTA TAAATATATA AAAATATATA TTTGCTTATA

1621 AATAAATAAA AATTTTATAA AAatgactag tagtaaagga gaagaacttt tcactggagt

1681 tgtcccAATT CTTGTTGAAT TAGATGGTGA TGTTAATGGG CACAAATTTT CTGTCAGTGG

1741 AGAGGGTGAA GGTGATGCAA CATACGGAAA ACTTACCCTT AAATTTATTT GCACTACTGG

1801 AAAACTACCT GTTCCATGGC CAACACTTGT CACTACTTTC GCGTATGGTC TTCAATGCTT

1861 TGCGAGATAC CCAGATCATA TGAAACAGCA TGACTTTTTC AAGAGTGCCA TGCCCGAAGG

1921 TTATGTACAG GAAAGAACTA TATTTTTCAA AGATGACGGG AACTACAAGA CACGTGCTGA

1981 AGTCAAGTTT GAAGGTGATA CCCTTGTTAA TAGAATCGAG TTAAAAGGTA TTGATTTTAA

2041 AGAAGATGGA AACATTCTTG GACACAAATT GGAATACAAC TATAACTCAC ACAATGTATA

2101 CATCATGGCA GACAAACAAA AGAATGGAAT CAAAGTTAAC TTCAAAATTA GACACAACAT

2161 TGAAGATGGA AGCGTTCAAC TAGCAGACCA TTATCAACAA AATACTCCAA TTGGCGATGG

2221 CCCTGTCCTT TTACCAGACA ACCATTACCT GTCCACACAA TCTGCCCTTT CGAAAGATCC

2281 CAACGAAAAG AGAGACCACA TGGTCCTTCT TGAGTTTGTA ACAGCTGCTG GGATTACACA

2341 TGGCATGGAT GAACTATACA AATAAggatc cGTTTTTCTT ACTTATATAT TTATACCAAT

2401 TGATTGTATT TATAACTGTA AAAATGTGTA TGTTGTGTGC ATATTTTTTT TTGTGCATGC

2461 ACATGCATGT AAATAGCTAA AATTATGAAC ATTTTATTTT TTGTTCAGAA AAAAAAAACT

2521 TTACACACAT AAAATGGCTA GTATGAATAG CCATATTTTA TATAAATTAA ATCCTATGAA

2581 TTTATGACCA TATTAAAAAT TTAGATATTT ATGGAACATA ATATGTTTGA AACAATAAGA

2641 CAAAATTATT ATTATTATTA TTATTTTTAC TGTTATAATT ATGTTGTCTC TTCAATGATT

2701 CATAAATAGT TGGACTTGAT TTTTAAAATG TTTATAATAT GATTAGCATA GTTAAATAAA

2761 AAAAGTTGAA AAATTAAAAA AAAACATATA AACACAAATG ATGTTTTTTC CTTCAATTTc

2821 ggcgcctgat gcggtatttt ctccttacgc atctgtgcgg tatttcacac cgcatatggt

2881 gcactctcag tacaatctgc tctgatgccg catagttaag ccagccccga cacccgccaa

2941 cacccgctga cgcgccctga cgggcttgtc tgctcccggc atccgcttac agacaagctg

3001 tgaccgtctc cgggagctgc atgtgtcaga ggttttcacc gtcatcaccg aaacgcgcga

3061 gacgaaaggg cctcgtgata cgcctatttt tataggttaa tgtcatgata ataatggttt

3121 cttagacgtc aggtggcact tttcggggaa atgtgcgcgg aacccctatt tgtttatttt

3181 tctaaataca ttcaaatatg tatccgctca tgagacaata accctgataa atgcttcaat

3241 aatattgaaa aaggaagagt atgagtattc aacatttccg tgtcgccctt attccctttt

3301 ttgcggcatt ttgccttcct gtttttgctc acccagaaac gctggtgaaa gtaaaagatg

3361 ctgaagatca gttgggtgca cgagtgggtt acatcgaact ggatctcaac agcggtaaga

3421 tccttgagag ttttcgcccc gaagaacgtt ttccaatgat gagcactttt aaagttctgc

3481 tatgtggcgc ggtattatcc cgtattgacg ccgggcaaga gcaactcggt cgccgcatac

3541 actattctca gaatgacttg gttgagtact caccagtcac agaaaagcat cttacggatg

3601 gcatgacagt aagagaatta tgcagtgctg ccataaccat gagtgataac actgcggcca

3661 acttacttct gacaacgatc ggaggaccga aggagctaac cgcttttttg cacaacatgg

3721 gggatcatgt aactcgcctt gatcgttggg aaccggagct gaatgaagcc ataccaaacg

3781 acgagcgtga caccacgatg cctgtagcaa tggcaacaac gttgcgcaaa ctattaactg

3841 gcgaactact tactctagct tcccggcaac aattaataga ctggatggag gcggataaag

3901 ttgcaggacc acttctgcgc tcggcccttc cggctggctg gtttattgct gataaatctg

3961 gagccggtga gcgtgggtct cgcggtatca ttgcagcact ggggccagat ggtaagccct

4021 cccgtatcgt agttatctac acgacgggga gtcaggcaac tatggatgaa cgaaatagac

4081 agatcgctga gataggtgcc tcactgatta agcattggta actgtcagac caagtttact

4141 catatatact ttagattgat ttaaaacttc atttttaatt taaaaggatc taggtgaaga

4201 tcctttttga taatctcatg accaaaatcc cttaacgtga gttttcgttc cactgagcgt

4261 cagaccccgt agaaaagatc aaaggatctt cttgagatcc tttttttctg cgcgtaatct

4321 gctgcttgca aacaaaaaaa ccaccgctac cagcggtggt ttgtttgccg gatcaagagc

4381 taccaactct ttttccgaag gtaactggct tcagcagagc gcagatacca aatactgtcc

4441 ttctagtgta gccgtagtta ggccaccact tcaagaactc tgtagcaccg cctacatacc

4501 tcgctctgct aatcctgtta ccagtggctg ctgccagtgg cgataagtcg tgtcttaccg

4561 ggttggactc aagacgatag ttaccggata aggcgcagcg gtcgggctga acggggggtt

4621 cgtgcacaca gcccagcttg gagcgaacga cctacaccga actgagatac ctacagcgtg

4681 agcattgaga aagcgccacg cttcccgaag ggagaaaggc ggacaggtat ccggtaagcg

4741 gcagggtcgg aacaggagag cgcacgaggg agcttccagg gggaaacgcc tggtatcttt

4801 atagtcctgt cgggtttcgc cacctctgac ttgagcgtcg atttttgtga tgctcgtcag

4861 gggggcggag cctatggaaa aacgccagca acgcggcctt tttacggttc ctggcctttt

4921 gctggccttt tgctcacatg ttctttcctg cgttatcccc tgattctgtg gataaccgta

4981 ttaccgcctt tgagtgagct gataccgctc gccgcagccg aacgaccgag cgcagcgagt

5041 cagtgagcga ggaagcggaa gagcgcccaa tacgcaaacc gcctctcccc gcgcgttggc

5101 cgattcatta atgcagctgg cacgacaggt ttcccgactg gaaagcgggc agtgagcgca

5161 acgcaattaa tgtgagttag ctcactcatt aggcacccca ggctttacac tttatgcttc

5221 cggctcgtat gttgtgtgga attgtgagcg gataacaatt tcacacagga aacagctatg

5281 accatgatta cgccaagctt gataattcct gcagcccagc ttaattcttt tcgagctctt

5341 tatgcttaag tttacaattt aatattcata ctttaagtat tttttgtagt atcctagata

5401 ttgtgcttta aatgctcacc cctcaaagca ccagtaatat tttcatccac tgaaatacca

5461 ttaaattttc aaaaaaatac tatgcatata atgttataca tataaacata aaacgccatg

5521 taaatcaaaa aatatataaa aatatgtata aaaataaata tgcactaaat ataagctaat

5581 tatgcataaa aattaaagtg ccctttatta actagctagt cgtaattatt tatatttcta

5641 tgttataaaa aaatcctcat ataataatat aattaatata tgtaatgttt tttttatttt

5701 ataattttaa tataaaataa tatgtaaatt aattcaaaaa ataaatataa ttgttgtgaa

5761 acaaaaaacg taattttttc atttgccttc aaaatttaaa tttattttaa tatttcctaa

5821 aatatatata ctttgtgtat aaatatataa aaatatatat ttgcttataa ataaataaaa

5881 aattttataa aacatagggg gatccatggt tggttcgcta aactgcatcg tcgctgtgtc

5941 ccagaacatg ggcatcggca agaacgggga cctgccctgg ccaccgctca ggaacgaatt

6001 tagatatttc cagagaatga ccacaacctc ttcagtagaa ggtaaacaga atctggtgat

6061 tatgggtaag aagacctggt tctccattcc tgagaagaat cgacctttaa agggtagaat

6121 taatttagtt ctcagcagag aactcaagga acctccacaa ggagctcatt ttctttccag

6181 aagtctagat gatgccttaa aacttactga acaaccagaa ttagcaaata aagtagacat

6241 ggtctggata gttggtggca gttctgttta taaggaagcc atgaatcacc caggccatct

6301 taaactattt gtgacaagga tcatgcaaga ctttgaaagt gacacgtttt ttccagaaat

6361 tgatttggag aaatataaac ttctgccaga atacccaggt gttctctctg atgtccagga

6421 ggagaaaggc attaagtaca aatttgaagt atatgagaag aatgattaag gatcccgttt

6481 ttcttactta tatatttata ccaattgatt gtatttataa ctgtaaaaat gtgtatgttg

6541 tgtgcatatt tttttttgtg catgcacatg catgtaaata gctaaaatta tgaacatttt

6601 attttttgtt cagaaaaaaa aaactttaca cacataaaat ggctagtatg aatagccata

6661 ttttatataa attaaatcct atgaatttat gaccatatta aaaatttaga tatttatgga

6721 acataatatg tttgaaacaa taagacaaaa ttattattat tattattatt tttactgtta

6781 taattatgtt gtctcttcaa tgattcataa atagttggac ttgattttta aaatgtttat

6841 aatatgatta gcatagttaa ataaaaaaag ttgaaaaatt aaaaaaaaac atataaacac

6901 aaatgatgtt ttttccttca atttcgatg

//

**pSL1702**

LOCUS	pSL1702	7077 bp	DNA	circular	21-OCT-2024

DEFINITION .

ACCESSION

VERSION

SOURCE	.

ORGANISM .

COMMENT

COMMENT	ApEinfo:methylated:1

FEATURES	Location/Qualifiers

exon	6054..6331

/vntifkey="61"

/locus_tag="hDHFR"

/label="hDHFR"

/ApEinfo_label="hDHFR"

/ApEinfo_fwdcolor="pink"

/ApEinfo_revcolor="pink"

/ApEinfo_graphicformat="arrow_data {{0 1 2 0 0 -1} {} 0}

width 5 offset 0"

exon	6332..6617

/vntifkey="61"

/locus_tag="hDHFR(1)"

/label="hDHFR(1)"

/ApEinfo_label="hDHFR"

/ApEinfo_fwdcolor="pink"

/ApEinfo_revcolor="pink"

/ApEinfo_graphicformat="arrow_data {{0 1 2 0 0 -1} {} 0}

width 5 offset 0"

misc_feature	join(5450..5759,5764..6053)

/vntifkey="21"

/locus_tag="PbEF1a-A\5'UTR"

/label="PbEF1a-A\5'UTR"

/ApEinfo_label="PbEF1a-A\5'UTR"

/ApEinfo_fwdcolor="#eb0214"

/ApEinfo_revcolor="#eb0214"

/ApEinfo_graphicformat="arrow_data {{0 1 2 0 0 -1} {} 0}

width 5 offset 0"

misc_feature	6618..7074

/vntifkey="21"

/locus_tag="PbDHFR/TS\3'UTR"

/label="PbDHFR/TS\3'UTR"

/ApEinfo_label="PbDHFR/TS\3'UTR"

/ApEinfo_fwdcolor="#eb0214"

/ApEinfo_revcolor="#eb0214"

/ApEinfo_graphicformat="arrow_data {{0 1 2 0 0 -1} {} 0}

width 5 offset 0"

misc_feature	1217..1790

/locus_tag="PbEF1a promoter (from pSL0281)"

/label="PbEF1a promoter (from pSL0281)"

/ApEinfo_label="PbEF1a promoter (from pSL0281)"

/ApEinfo_fwdcolor="cyan"

/ApEinfo_revcolor="green"

/ApEinfo_graphicformat="arrow_data {{0 1 2 0 0 -1} {} 0}

width 5 offset 0"

misc_feature	6624..7074

/locus_tag="PbDHFR-TS 3'UTR"

/label="PbDHFR-TS 3'UTR"

/ApEinfo_label="PbDHFR-TS 3'UTR"

/ApEinfo_fwdcolor="cyan"

/ApEinfo_revcolor="green"

/ApEinfo_graphicformat="arrow_data {{0 1 2 0 0 -1} {} 0}

width 5 offset 0"

misc_feature	1835..2513

/locus_tag="GFPmut2"

/label="GFPmut2"

/ApEinfo_label="GFPmut2"

/ApEinfo_fwdcolor="cyan"

/ApEinfo_revcolor="green"

/ApEinfo_graphicformat="arrow_data {{0 1 2 0 0 -1} {} 0}

width 5 offset 0"

misc_feature	6625..7072

/locus_tag="PbDHFR-TS 3'UTR(1)"

/label="PbDHFR-TS 3'UTR(1)"

/ApEinfo_label="PbDHFR-TS 3'UTR"

/ApEinfo_fwdcolor="cyan"

/ApEinfo_revcolor="green"

/ApEinfo_graphicformat="arrow_data {{0 1 2 0 0 -1} {} 0}

width 5 offset 0"

misc_feature	2520..2967

/locus_tag="PbDHFR/TS\3'UTR(1)"

/label="PbDHFR/TS\3'UTR(1)"

/ApEinfo_label="PbDHFR/TS\3'UTR"

/ApEinfo_fwdcolor="#ff0000"

/ApEinfo_revcolor="green"

/ApEinfo_graphicformat="arrow_data {{0 1 2 0 0 -1} {} 0}

width 5 offset 0"

rep_origin	4364..5046

/locus_tag="ColE1 origin"

/label="ColE1 origin"

/ApEinfo_label="ColE1 origin"

/ApEinfo_fwdcolor="gray50"

/ApEinfo_revcolor="gray50"

/ApEinfo_graphicformat="arrow_data {{0 1 2 0 0 -1} {} 0}

width 5 offset 0"

CDS	3607..4266

/locus_tag="AmpR"

/label="AmpR"

/ApEinfo_label="AmpR"

/ApEinfo_fwdcolor="yellow"

/ApEinfo_revcolor="yellow"

/ApEinfo_graphicformat="arrow_data {{0 1 2 0 0 -1} {} 0}

width 5 offset 0"

misc_feature	join(5450..5759,5764..6048)

/locus_tag="PbEF1a 5'UTR"

/label="PbEF1a 5'UTR"

/ApEinfo_label="PbEF1a 5'UTR"

/ApEinfo_fwdcolor="#804040"

/ApEinfo_revcolor="#804040"

/ApEinfo_graphicformat="arrow_data {{0 1 2 0 0 -1} {} 0}

width 5 offset 0"

misc_feature	1..34

/locus_tag="loxP Site"

/label="loxP Site"

/ApEinfo_label="loxP Site"

/ApEinfo_fwdcolor="#ffff00"

/ApEinfo_revcolor="green"

/ApEinfo_graphicformat="arrow_data {{0 1 2 0 0 -1} {} 0}

width 5 offset 0"

misc_feature	1177..1215

/locus_tag="loxP Site(1)"

/label="loxP Site(1)"

/ApEinfo_label="loxP Site"

/ApEinfo_fwdcolor="#ffff00"

/ApEinfo_revcolor="green"

/ApEinfo_graphicformat="arrow_data {{0 1 2 0 0 -1} {} 0}

width 5 offset 0"

misc_feature	635..1170

/locus_tag="28 Del 5' HR"

/label="28 Del 5' HR"

/ApEinfo_label="28 Del 5' HR"

/ApEinfo_fwdcolor="#0080ff"

/ApEinfo_revcolor="green"

/ApEinfo_graphicformat="arrow_data {{0 0.5 0 1 2 0 0 -1 0

-0.5} {} 0} width 5 offset 0"

misc_feature	41..614

/locus_tag="3' HR"

/label="3' HR"

/ApEinfo_label="3' HR"

/ApEinfo_fwdcolor="#00ff00"

/ApEinfo_revcolor="green"

/ApEinfo_graphicformat="arrow_data {{0 0.5 0 1 2 0 0 -1 0

-0.5} {} 0} width 5 offset 0"

ORIGIN

1 ATAACTTCGT ATAGCATACA TTATACGAAG TTATggtacc GTTCTTCTTT GACTTACTGC

61 TGATTTAAAA GCGAAACTAT ATGTATAATA CTTGTTATAT ACAATTTATA CAATTAATCC

121 TATTGCCATT TGGAGGAGGA GTCATGCCTC CTTTTGGTTC TTGTAATTAA AAACAAAGAT

181 GAATTAATAC AAACTGTAGA CGACTTTTAG GCCTCGGGGT GCTGTAAACA TGAAAGTAAA

241 CTTAGTTTTA CGATCTGTTA AGGCTTATCC TCTGTGGTAA AGTATTTTTT TAAACTTAAA

301 TAGTATTTTT TTAAGCTTAA ATATAAATAA AAAAAATACA TATTATAATA TATTATAAAT

361 TTATATTGCA TAAGTTTGCA AATATAAATT TATGATATAT TATATATATT AACAATTTTT

421 AATATCAATT TTTTACATTT TTGATTATGC AATTATTTGT ATAAAAAAAA TATATATTAT

481 AATATATTAT AAATTTATAT TGCAAACTTA TGCAGTATAA ATTTATGATA TATTATATAT

541 ATTAACAATT TTTAATATCA ATTTTTTACA TTATTTATGC TCGTGATTGC ATGGATTTCC

601 TACTTAGGGt gtcggatcgt cgacgatcac cggtGTGGTT GGAAACTCAT AGGGTTAGGA

661 CACCAGTGTT ATGTTACATT TTTACAATGG ACGTGTCGGA AACACATATA GGTGTTAGGG

721 GCTACATTGC TTAAAGCATA TTTGCACTTT GTATGTTGAA GGCAGTATAA TGTTACAACA

781 TATCAATGCT GCGAAGCATA ATGGTGTAAA GTGTTATGTT ATCTGTGAAC TTTGAGATAT

841 TTTAATTAGC ATATATTGCG CGGTGTAGTC GTAAGCTAAT AAAATGACCT GAACAGTCAT

901 GTTTAAAGAT AAAGACAATA GAAATACATT AACAACTTCA AGTAACTAAC AACTTCTAAG

961 TTAAACAACT TAAACTGGTT CACCCAAACC AGTTAACTAC TTAACTGGAA AAATACTCAT

1021 ATAGTTAGTT GTATAAAATC CTATATAAAT GTGAGCAAAC CTGTTGTAAT GACAAATAAG

1081 CTATAAATAG TTAGATGGAG ATGGATAAAT GGCTAATTAC ATGGAAACTA TGAGAGGTGA

1141 TTGCATATTG ACATATGGAA TAGGAGAGCG ctcgagATAA CTTCGTATAG CATACATTAT

1201 ACGAAGTTAT gtcgagAGCT TAATTCTTTT CGAGCTCTTT ATGCTTAAGT TTACAATTTA

1261 ATATTCATAC TTTAAGTATT TTTTGTAGTA TCCTAGATAT TGTGCTTTAA ATGCTCACCC

1321 CTCAAAGCAC CAGTAATATT TTCATCCACT GAAATACCAT TAAATTTTCA AAAAAATACT

1381 ATGCATATAA TGTTATACAT ATAAACATAA AACGCCATGT AAATCAAAAA ATATATAAAA

1441 ATATGTATAA AAATAAATAT GCACTAAATA TAAGCTAATT ATGCATAAAA ATTAAAGTGC

1501 CCTTTATTAA CTAGctagTC GTAATTATTT ATATTTCTAT GTTATAAAAA AATCCTCATA

1561 TAATAATATA ATTAATATAT GTAATGTTTT TTTTATTTTA TAATTTTAAT ATAAAATAAT

1621 ATGTAAATTA ATTCAAAAAA TAAATATAAT TGTTGTGAAA CAAAAAACGT AATTTTTTCA

1681 TTTGCCTTCA AAATTTAAAT TTATTTTAAT ATTTCCTAAA ATATATATAC TTTGTGTATA

1741 AATATATAAA AATATATATT TGCTTATAAA TAAATAAAAA TTTTATAAAA atgactagta

1801 gtaaaggaga agaacttttc actggagttg tcccAATTCT TGTTGAATTA GATGGTGATG

1861 TTAATGGGCA CAAATTTTCT GTCAGTGGAG AGGGTGAAGG TGATGCAACA TACGGAAAAC

1921 TTACCCTTAA ATTTATTTGC ACTACTGGAA AACTACCTGT TCCATGGCCA ACACTTGTCA

1981 CTACTTTCGC GTATGGTCTT CAATGCTTTG CGAGATACCC AGATCATATG AAACAGCATG

2041 ACTTTTTCAA GAGTGCCATG CCCGAAGGTT ATGTACAGGA AAGAACTATA TTTTTCAAAG

2101 ATGACGGGAA CTACAAGACA CGTGCTGAAG TCAAGTTTGA AGGTGATACC CTTGTTAATA

2161 GAATCGAGTT AAAAGGTATT GATTTTAAAG AAGATGGAAA CATTCTTGGA CACAAATTGG

2221 AATACAACTA TAACTCACAC AATGTATACA TCATGGCAGA CAAACAAAAG AATGGAATCA

2281 AAGTTAACTT CAAAATTAGA CACAACATTG AAGATGGAAG CGTTCAACTA GCAGACCATT

2341 ATCAACAAAA TACTCCAATT GGCGATGGCC CTGTCCTTTT ACCAGACAAC CATTACCTGT

2401 CCACACAATC TGCCCTTTCG AAAGATCCCA ACGAAAAGAG AGACCACATG GTCCTTCTTG

2461 AGTTTGTAAC AGCTGCTGGG ATTACACATG GCATGGATGA ACTATACAAA TAAggatccG

2521 TTTTTCTTAC TTATATATTT ATACCAATTG ATTGTATTTA TAACTGTAAA AATGTGTATG

2581 TTGTGTGCAT ATTTTTTTTT GTGCATGCAC ATGCATGTAA ATAGCTAAAA TTATGAACAT

2641 TTTATTTTTT GTTCAGAAAA AAAAAACTTT ACACACATAA AATGGCTAGT ATGAATAGCC

2701 ATATTTTATA TAAATTAAAT CCTATGAATT TATGACCATA TTAAAAATTT AGATATTTAT

2761 GGAACATAAT ATGTTTGAAA CAATAAGACA AAATTATTAT TATTATTATT ATTTTTACTG

2821 TTATAATTAT GTTGTCTCTT CAATGATTCA TAAATAGTTG GACTTGATTT TTAAAATGTT

2881 TATAATATGA TTAGCATAGT TAAATAAAAA AAGTTGAAAA ATTAAAAAAA AACATATAAA

2941 CACAAATGAT GTTTTTTCCT TCAATTTcgg cgcctgatgc ggtattttct ccttacgcat

3001 ctgtgcggta tttcacaccg catatggtgc actctcagta caatctgctc tgatgccgca

3061 tagttaagcc agccccgaca cccgccaaca cccgctgacg cgccctgacg ggcttgtctg

3121 ctcccggcat ccgcttacag acaagctgtg accgtctccg ggagctgcat gtgtcagagg

3181 ttttcaccgt catcaccgaa acgcgcgaga cgaaagggcc tcgtgatacg cctattttta

3241 taggttaatg tcatgataat aatggtttct tagacgtcag gtggcacttt tcggggaaat

3301 gtgcgcggaa cccctatttg tttatttttc taaatacatt caaatatgta tccgctcatg

3361 agacaataac cctgataaat gcttcaataa tattgaaaaa ggaagagtat gagtattcaa

3421 catttccgtg tcgcccttat tccctttttt gcggcatttt gccttcctgt ttttgctcac

3481 ccagaaacgc tggtgaaagt aaaagatgct gaagatcagt tgggtgcacg agtgggttac

3541 atcgaactgg atctcaacag cggtaagatc cttgagagtt ttcgccccga agaacgtttt

3601 ccaatgatga gcacttttaa agttctgcta tgtggcgcgg tattatcccg tattgacgcc

3661 gggcaagagc aactcggtcg ccgcatacac tattctcaga atgacttggt tgagtactca

3721 ccagtcacag aaaagcatct tacggatggc atgacagtaa gagaattatg cagtgctgcc

3781 ataaccatga gtgataacac tgcggccaac ttacttctga caacgatcgg aggaccgaag

3841 gagctaaccg cttttttgca caacatgggg gatcatgtaa ctcgccttga tcgttgggaa

3901 ccggagctga atgaagccat accaaacgac gagcgtgaca ccacgatgcc tgtagcaatg

3961 gcaacaacgt tgcgcaaact attaactggc gaactactta ctctagcttc ccggcaacaa

4021 ttaatagact ggatggaggc ggataaagtt gcaggaccac ttctgcgctc ggcccttccg

4081 gctggctggt ttattgctga taaatctgga gccggtgagc gtgggtctcg cggtatcatt

4141 gcagcactgg ggccagatgg taagccctcc cgtatcgtag ttatctacac gacggggagt

4201 caggcaacta tggatgaacg aaatagacag atcgctgaga taggtgcctc actgattaag

4261 cattggtaac tgtcagacca agtttactca tatatacttt agattgattt aaaacttcat

4321 ttttaattta aaaggatcta ggtgaagatc ctttttgata atctcatgac caaaatccct

4381 taacgtgagt tttcgttcca ctgagcgtca gaccccgtag aaaagatcaa aggatcttct

4441 tgagatcctt tttttctgcg cgtaatctgc tgcttgcaaa caaaaaaacc accgctacca

4501 gcggtggttt gtttgccgga tcaagagcta ccaactcttt ttccgaaggt aactggcttc

4561 agcagagcgc agataccaaa tactgtcctt ctagtgtagc cgtagttagg ccaccacttc

4621 aagaactctg tagcaccgcc tacatacctc gctctgctaa tcctgttacc agtggctgct

4681 gccagtggcg ataagtcgtg tcttaccggg ttggactcaa gacgatagtt accggataag

4741 gcgcagcggt cgggctgaac ggggggttcg tgcacacagc ccagcttgga gcgaacgacc

4801 tacaccgaac tgagatacct acagcgtgag cattgagaaa gcgccacgct tcccgaaggg

4861 agaaaggcgg acaggtatcc ggtaagcggc agggtcggaa caggagagcg cacgagggag

4921 cttccagggg gaaacgcctg gtatctttat agtcctgtcg ggtttcgcca cctctgactt

4981 gagcgtcgat ttttgtgatg ctcgtcaggg gggcggagcc tatggaaaaa cgccagcaac

5041 gcggcctttt tacggttcct ggccttttgc tggccttttg ctcacatgtt ctttcctgcg

5101 ttatcccctg attctgtgga taaccgtatt accgcctttg agtgagctga taccgctcgc

5161 cgcagccgaa cgaccgagcg cagcgagtca gtgagcgagg aagcggaaga gcgcccaata

5221 cgcaaaccgc ctctccccgc gcgttggccg attcattaat gcagctggca cgacaggttt

5281 cccgactgga aagcgggcag tgagcgcaac gcaattaatg tgagttagct cactcattag

5341 gcaccccagg ctttacactt tatgcttccg gctcgtatgt tgtgtggaat tgtgagcgga

5401 taacaatttc acacaggaaa cagctatgac catgattacg ccaagcttga taattcctgc

5461 agcccagctt aattcttttc gagctcttta tgcttaagtt tacaatttaa tattcatact

5521 ttaagtattt tttgtagtat cctagatatt gtgctttaaa tgctcacccc tcaaagcacc

5581 agtaatattt tcatccactg aaataccatt aaattttcaa aaaaatacta tgcatataat

5641 gttatacata taaacataaa acgccatgta aatcaaaaaa tatataaaaa tatgtataaa

5701 aataaatatg cactaaatat aagctaatta tgcataaaaa ttaaagtgcc ctttattaac

5761 tagctagtcg taattattta tatttctatg ttataaaaaa atcctcatat aataatataa

5821 ttaatatatg taatgttttt tttattttat aattttaata taaaataata tgtaaattaa

5881 ttcaaaaaat aaatataatt gttgtgaaac aaaaaacgta attttttcat ttgccttcaa

5941 aatttaaatt tattttaata tttcctaaaa tatatatact ttgtgtataa atatataaaa

6001 atatatattt gcttataaat aaataaaaaa ttttataaaa cataggggga tccatggttg

6061 gttcgctaaa ctgcatcgtc gctgtgtccc agaacatggg catcggcaag aacggggacc

6121 tgccctggcc accgctcagg aacgaattta gatatttcca gagaatgacc acaacctctt

6181 cagtagaagg taaacagaat ctggtgatta tgggtaagaa gacctggttc tccattcctg

6241 agaagaatcg acctttaaag ggtagaatta atttagttct cagcagagaa ctcaaggaac

6301 ctccacaagg agctcatttt ctttccagaa gtctagatga tgccttaaaa cttactgaac

6361 aaccagaatt agcaaataaa gtagacatgg tctggatagt tggtggcagt tctgtttata

6421 aggaagccat gaatcaccca ggccatctta aactatttgt gacaaggatc atgcaagact

6481 ttgaaagtga cacgtttttt ccagaaattg atttggagaa atataaactt ctgccagaat

6541 acccaggtgt tctctctgat gtccaggagg agaaaggcat taagtacaaa tttgaagtat

6601 atgagaagaa tgattaagga tcccgttttt cttacttata tatttatacc aattgattgt

6661 atttataact gtaaaaatgt gtatgttgtg tgcatatttt tttttgtgca tgcacatgca

6721 tgtaaatagc taaaattatg aacattttat tttttgttca gaaaaaaaaa actttacaca

6781 cataaaatgg ctagtatgaa tagccatatt ttatataaat taaatcctat gaatttatga

6841 ccatattaaa aatttagata tttatggaac ataatatgtt tgaaacaata agacaaaatt

6901 attattatta ttattatttt tactgttata attatgttgt ctcttcaatg attcataaat

6961 agttggactt gatttttaaa atgtttataa tatgattagc atagttaaat aaaaaaagtt

7021 gaaaaattaa aaaaaaacat ataaacacaa atgatgtttt ttccttcaat ttcgatg

//

**pSL1754**

LOCUS	pSL1754	10658 bp	DNA	circular	21-OCT-2024

DEFINITION .

ACCESSION

VERSION

SOURCE	.

ORGANISM .

COMMENT

COMMENT	ApEinfo:methylated:1

FEATURES	Location/Qualifiers

exon	6990..7847

/vntifkey="61"

/locus_tag="AMP"

/label="AMP"

/ApEinfo_label="AMP"

/ApEinfo_fwdcolor="pink"

/ApEinfo_revcolor="pink"

/ApEinfo_graphicformat="arrow_data {{0 1 2 0 0 -1} {} 0}

width 5 offset 0"

rep_origin	7945..8627

/locus_tag="ColE1 origin"

/label="ColE1 origin"

/ApEinfo_label="ColE1 origin"

/ApEinfo_fwdcolor="gray50"

/ApEinfo_revcolor="gray50"

/ApEinfo_graphicformat="arrow_data {{0 1 2 0 0 -1} {} 0}

width 5 offset 0"

exon	9635..10198

/vntifkey="61"

/locus_tag="hDHFR"

/label="hDHFR"

/ApEinfo_label="hDHFR"

/ApEinfo_fwdcolor="pink"

/ApEinfo_revcolor="pink"

/ApEinfo_graphicformat="arrow_data {{0 1 2 0 0 -1} {} 0}

width 5 offset 0"

misc_feature	join(9031..9340,9345..9634)

/vntifkey="21"

/locus_tag="PbEF1a-A\5'UTR"

/label="PbEF1a-A\5'UTR"

/ApEinfo_label="PbEF1a-A\5'UTR"

/ApEinfo_fwdcolor="#eb0214"

/ApEinfo_revcolor="#eb0214"

/ApEinfo_graphicformat="arrow_data {{0 1 2 0 0 -1} {} 0}

width 5 offset 0"

misc_feature	4799..5372

/locus_tag="PbEF1a promoter (from pSL0281)"

/label="PbEF1a promoter (from pSL0281)"

/ApEinfo_label="PbEF1a promoter (from pSL0281)"

/ApEinfo_fwdcolor="cyan"

/ApEinfo_revcolor="green"

/ApEinfo_graphicformat="arrow_data {{0 1 2 0 0 -1} {} 0}

width 5 offset 0"

misc_feature	5417..6095

/locus_tag="GFPmut2"

/label="GFPmut2"

/ApEinfo_label="GFPmut2"

/ApEinfo_fwdcolor="cyan"

/ApEinfo_revcolor="green"

/ApEinfo_graphicformat="arrow_data {{0 1 2 0 0 -1} {} 0}

width 5 offset 0"

misc_feature	10199..10655

/vntifkey="21"

/locus_tag="PbDHFR/TS\3'UTR"

/label="PbDHFR/TS\3'UTR"

/ApEinfo_label="PbDHFR/TS\3'UTR"

/ApEinfo_fwdcolor="#eb0214"

/ApEinfo_revcolor="#eb0214"

/ApEinfo_graphicformat="arrow_data {{0 1 2 0 0 -1} {} 0}

width 5 offset 0"

misc_feature	6102..6548

/locus_tag="PbDHFR/TS\3'UTR(1)"

/label="PbDHFR/TS\3'UTR(1)"

/ApEinfo_label="PbDHFR/TS\3'UTR"

/ApEinfo_fwdcolor="#ff0000"

/ApEinfo_revcolor="green"

/ApEinfo_graphicformat="arrow_data {{0 1 2 0 0 -1} {} 0}

width 5 offset 0"

CDS	7188..7847

/locus_tag="AmpR"

/label="AmpR"

/ApEinfo_label="AmpR"

/ApEinfo_fwdcolor="yellow"

/ApEinfo_revcolor="yellow"

/ApEinfo_graphicformat="arrow_data {{0 1 2 0 0 -1} {} 0}

width 5 offset 0"

misc_feature	10205..10655

/locus_tag="PbDHFR-TS 3'UTR"

/label="PbDHFR-TS 3'UTR"

/ApEinfo_label="PbDHFR-TS 3'UTR"

/ApEinfo_fwdcolor="cyan"

/ApEinfo_revcolor="green"

/ApEinfo_graphicformat="arrow_data {{0 1 2 0 0 -1} {} 0}

width 5 offset 0"

misc_feature	join(9031..9340,9345..9629)

/locus_tag="PbEF1a 5'UTR"

/label="PbEF1a 5'UTR"

/ApEinfo_label="PbEF1a 5'UTR"

/ApEinfo_fwdcolor="#804040"

/ApEinfo_revcolor="#804040"

/ApEinfo_graphicformat="arrow_data {{0 1 2 0 0 -1} {} 0}

width 5 offset 0"

misc_feature	10206..10653

/locus_tag="PbDHFR-TS 3'UTR(1)"

/label="PbDHFR-TS 3'UTR(1)"

/ApEinfo_label="PbDHFR-TS 3'UTR"

/ApEinfo_fwdcolor="cyan"

/ApEinfo_revcolor="green"

/ApEinfo_graphicformat="arrow_data {{0 1 2 0 0 -1} {} 0}

width 5 offset 0"

misc_feature	7..535

/locus_tag="DiCre 3' HR(1)"

/label="DiCre 3' HR(1)"

/ApEinfo_label="DiCre 3' HR"

/ApEinfo_fwdcolor="#ffff80"

/ApEinfo_revcolor="green"

/ApEinfo_graphicformat="arrow_data {{0 0.5 0 1 2 0 0 -1 0

-0.5} {} 0} width 5 offset 0"

misc_feature	721..1176

/locus_tag="DiCre 5'HR "

/label="DiCre 5'HR "

/ApEinfo_label="DiCre 5'HR "

/ApEinfo_fwdcolor="#804000"

/ApEinfo_revcolor="green"

/ApEinfo_graphicformat="arrow_data {{0 0.5 0 1 2 0 0 -1 0

-0.5} {} 0} width 5 offset 0"

misc_feature	1183..4783

/locus_tag="S2 Promoter-ETS-18S-ITS1-5.8S"

/label="S2 Promoter-ETS-18S-ITS1-5.8S"

/ApEinfo_label="S2 Promoter-ETS-18S-ITS1-5.8S"

/ApEinfo_fwdcolor="#800080"

/ApEinfo_revcolor="green"

/ApEinfo_graphicformat="arrow_data {{0 0.5 0 1 2 0 0 -1 0

-0.5} {} 0} width 5 offset 0"

ORIGIN

1 ggtaccCTTA ACTAACGACA AATTGTGATG ATCTTACTGA AATTAAAATA TTACAAAATA

61 AGTTGCAAAT ATTTGTCTTT GCATCTATAA AAATATAATT TTATTTTGCT TTTCTATGAT

121 TCACTAATAT ATCACAATAG ATGTATTCAA AGCAAAATAA ATAAGCATTA CATGTGTAAT

181 ATATTTGCAA CACTTTCTCA TAAAATAAAA ATTAATAACT ACATGAATAT AACGAATATG

241 ATTTTATTTT TCTTTTTTTT ATCCAAAGAT ATATAGCGAT ATCATCTATA TTATTTTATT

301 AGTTTTTTTC GAAAATTATA AAACAAACAT AACTGTCTAT TATATTATTT ATCTCATAAT

361 ATATATATAT ACAAAATAAA ATACATTTTA AAAAAATATA AATTTAATTT TTTTATATTT

421 TCAAAAGTAT AAAAATAACC TAAAAACTTA TTAATTTTTT ACATAAAAAA TATAGACAAG

481 TATTAAAAAA AAAAAAAAAA AAAAACGACA AATCGCACAT TTAAAAAAAG ATACACATGT

541 GCAATGTAGT TATATATAAT TTTAAAGATA ATTTATTTAA TAAAACTGGT TTTAAATTAT

601 TAATTATAAA ATGGAAAAAT AAATAAAGAG TGTAAACAAT TATATATATT TTTTTTCTCC

661 CATTTCTCAT TATCTTTATA TAGTGCATCT TTCTACgatc CCCGGGgatc AGGCCTgatc

721 CGTGTTTTGC GATTTCTTTG GGTTCGTGTT TTGCGATTTG TTTGGGTTCG TGTTTTGTGA

781 TTTCTTTGCG ATTTTTGTGT GTTTAGTGAT TTTTGTGTGT GTCGATTGTG CGTTTTTTAT

841 TATATTATTG TTATCATATT TGATTTACCT TTTGATATTA GGACATATAT ATATGCATAT

901 ATTTTTTTGT ATTTCCTGAT TTGTTTGTGA TCATAATTTT CGATAAAATG AGTTTTTATT

961 ATTGATGCTT TAAACATTCC AGTTGGTTCC AGATCTCTTT TTTCTCGTCT CGTTTAGAAA

1021 TTATTTCATT TTTGCATATT CTTGCTCATA TGCCAATGTC TTACATGTTT TTCGTAACAA

1081 CTTTCACGCT AGCTACTTAT TTCACAGTGA TACTTCACTA GTATAAGTTG CATCTAATTT

1141 ATATTGGCAA CTCTAATAGA ACCCCTAAAT TTATGGgggc ccGCATTTCC GAATTTCCGT

1201 TTGGTATTTA TTGTGTGTAT AAGTATTATA CAAATATGAA ATATTGTGAA ATTAATATTA

1261 TTTTAAATAT ATTATAACAA GAGCTGTTAT TGTATGTTTT TTCTATGTAA TAATATTATT

1321 TACGATGTTT TAAAATACTT CTTTCAATTT AGTTTTCTAA TATTGGAGAG TATATAGTGA

1381 AACCAGACCA ATATTATATT TATTTGGATA ATTAATTGTG CGTGCAATAT TTGAATAATA

1441 TATAATAAAT ATGTTTTAGG ATATATTATA TCATTTAATT AAAAAAATGT GTGTGTATGT

1501 GTGTTATGTA TAGTTATAAA AAACAACATC GTGGTGTTGT TTATGTATAG CAATATAAAA

1561 CAAAACATTC CGTAATATAG AATAAAAAAT ATAATACATA ATAATGTATA TTATTTAAAC

1621 AAAAAAATAA TTTTGTAACC CTTTATTAAT ATGGACCACA TTTGTTAAAA AATTATTTAA

1681 ATATAAATAA ATTTTTAATA AATAAAATCG TAAAGAATTG TAAAAGAGTT TCAATATTAT

1741 CGGGACCTTT TATGATATTA ATGTATAAAT CCGTAATTTG TATACTATAT AAATTAGTTT

1801 TTGAAATTAG CACTATAGCA AAAATTATAT TGTCTTTTAT GTGTGTGTAT CGAATATTAC

1861 CACCAAAAAA ATAATCGATG ATATATTTTT ATGCTAGGTG CATTTATTTT TTTTATTATA

1921 CTATAAATAA AATATAGTAT GATAAAGAAA AAATAGCAAA TAAAAATCAC ATAGGAACGG

1981 CTTTTAAGAG ATATAAAATC CCTTAGGAGT GGTTTTCTCT TTTGTGTTGT ATAACATATA

2041 GTAATGTTAT TAATATGAGA TATCATATGT ATATATTATA TATAATATAA TTGATATACA

2101 TAATTTTTTT TATGTATATT AAAATTATTA AATATAATAT TTTGATATTT CATATATTAT

2161 TAATATTAAT AGTAACCTGG TTGATCTTGC CAGTAGTCAT ATGCTTGTCT CAAAGATTAA

2221 GCCATGCAAG TGAAAGTATA TACACAATTA TTGTAGAAAC TGCGAACGGC TCATTAAAAC

2281 AGTTATAATC TACTTGACAT TTTATTATAA GGATAACTAC GGAAAAGCTG TAGCTAATAC

2341 TTGTAAGTAC ATTTACTCTA AGGAGTAATT AGTACGTATT TGTTAAGACC CCTAAGAAAA

2401 AATGATATTA AAGGAATTAT AACAAAGAAG CGACACATAA TGTAATATTC AGTGTGTATC

2461 AATCGAGTTT CTGACCTATC AGCTTTTGAT GTTAGGGTAT TGCCCTAACA TGGCTTTGAC

2521 GGGTAACGGG GAATTAGAGT TCGATTCCGG AGAGGGAGCC TGAGAAATAG CTACCACATC

2581 TAAGGAAGGC AGCAGGCGCG TAAATTACCC AATTCTAAAT AAGAGAGGTA GTGACAAGAA

2641 ATAACAATAT AAGGCCAAAT TTTGGTTTTA TAATTGGAAT GATGGGAACT TAAAACCTTC

2701 CCAAAAATCA ATTGGAGGGC AAGTCTGGTG CCAGCAGCCG CGGTAATTCC AGCTCCAATA

2761 GCGTATATTA AAATTGTTGC AGTTAAAACG CTCGTAGTTT AATTTCAAGG GTAATATTAT

2821 TTTAAGTACT CACTTGGAAA GAATGTGACT TCGGTCATAG TTTGTATCCT AGATGGCGTT

2881 CTTTTAATTA CAGGCCCTTT GAGAACTCAT TAATTTATAA CTGAGTTTCT CGTTACTTTG

2941 AGTAAATTAG AGTGTTTAAA GCATACAGAT AAAACGTATT TTACTGTGTT TGAATACTAT

3001 AGCATGGAAT AACAACATTG AATAAGTCAA AAATTTTTGA AAAATTTTTC TTATTTTGGC

3061 TTAGATACAG TTAATAGGAG TAGCTTGGGG GCATTTGTAT TCAGATGTCA GAGGTGAAAT

3121 TCTTAGATTT TCTGGAGACA AACAACTGCG AAAGCATTTG CCTAAAATAC TTCCATTAAT

3181 CAAGAACGAA AGTTAAGGGA GTGAAGACGA TCAGATACCG TCGTAATCTT AACCATAAAC

3241 TATGCCGACT AAGTGTTGGA TGAAAATTTA TAAATAAAGC TATCTTCTTT AAAAGGAGTA

3301 GTTTTTTAGA TGCTTCCTTC AGTACCTTAT GAGAAATCAA AGTCTTTGGG TTCTGGGGCG

3361 AGTATTCGCG CAAGCGAGAA AGTTAAAAGA ATTGACGGAA GGGCACCACC AGGCGTGGAG

3421 CTTGCGGCTT AATTTGACTC AACACGGGGA ACCTCACTAG TTTAAGACAA GAGTAGGATT

3481 GACAGATTAA TAGCTCTTTC TTGATTTCTT GGATGGTGAT GCATGGCCGT TTTTAGTTCG

3541 TGAATATGAT TTGTCTGGTT AATTCCGATA ACGAACGAGA TCTTAACCTG CTAATTAGCG

3601 GTAGTTACGT GATATTCTTC GAAGTGGAAT TACTATAACG TTTCCAAAGT TATGTTGCAT

3661 CATAATCAAA ATGGATTTAC CTTTTGTTTT ATTGTAGCAT ATTCGGTGGA TTTCGTTGGA

3721 TTCTTTCCCT AGTAAGGATG TATCTACTTT ATTTAAAGCT TCTTAGAGGA ACGATGTGTG

3781 TCTAACACAA GGAAGTTTAA GGCAACAACA GGTCTGTGAT GTCCTTAGAT ATACTAGGCT

3841 GCACGCGTGC TACACTGATA TGTAAAACGA GTATTTAAAA TTATATCTGT ATAATAGATA

3901 ACTTAATTTC TATATATATC AGCATATATT TTTCCTACAC TGAAATAGTG AAGGTAATCT

3961 TTATCAATAC ATATCGTGAT GGGGATAGAT TATTGCAATT ATTAATCTTG AACGAGGAAT

4021 GCCTAGTAAG CATGATTCAT CAGATTGTGC TGACTACGTC CCTGCCCTTT GTACACACCG

4081 CCCGTCGCTC CTACCGATTG AAAGATATGA TGAATTGTTT GGACAAGAAA ATCGAAATTT

4141 TATTTTTATT TTTTTGGAAG GACCGTAAAT CCTATCTTTT AAAGGAAGGA GAAGTCGTAA

4201 CAAGGTTTCC GTAGGTGAAC CTGCGGAAGG ATCATTTTTA CACCCGGATA CGGTACCTTA

4261 GTTCGTGTGC TTTATTATTC AATATTTATA AAAATGTGTA ACAGAGGTGT AGATTGTGAG

4321 GGTATATGTT TTGTATTAAT ATTCGATGTC ATCATATAAA AGAAATATAT TTTAATATAT

4381 CAAGATATGT ATACCAGTCG ACGTTAATAC AGAATGCTAA GTATATTCTC ATAAAATATA

4441 TAGTATGTGC TCATTTCTTC ATAATGAGGA TATAAAGTGT GTGTATTAAG GTCCGTATCC

4501 ACTAATTAAC CATATATAGC GTACGATACC TCTGCTTGGT ATATTTGGAG TTGATTGTGT

4561 CTTTTCATTT TTTCCGAAAG GTTAATATGG AAATGATATG ATTGATTTCG AACGTATCAC

4621 TCATGTGTCC CATGCATATG TGACAATAAA CAACTATCAA TTTTTGCATG ATAATGTGTT

4681 TGTTTATTTT GTTTGAAACG CAATATTTCG ATAAAGCAAG TTCAAGGAAA ATATAAAAAA

4741 TTTAAGCACA ATCTTAACGA TGGATGTCTT GGTTCCTACA GCGgcggccg cTctcgagAG

4801 CTTAATTCTT TTCGAGCTCT TTATGCTTAA GTTTACAATT TAATATTCAT ACTTTAAGTA

4861 TTTTTTGTAG TATCCTAGAT ATTGTGCTTT AAATGCTCAC CCCTCAAAGC ACCAGTAATA

4921 TTTTCATCCA CTGAAATACC ATTAAATTTT CAAAAAAATA CTATGCATAT AATGTTATAC

4981 ATATAAACAT AAAACGCCAT GTAAATCAAA AAATATATAA AAATATGTAT AAAAATAAAT

5041 ATGCACTAAA TATAAGCTAA TTATGCATAA AAATTAAAGT GCCCTTTATT AACTAGctag

5101 TCGTAATTAT TTATATTTCT ATGTTATAAA AAAATCCTCA TATAATAATA TAATTAATAT

5161 ATGTAATGTT TTTTTTATTT TATAATTTTA ATATAAAATA ATATGTAAAT TAATTCAAAA

5221 AATAAATATA ATTGTTGTGA AACAAAAAAC GTAATTTTTT CATTTGCCTT CAAAATTTAA

5281 ATTTATTTTA ATATTTCCTA AAATATATAT ACTTTGTGTA TAAATATATA AAAATATATA

5341 TTTGCTTATA AATAAATAAA AATTTTATAA AAatgactag tagtaaagga gaagaacttt

5401 tcactggagt tgtcccAATT CTTGTTGAAT TAGATGGTGA TGTTAATGGG CACAAATTTT

5461 CTGTCAGTGG AGAGGGTGAA GGTGATGCAA CATACGGAAA ACTTACCCTT AAATTTATTT

5521 GCACTACTGG AAAACTACCT GTTCCATGGC CAACACTTGT CACTACTTTC GCGTATGGTC

5581 TTCAATGCTT TGCGAGATAC CCAGATCATA TGAAACAGCA TGACTTTTTC AAGAGTGCCA

5641 TGCCCGAAGG TTATGTACAG GAAAGAACTA TATTTTTCAA AGATGACGGG AACTACAAGA

5701 CACGTGCTGA AGTCAAGTTT GAAGGTGATA CCCTTGTTAA TAGAATCGAG TTAAAAGGTA

5761 TTGATTTTAA AGAAGATGGA AACATTCTTG GACACAAATT GGAATACAAC TATAACTCAC

5821 ACAATGTATA CATCATGGCA GACAAACAAA AGAATGGAAT CAAAGTTAAC TTCAAAATTA

5881 GACACAACAT TGAAGATGGA AGCGTTCAAC TAGCAGACCA TTATCAACAA AATACTCCAA

5941 TTGGCGATGG CCCTGTCCTT TTACCAGACA ACCATTACCT GTCCACACAA TCTGCCCTTT

6001 CGAAAGATCC CAACGAAAAG AGAGACCACA TGGTCCTTCT TGAGTTTGTA ACAGCTGCTG

6061 GGATTACACA TGGCATGGAT GAACTATACA AATAAggatc cGTTTTTCTT ACTTATATAT

6121 TTATACCAAT TGATTGTATT TATAACTGTA AAAATGTGTA TGTTGTGTGC ATATTTTTTT

6181 TTGTGCATGC ACATGCATGT AAATAGCTAA AATTATGAAC ATTTTATTTT TTGTTCAGAA

6241 AAAAAAACTT TACACACATA AAATGGCTAG TATGAATAGC CATATTTTAT ATAAATTAAA

6301 TCCTATGAAT TTATGACCAT ATTAAAAATT TAGATATTTA TGGAACATAA TATGTTTGAA

6361 ACAATAAGAC AAAATTATTA TTATTATTAT TATTTTTACT GTTATAATTA TGTTGTCTCT

6421 TCAATGATTC ATAAATAGTT GGACTTGATT TTTAAAATGT TTATAATATG ATTAGCATAG

6481 TTAAATAAAA AAAGTTGAAA AATTAAAAAA AAACATATAA ACACAAATGA TGTTTTTTCC

6541 TTCAATTTcg gcgcctgatg cggtattttc tccttacgca tctgtgcggt atttcacacc

6601 gcatatggtg cactctcagt acaatctgct ctgatgccgc atagttaagc cagccccgac

6661 acccgccaac acccgctgac gcgccctgac gggcttgtct gctcccggca tccgcttaca

6721 gacaagctgt gaccgtctcc gggagctgca tgtgtcagag gttttcaccg tcatcaccga

6781 aacgcgcgag acgaaagggc ctcgtgatac gcctattttt ataggttaat gtcatgataa

6841 taatggtttc ttagacgtca ggtggcactt ttcggggaaa tgtgcgcgga acccctattt

6901 gtttattttt ctaaatacat tcaaatatgt atccgctcat gagacaataa ccctgataaa

6961 tgcttcaata atattgaaaa aggaagagta tgagtattca acatttccgt gtcgccctta

7021 ttcccttttt tgcggcattt tgccttcctg tttttgctca cccagaaacg ctggtgaaag

7081 taaaagatgc tgaagatcag ttgggtgcac gagtgggtta catcgaactg gatctcaaca

7141 gcggtaagat ccttgagagt tttcgccccg aagaacgttt tccaatgatg agcactttta

7201 aagttctgct atgtggcgcg gtattatccc gtattgacgc cgggcaagag caactcggtc

7261 gccgcataca ctattctcag aatgacttgg ttgagtactc accagtcaca gaaaagcatc

7321 ttacggatgg catgacagta agagaattat gcagtgctgc cataaccatg agtgataaca

7381 ctgcggccaa cttacttctg acaacgatcg gaggaccgaa ggagctaacc gcttttttgc

7441 acaacatggg ggatcatgta actcgccttg atcgttggga accggagctg aatgaagcca

7501 taccaaacga cgagcgtgac accacgatgc ctgtagcaat ggcaacaacg ttgcgcaaac

7561 tattaactgg cgaactactt actctagctt cccggcaaca attaatagac tggatggagg

7621 cggataaagt tgcaggacca cttctgcgct cggcccttcc ggctggctgg tttattgctg

7681 ataaatctgg agccggtgag cgtgggtctc gcggtatcat tgcagcactg gggccagatg

7741 gtaagccctc ccgtatcgta gttatctaca cgacggggag tcaggcaact atggatgaac

7801 gaaatagaca gatcgctgag ataggtgcct cactgattaa gcattggtaa ctgtcagacc

7861 aagtttactc atatatactt tagattgatt taaaacttca tttttaattt aaaaggatct

7921 aggtgaagat cctttttgat aatctcatga ccaaaatccc ttaacgtgag ttttcgttcc

7981 actgagcgtc agaccccgta gaaaagatca aaggatcttc ttgagatcct ttttttctgc

8041 gcgtaatctg ctgcttgcaa acaaaaaaac caccgctacc agcggtggtt tgtttgccgg

8101 atcaagagct accaactctt tttccgaagg taactggctt cagcagagcg cagataccaa

8161 atactgttct tctagtgtag ccgtagttag gccaccactt caagaactct gtagcaccgc

8221 ctacatacct cgctctgcta atcctgttac cagtggctgc tgccagtggc gataagtcgt

8281 gtcttaccgg gttggactca agacgatagt taccggataa ggcgcagcgg tcgggctgaa

8341 cggggggttc gtgcacacag cccagcttgg agcgaacgac ctacaccgaa ctgagatacc

8401 tacagcgtga gctatgagaa agcgccacgc ttcccgaagg gagaaaggcg gacaggtatc

8461 cggtaagcgg cagggtcgga acaggagagc gcacgaggga gcttccaggg ggaaacgcct

8521 ggtatcttta tagtcctgtc gggtttcgcc acctctgact tgagcgtcga tttttgtgat

8581 gctcgtcagg ggggcggagc ctatggaaaa acgccagcaa cgcggccttt ttacggttcc

8641 tggccttttg ctggcctttt gctcacatgt tctttcctgc gttatcccct gattctgtgg

8701 ataaccgtat taccgccttt gagtgagctg ataccgctcg ccgcagccga acgaccgagc

8761 gcagcgagtc agtgagcgag gaagcggaag agcgcccaat acgcaaaccg cctctccccg

8821 cgcgttggcc gattcattaa tgcagctggc acgacaggtt tcccgactgg aaagcgggca

8881 gtgagcgcaa cgcaattaat gtgagttagc tcactcatta ggcaccccag gctttacact

8941 ttatgcttcc ggctcgtatg ttgtgtggaa ttgtgagcgg ataacaattt cacacaggaa

9001 acagctatga ccatgattac gccaagcttg ataattcctg cagcccagct taattctttt

9061 cgagctcttt atgcttaagt ttacaattta atattcatac tttaagtatt ttttgtagta

9121 tcctagatat tgtgctttaa atgctcaccc ctcaaagcac cagtaatatt ttcatccact

9181 gaaataccat taaattttca aaaaaatact atgcatataa tgttatacat ataaacataa

9241 aacgccatgt aaatcaaaaa atatataaaa atatgtataa aaataaatat gcactaaata

9301 taagctaatt atgcataaaa attaaagtgc cctttattaa ctagctagtc gtaattattt

9361 atatttctat gttataaaaa aatcctcata taataatata attaatatat gtaatgtttt

9421 ttttatttta taattttaat ataaaataat atgtaaatta attcaaaaaa taaatataat

9481 tgttgtgaaa caaaaaacgt aattttttca tttgccttca aaatttaaat ttattttaat

9541 atttcctaaa atatatatac tttgtgtata aatatataaa aatatatatt tgcttataaa

9601 taaataaaaa attttataaa acataggggg atccatggtt ggttcgctaa actgcatcgt

9661 cgctgtgtcc cagaacatgg gcatcggcaa gaacggggac ctgccctggc caccgctcag

9721 gaacgaattt agatatttcc agagaatgac cacaacctct tcagtagaag gtaaacagaa

9781 tctggtgatt atgggtaaga agacctggtt ctccattcct gagaagaatc gacctttaaa

9841 gggtagaatt aatttagttc tcagcagaga actcaaggaa cctccacaag gagctcattt

9901 tctttccaga agtctagatg atgccttaaa acttactgaa caaccagaat tagcaaataa

9961 agtagacatg gtctggatag ttggtggcag ttctgtttat aaggaagcca tgaatcaccc

10021 aggccatctt aaactatttg tgacaaggat catgcaagac tttgaaagtg acacgttttt

10081 tccagaaatt gatttggaga aatataaact tctgccagaa tacccaggtg ttctctctga

10141 tgtccaggag gagaaaggca ttaagtacaa atttgaagta tatgagaaga atgattaagg

10201 atcccgtttt tcttacttat atatttatac caattgattg tatttataac tgtaaaaatg

10261 tgtatgttgt gtgcatattt ttttttgtgc atgcacatgc atgtaaatag ctaaaattat

10321 gaacatttta ttttttgttc agaaaaaaaa aactttacac acataaaatg gctagtatga

10381 atagccatat tttatataaa ttaaatccta tgaatttatg accatattaa aaatttagat

10441 atttatggaa cataatatgt ttgaaacaat aagacaaaat tattattatt attattattt

10501 ttactgttat aattatgttg tctcttcaat gattcataaa tagttggact tgatttttaa

10561 aatgtttata atatgattag catagttaaa taaaaaaagt tgaaaaatta aaaaaaaaca

10621 tataaacaca aatgatgttt tttccttcaa tttcgatg

//

**pSL1756**

LOCUS	pSL1756	10515 bp	DNA	circular	21-OCT-2024

DEFINITION .

ACCESSION

VERSION

SOURCE	.

ORGANISM .

COMMENT

COMMENT	ApEinfo:methylated:1

FEATURES	Location/Qualifiers

exon	6847..7704

/vntifkey="61"

/locus_tag="AMP"

/label="AMP"

/ApEinfo_label="AMP"

/ApEinfo_fwdcolor="pink"

/ApEinfo_revcolor="pink"

/ApEinfo_graphicformat="arrow_data {{0 1 2 0 0 -1} {} 0}

width 5 offset 0"

rep_origin	7802..8484

/locus_tag="ColE1 origin"

/label="ColE1 origin"

/ApEinfo_label="ColE1 origin"

/ApEinfo_fwdcolor="gray50"

/ApEinfo_revcolor="gray50"

/ApEinfo_graphicformat="arrow_data {{0 1 2 0 0 -1} {} 0}

width 5 offset 0"

exon	9492..10055

/vntifkey="61"

/locus_tag="hDHFR"

/label="hDHFR"

/ApEinfo_label="hDHFR"

/ApEinfo_fwdcolor="pink"

/ApEinfo_revcolor="pink"

/ApEinfo_graphicformat="arrow_data {{0 1 2 0 0 -1} {} 0}

width 5 offset 0"

misc_feature	5274..5952

/locus_tag="GFPmut2"

/label="GFPmut2"

/ApEinfo_label="GFPmut2"

/ApEinfo_fwdcolor="cyan"

/ApEinfo_revcolor="green"

/ApEinfo_graphicformat="arrow_data {{0 1 2 0 0 -1} {} 0}

width 5 offset 0"

misc_feature	721..1176

/locus_tag="DiCre 5'HR "

/label="DiCre 5'HR "

/ApEinfo_label="DiCre 5'HR "

/ApEinfo_fwdcolor="#804000"

/ApEinfo_revcolor="green"

/ApEinfo_graphicformat="arrow_data {{0 0.5 0 1 2 0 0 -1 0

-0.5} {} 0} width 5 offset 0"

misc_feature	10063..10510

/locus_tag="PbDHFR-TS 3'UTR"

/label="PbDHFR-TS 3'UTR"

/ApEinfo_label="PbDHFR-TS 3'UTR"

/ApEinfo_fwdcolor="cyan"

/ApEinfo_revcolor="green"

/ApEinfo_graphicformat="arrow_data {{0 1 2 0 0 -1} {} 0}

width 5 offset 0"

misc_feature	join(8888..9197,9202..9491)

/vntifkey="21"

/locus_tag="PbEF1a-A\5'UTR"

/label="PbEF1a-A\5'UTR"

/ApEinfo_label="PbEF1a-A\5'UTR"

/ApEinfo_fwdcolor="#eb0214"

/ApEinfo_revcolor="#eb0214"

/ApEinfo_graphicformat="arrow_data {{0 1 2 0 0 -1} {} 0}

width 5 offset 0"

misc_feature	7..696

/locus_tag="DiCre 3' HR"

/label="DiCre 3' HR"

/ApEinfo_label="DiCre 3' HR"

/ApEinfo_fwdcolor="#c0c0c0"

/ApEinfo_revcolor="green"

/ApEinfo_graphicformat="arrow_data {{0 0.5 0 1 2 0 0 -1 0

-0.5} {} 0} width 5 offset 0"

misc_feature	4656..5229

/locus_tag="PbEF1a promoter (from pSL0281)"

/label="PbEF1a promoter (from pSL0281)"

/ApEinfo_label="PbEF1a promoter (from pSL0281)"

/ApEinfo_fwdcolor="cyan"

/ApEinfo_revcolor="green"

/ApEinfo_graphicformat="arrow_data {{0 1 2 0 0 -1} {} 0}

width 5 offset 0"

misc_feature	10062..10512

/locus_tag="PbDHFR-TS 3'UTR(1)"

/label="PbDHFR-TS 3'UTR(1)"

/ApEinfo_label="PbDHFR-TS 3'UTR"

/ApEinfo_fwdcolor="cyan"

/ApEinfo_revcolor="green"

/ApEinfo_graphicformat="arrow_data {{0 1 2 0 0 -1} {} 0}

width 5 offset 0"

misc_feature	10056..10512

/vntifkey="21"

/locus_tag="PbDHFR/TS\3'UTR"

/label="PbDHFR/TS\3'UTR"

/ApEinfo_label="PbDHFR/TS\3'UTR"

/ApEinfo_fwdcolor="#eb0214"

/ApEinfo_revcolor="#eb0214"

/ApEinfo_graphicformat="arrow_data {{0 1 2 0 0 -1} {} 0}

width 5 offset 0"

misc_feature	5959..6405

/locus_tag="PbDHFR/TS\3'UTR(1)"

/label="PbDHFR/TS\3'UTR(1)"

/ApEinfo_label="PbDHFR/TS\3'UTR"

/ApEinfo_fwdcolor="#ff0000"

/ApEinfo_revcolor="green"

/ApEinfo_graphicformat="arrow_data {{0 1 2 0 0 -1} {} 0}

width 5 offset 0"

CDS	7045..7704

/locus_tag="AmpR"

/label="AmpR"

/ApEinfo_label="AmpR"

/ApEinfo_fwdcolor="yellow"

/ApEinfo_revcolor="yellow"

/ApEinfo_graphicformat="arrow_data {{0 1 2 0 0 -1} {} 0}

width 5 offset 0"

misc_feature	join(8888..9197,9202..9486)

/locus_tag="PbEF1a 5'UTR"

/label="PbEF1a 5'UTR"

/ApEinfo_label="PbEF1a 5'UTR"

/ApEinfo_fwdcolor="#804040"

/ApEinfo_revcolor="#804040"

/ApEinfo_graphicformat="arrow_data {{0 1 2 0 0 -1} {} 0}

width 5 offset 0"

misc_feature	1183..4640

/locus_tag="S1 Promoter-ETS-18S-ITS1-5.8S"

/label="S1 Promoter-ETS-18S-ITS1-5.8S"

/ApEinfo_label="S1 Promoter-ETS-18S-ITS1-5.8S"

/ApEinfo_fwdcolor="#ff8040"

/ApEinfo_revcolor="green"

/ApEinfo_graphicformat="arrow_data {{0 0.5 0 1 2 0 0 -1 0

-0.5} {} 0} width 5 offset 0"

ORIGIN

1 ggtaccCTTA ACTAACGACA AATTGTGATG ATCTTACTGA AATTAAAATA TTACAAAATA

61 AGTTGCAAAT ATTTGTCTTT GCATCTATAA AAATATAATT TTATTTTGCT TTTCTATGAT

121 TCACTAATAT ATCACAATAG ATGTATTCAA AGCAAAATAA ATAAGCATTA CATGTGTAAT

181 ATATTTGCAA CACTTTCTCA TAAAATAAAA ATTAATAACT ACATGAATAT AACGAATATG

241 ATTTTATTTT TCTTTTTTTT ATCCAAAGAT ATATAGCGAT ATCATCTATA TTATTTTATT

301 AGTTTTTTTC GAAAATTATA AAACAAACAT AACTGTCTAT TATATTATTT ATCTCATAAT

361 ATATATATAT ACAAAATAAA ATACATTTTA AAAAAATATA AATTTAATTT TTTTATATTT

421 TCAAAAGTAT AAAAATAACC TAAAAACTTA TTAATTTTTT ACATAAAAAA TATAGACAAG

481 TATTAAAAAA AAAAAAAAAA AAAAACGACA AATCGCACAT TTAAAAAAAG ATACACATGT

541 GCAATGTAGT TATATATAAT TTTAAAGATA ATTTATTTAA TAAAACTGGT TTTAAATTAT

601 TAATTATAAA ATGGAAAAAT AAATAAAGAG TGTAAACAAT TATATATATT TTTTTTCTCC

661 CATTTCTCAT TATCTTTATA TAGTGCATCT TTCTACgatc CCCGGGgatc AGGCCTgatc

721 CGTGTTTTGC GATTTCTTTG GGTTCGTGTT TTGCGATTTG TTTGGGTTCG TGTTTTGTGA

781 TTTCTTTGCG ATTTTTGTGT GTTTAGTGAT TTTTGTGTGT GTCGATTGTG CGTTTTTTAT

841 TATATTATTG TTATCATATT TGATTTACCT TTTGATATTA GGACATATAT ATATGCATAT

901 ATTTTTTTGT ATTTCCTGAT TTGTTTGTGA TCATAATTTT CGATAAAATG AGTTTTTATT

961 ATTGATGCTT TAAACATTCC AGTTGGTTCC AGATCTCTTT TTTCTCGTCT CGTTTAGAAA

1021 TTATTTCATT TTTGCATATT CTTGCTCATA TGCCAATGTC TTACATGTTT TTCGTAACAA

1081 CTTTCACGCT AGCTACTTAT TTCACAGTGA TACTTCACTA GTATAAGTTG CATCTAATTT

1141 ATATTGGCAA CTCTAATAGA ACCCCTAAAT TTATGGgggc ccCCGAACTC GTATGGGAAA

1201 GAAGTATCAA GTCAATTGCA TATAAAAATA AAATTGGACC AATGATTGAA AAATCGAAAT

1261 TGTTAATCAT TATATGATAT TATATAATAG TTGATAATAA GTAGTTTAGT TAGTTTTAGT

1321 TATTATATGT AAATTAATAA TTTATGTATG TTTCTGAAAG GTACGAGGTT TTAAATAATA

1381 CTTTCAATTT TAATTTTTTA ATATTCGAAT GTATAATGAA ACGACGACCA ACCAATGTAT

1441 ATAATTATTT TAAATAATTA TATATGCAAT ATGTAATAAA TATGTTTTAG GATATATTAT

1501 AGCATTTGAA AAAATGTGTG TAGTGTGTTT GTAAAGTTAT AGAAAACATC GTGGTGTTAT

1561 GTTATAACAA TATAAAACAA AATACACGAT AATATAGAAT AAAAAATATA ATACATAATA

1621 ATGTATATTA TTTTAAGCAA AAAAATAATT TTGTTAACCC CTTGTTTTTA ATTTGGACCA

1681 CATTTGTTAA AAAATTAGTT TAATATAAAA TAAATTTTTA ATAAATAAAA TCGTAAAGAA

1741 TTGTAAAAGA GTTTCAATAT TATCGGGACC TTTTATGATA TTAATGTATA AATCCGTAAT

1801 TTGTATACTA TATAAATTAG TTTTTGAAAT TAGCAGCATA GCAAAAAATA TATTGTCTTT

1861 TAAGTGTGTG TATCTAATAC CACCCAAAAT AAAGGATGAT ATATTTTTAT GCTAGGTGCA

1921 TTTATTTTTT TTATTATACT ATAAATAAAA TATAGTATGA TAAAGAAAAA ATAGCAAATA

1981 AAATATCACA TGTAAATGAA ATAAACACTA TAATAAGTGT ATTTTTCTAA AATATGTAAA

2041 ACCTATATAA TATTATTAAA ATGATGATGT ACCCATCTAT TATATATAAT ATAATTGATA

2101 TACTTAATTT TTTTTATGTA TATTAAAATT ATTAAATGTA ATATTTGGAT ATTTCATATA

2161 TTATTAATAT TTATATTAAC CTGGTTGATC TTGCCAGTAG TCATATGCTT GTCTCAAAGA

2221 TTAAGCCATG CAAGTGAAAG TATATACACA ATTATTGTAG AAACTGCGAA CGGCTCATTA

2281 AAACAGTTAT AATCTACTTG ACATTTTATT ATAAGGATAA CTACGGAAAA TCTGTAGCTA

2341 ATACTTGTTT TAAGTACTTT TACTCCTCGG AGTAATTGTA TGTATTTGTT AAGATCCCTA

2401 AGAAAAAATG ATATTAAAGG AATTATAACA AAGAAGCGAC ACATAATGTA ATATTCAGTG

2461 TGTATCAATC GAGTTTCTGA CCTATCAGCT TTTGATGTTA GGGTATTGAC CTAACATGGC

2521 TTTGACGGGT AACGGGGAAT TAGAGTTCGA TTCCGGAGAG GGAGCCTGAG AAATAGCTAC

2581 CACATCTAAG GAAGGCAGCA GGCGCGTAAA TTACCCAATT CTAAATAAGA GAGGTAGTGA

2641 CAAGAAATAA CAATATAAGG CCAAATTTTG GTTTTATAAT TGGAATGATG GGAATTTAAA

2701 ACCTTCCCAA AAATCAATTG GAGGGCAAGT CTGGTGCCAG CAGCCGCGGT AATTCCAGCT

2761 CCAATAGCGT ATATTAAAAT TGTTGCAGTT AAAACGCTCG TAGTTTAATT TCAAGGGTAA

2821 TATTATTTTA AGTACTAACT TGGAGAAAAT TGCGACTTCT GTCACTGTTT TCATCCTTGC

2881 TGATGTTCTT TTAATTACAG ACCCTTTGAG AGCCCATTAA TTTATGACTG GGTTTCTCGT

2941 TACTTTGAGT AAATTAGAGT GTTCAAAGCA AACAGTTAAA GCGTTTTCGC GTTTGAATAT

3001 TATAGCATGG AATAACAACA TTGAATAAGT CAAAAGTTTT TGAAAAACTT TTCTTATTTT

3061 GGCTTAGTTA CGATTAATAG GAGTAGCTTG GGGGCATTTG TATTCAGATG TCAGAGGTGA

3121 AATTCTTAGA TTTTCTGGAG ACAAACAACT GCGAAAGCAT TTGCCTAAAA TACTTCCATT

3181 AATCAAGAAC GAAAGTTAAG GGAGTGAAGA CGATCAGATA CCGTCGTAAT CTTAACCATA

3241 AACTATGCCG ACTAGGTTTT GGATGAAAAT TTATAAATAA GATTTCCCTC CGGGGAATTC

3301 TTAGATTGCT TCCTTCAGTA CCTTATGAGA AATCAAAGTC TTTGGGTTCT GGGGCGAGTA

3361 TTCGCGCAAG CGAGAAAGTT AAAAGAATTG ACGGAAGGGC ACCACCAGGC GTGGAGCTTG

3421 CGGCTTAATT TGACTCAACA CGGGGAAACT CACTAGTTTA AGACAAGAGT AGGATTGACA

3481 GATTAATAGC TCTTTCTTGA TTTCTTGGAT GGTGATGCAT GGCCGTTTTT AGTTCGTGAA

3541 TATGATTTGT CTGGTTAATT CCGATAACGA ACGAGATCTT AACCTGCTAA TTAGCGGCGA

3601 GTACGCTATA TCCTTTATCG GGGGATTGGT TTTGACGTTT ATGCGGTCAT ACTGCTTAAT

3661 CAATTGGTTT ACCTTTTGCT CTTTTGCGGT ATGTTTCTCG CATCTTCGAC ATGCCTCTCT

3721 TCTGATAAGG ATGTATTCGC TTTATTTAAT GCTTCTTAGA GGAACGATGT GTGTCTAACA

3781 CAAGGAAGTT TAAGGCAACA ACAGGTCTGT GATGTCCTTA GATATACTAG GCTGCACGCG

3841 TGCTACACTG ATATGTAAAA CGAGTATTTA AAATTATATC TGCGCGTGGG TGTCAAAGCC

3901 TACACGTCAG CATATATTTT TCCTCCACTG AAAAGTGTAG GTAATCTTTA TCAATACATA

3961 TCGTGATGGG GATAGATTAT TGCAATTATT AATCTTGAAC GAGGAATGCC TAGTAAGCAT

4021 GATTCATCAG ATTGTGCTGA CTACGTCCCT GCCCTTTGTA CACACCGCCC GTCGCTCCTA

4081 CCGATTGAAA GATATGATGA ATTGTTTGGA CAAGAAAATA GAAATTTTAT TTTTATTTTT

4141 TTTGGAAGGA CCGTAAATCC TATCTTTTAA AGGAAGGAGA AGTCGTAACA AGGTTTCCGT

4201 AGGTGAACCT GCGGAAGGAT CATTATTAAT ACGATTTAAA GATTTGTATA CTAATATTTA

4261 ATGGATAATA ATTTTGTATT TATATGAAAT TGTTATTAAT TTCATTAAAT AATAGTATAT

4321 TAATCTTTGT ATATATTTTT ATTATTGTAT TAAATATATA TATATTGATT TGCTATATAT

4381 ACTTGTATAT GTAATGAATT GATTTATATA TGTTTAGTAT ATATAATAAA AATATATACA

4441 TTACAAGTCT TAACTATCAA TTTTTGCATA ACTTTTATTT TAATTATAAA TTTTGTGTTT

4501 ATTTTTTTAT TATAAATCGA TTTAATATAT TATCTGTATT TAACATATAA AATTGTATTA

4561 TATAAAAATC TAAATTTATC ACAATCTTAT TAGCACAATC TTAACGATGG ATGTCTTGGT

4621 TCCTACAGCG ATGAAGGCCG gcggccgcTc tcgagAGCTT AATTCTTTTC GAGCTCTTTA

4681 TGCTTAAGTT TACAATTTAA TATTCATACT TTAAGTATTT TTTGTAGTAT CCTAGATATT

4741 GTGCTTTAAA TGCTCACCCC TCAAAGCACC AGTAATATTT TCATCCACTG AAATACCATT

4801 AAATTTTCAA AAAAATACTA TGCATATAAT GTTATACATA TAAACATAAA ACGCCATGTA

4861 AATCAAAAAA TATATAAAAA TATGTATAAA AATAAATATG CACTAAATAT AAGCTAATTA

4921 TGCATAAAAA TTAAAGTGCC CTTTATTAAC TAGctagTCG TAATTATTTA TATTTCTATG

4981 TTATAAAAAA ATCCTCATAT AATAATATAA TTAATATATG TAATGTTTTT TTTATTTTAT

5041 AATTTTAATA TAAAATAATA TGTAAATTAA TTCAAAAAAT AAATATAATT GTTGTGAAAC

5101 AAAAAACGTA ATTTTTTCAT TTGCCTTCAA AATTTAAATT TATTTTAATA TTTCCTAAAA

5161 TATATATACT TTGTGTATAA ATATATAAAA ATATATATTT GCTTATAAAT AAATAAAAAT

5221 TTTATAAAAa tgactagtag taaaggagaa gaacttttca ctggagttgt cccAATTCTT

5281 GTTGAATTAG ATGGTGATGT TAATGGGCAC AAATTTTCTG TCAGTGGAGA GGGTGAAGGT

5341 GATGCAACAT ACGGAAAACT TACCCTTAAA TTTATTTGCA CTACTGGAAA ACTACCTGTT

5401 CCATGGCCAA CACTTGTCAC TACTTTCGCG TATGGTCTTC AATGCTTTGC GAGATACCCA

5461 GATCATATGA AACAGCATGA CTTTTTCAAG AGTGCCATGC CCGAAGGTTA TGTACAGGAA

5521 AGAACTATAT TTTTCAAAGA TGACGGGAAC TACAAGACAC GTGCTGAAGT CAAGTTTGAA

5581 GGTGATACCC TTGTTAATAG AATCGAGTTA AAAGGTATTG ATTTTAAAGA AGATGGAAAC

5641 ATTCTTGGAC ACAAATTGGA ATACAACTAT AACTCACACA ATGTATACAT CATGGCAGAC

5701 AAACAAAAGA ATGGAATCAA AGTTAACTTC AAAATTAGAC ACAACATTGA AGATGGAAGC

5761 GTTCAACTAG CAGACCATTA TCAACAAAAT ACTCCAATTG GCGATGGCCC TGTCCTTTTA

5821 CCAGACAACC ATTACCTGTC CACACAATCT GCCCTTTCGA AAGATCCCAA CGAAAAGAGA

5881 GACCACATGG TCCTTCTTGA GTTTGTAACA GCTGCTGGGA TTACACATGG CATGGATGAA

5941 CTATACAAAT AAggatccGT TTTTCTTACT TATATATTTA TACCAATTGA TTGTATTTAT

6001 AACTGTAAAA ATGTGTATGT TGTGTGCATA TTTTTTTTTG TGCATGCACA TGCATGTAAA

6061 TAGCTAAAAT TATGAACATT TTATTTTTTG TTCAGAAAAA AAAACTTTAC ACACATAAAA

6121 TGGCTAGTAT GAATAGCCAT ATTTTATATA AATTAAATCC TATGAATTTA TGACCATATT

6181 AAAAATTTAG ATATTTATGG AACATAATAT GTTTGAAACA ATAAGACAAA ATTATTATTA

6241 TTATTATTAT TTTTACTGTT ATAATTATGT TGTCTCTTCA ATGATTCATA AATAGTTGGA

6301 CTTGATTTTT AAAATGTTTA TAATATGATT AGCATAGTTA AATAAAAAAA GTTGAAAAAT

6361 TAAAAAAAAA CATATAAACA CAAATGATGT TTTTTCCTTC AATTTcggcg cctgatgcgg

6421 tattttctcc ttacgcatct gtgcggtatt tcacaccgca tatggtgcac tctcagtaca

6481 atctgctctg atgccgcata gttaagccag ccccgacacc cgccaacacc cgctgacgcg

6541 ccctgacggg cttgtctgct cccggcatcc gcttacagac aagctgtgac cgtctccggg

6601 agctgcatgt gtcagaggtt ttcaccgtca tcaccgaaac gcgcgagacg aaagggcctc

6661 gtgatacgcc tatttttata ggttaatgtc atgataataa tggtttctta gacgtcaggt

6721 ggcacttttc ggggaaatgt gcgcggaacc cctatttgtt tatttttcta aatacattca

6781 aatatgtatc cgctcatgag acaataaccc tgataaatgc ttcaataata ttgaaaaagg

6841 aagagtatga gtattcaaca tttccgtgtc gcccttattc ccttttttgc ggcattttgc

6901 cttcctgttt ttgctcaccc agaaacgctg gtgaaagtaa aagatgctga agatcagttg

6961 ggtgcacgag tgggttacat cgaactggat ctcaacagcg gtaagatcct tgagagtttt

7021 cgccccgaag aacgttttcc aatgatgagc acttttaaag ttctgctatg tggcgcggta

7081 ttatcccgta ttgacgccgg gcaagagcaa ctcggtcgcc gcatacacta ttctcagaat

7141 gacttggttg agtactcacc agtcacagaa aagcatctta cggatggcat gacagtaaga

7201 gaattatgca gtgctgccat aaccatgagt gataacactg cggccaactt acttctgaca

7261 acgatcggag gaccgaagga gctaaccgct tttttgcaca acatggggga tcatgtaact

7321 cgccttgatc gttgggaacc ggagctgaat gaagccatac caaacgacga gcgtgacacc

7381 acgatgcctg tagcaatggc aacaacgttg cgcaaactat taactggcga actacttact

7441 ctagcttccc ggcaacaatt aatagactgg atggaggcgg ataaagttgc aggaccactt

7501 ctgcgctcgg cccttccggc tggctggttt attgctgata aatctggagc cggtgagcgt

7561 gggtctcgcg gtatcattgc agcactgggg ccagatggta agccctcccg tatcgtagtt

7621 atctacacga cggggagtca ggcaactatg gatgaacgaa atagacagat cgctgagata

7681 ggtgcctcac tgattaagca ttggtaactg tcagaccaag tttactcata tatactttag

7741 attgatttaa aacttcattt ttaatttaaa aggatctagg tgaagatcct ttttgataat

7801 ctcatgacca aaatccctta acgtgagttt tcgttccact gagcgtcaga ccccgtagaa

7861 aagatcaaag gatcttcttg agatcctttt tttctgcgcg taatctgctg cttgcaaaca

7921 aaaaaaccac cgctaccagc ggtggtttgt ttgccggatc aagagctacc aactcttttt

7981 ccgaaggtaa ctggcttcag cagagcgcag ataccaaata ctgttcttct agtgtagccg

8041 tagttaggcc accacttcaa gaactctgta gcaccgccta catacctcgc tctgctaatc

8101 ctgttaccag tggctgctgc cagtggcgat aagtcgtgtc ttaccgggtt ggactcaaga

8161 cgatagttac cggataaggc gcagcggtcg ggctgaacgg ggggttcgtg cacacagccc

8221 agcttggagc gaacgaccta caccgaactg agatacctac agcgtgagct atgagaaagc

8281 gccacgcttc ccgaagggag aaaggcggac aggtatccgg taagcggcag ggtcggaaca

8341 ggagagcgca cgagggagct tccaggggga aacgcctggt atctttatag tcctgtcggg

8401 tttcgccacc tctgacttga gcgtcgattt ttgtgatgct cgtcaggggg gcggagccta

8461 tggaaaaacg ccagcaacgc ggccttttta cggttcctgg ccttttgctg gccttttgct

8521 cacatgttct ttcctgcgtt atcccctgat tctgtggata accgtattac cgcctttgag

8581 tgagctgata ccgctcgccg cagccgaacg accgagcgca gcgagtcagt gagcgaggaa

8641 gcggaagagc gcccaatacg caaaccgcct ctccccgcgc gttggccgat tcattaatgc

8701 agctggcacg acaggtttcc cgactggaaa gcgggcagtg agcgcaacgc aattaatgtg

8761 agttagctca ctcattaggc accccaggct ttacacttta tgcttccggc tcgtatgttg

8821 tgtggaattg tgagcggata acaatttcac acaggaaaca gctatgacca tgattacgcc

8881 aagcttgata attcctgcag cccagcttaa ttcttttcga gctctttatg cttaagttta

8941 caatttaata ttcatacttt aagtattttt tgtagtatcc tagatattgt gctttaaatg

9001 ctcacccctc aaagcaccag taatattttc atccactgaa ataccattaa attttcaaaa

9061 aaatactatg catataatgt tatacatata aacataaaac gccatgtaaa tcaaaaaata

9121 tataaaaata tgtataaaaa taaatatgca ctaaatataa gctaattatg cataaaaatt

9181 aaagtgccct ttattaacta gctagtcgta attatttata tttctatgtt ataaaaaaat

9241 cctcatataa taatataatt aatatatgta atgttttttt tattttataa ttttaatata

9301 aaataatatg taaattaatt caaaaaataa atataattgt tgtgaaacaa aaaacgtaat

9361 tttttcattt gccttcaaaa tttaaattta ttttaatatt tcctaaaata tatatacttt

9421 gtgtataaat atataaaaat atatatttgc ttataaataa ataaaaaatt ttataaaaca

9481 tagggggatc catggttggt tcgctaaact gcatcgtcgc tgtgtcccag aacatgggca

9541 tcggcaagaa cggggacctg ccctggccac cgctcaggaa cgaatttaga tatttccaga

9601 gaatgaccac aacctcttca gtagaaggta aacagaatct ggtgattatg ggtaagaaga

9661 cctggttctc cattcctgag aagaatcgac ctttaaaggg tagaattaat ttagttctca

9721 gcagagaact caaggaacct ccacaaggag ctcattttct ttccagaagt ctagatgatg

9781 ccttaaaact tactgaacaa ccagaattag caaataaagt agacatggtc tggatagttg

9841 gtggcagttc tgtttataag gaagccatga atcacccagg ccatcttaaa ctatttgtga

9901 caaggatcat gcaagacttt gaaagtgaca cgttttttcc agaaattgat ttggagaaat

9961 ataaacttct gccagaatac ccaggtgttc tctctgatgt ccaggaggag aaaggcatta

10021 agtacaaatt tgaagtatat gagaagaatg attaaggatc ccgtttttct tacttatata

10081 tttataccaa ttgattgtat ttataactgt aaaaatgtgt atgttgtgtg catatttttt

10141 tttgtgcatg cacatgcatg taaatagcta aaattatgaa cattttattt tttgttcaga

10201 aaaaaaaaac tttacacaca taaaatggct agtatgaata gccatatttt atataaatta

10261 aatcctatga atttatgacc atattaaaaa tttagatatt tatggaacat aatatgtttg

10321 aaacaataag acaaaattat tattattatt attattttta ctgttataat tatgttgtct

10381 cttcaatgat tcataaatag ttggacttga tttttaaaat gtttataata tgattagcat

10441 agttaaataa aaaaagttga aaaattaaaa aaaaacatat aaacacaaat gatgtttttt

10501 ccttcaattt cgatg

//
