## Supplementary material for "Specialized S-type ribosomes of *Plasmodium yoelii* enhance host-to-vector malaria transmission": Figure S1

McGee *et al.* Figure S1

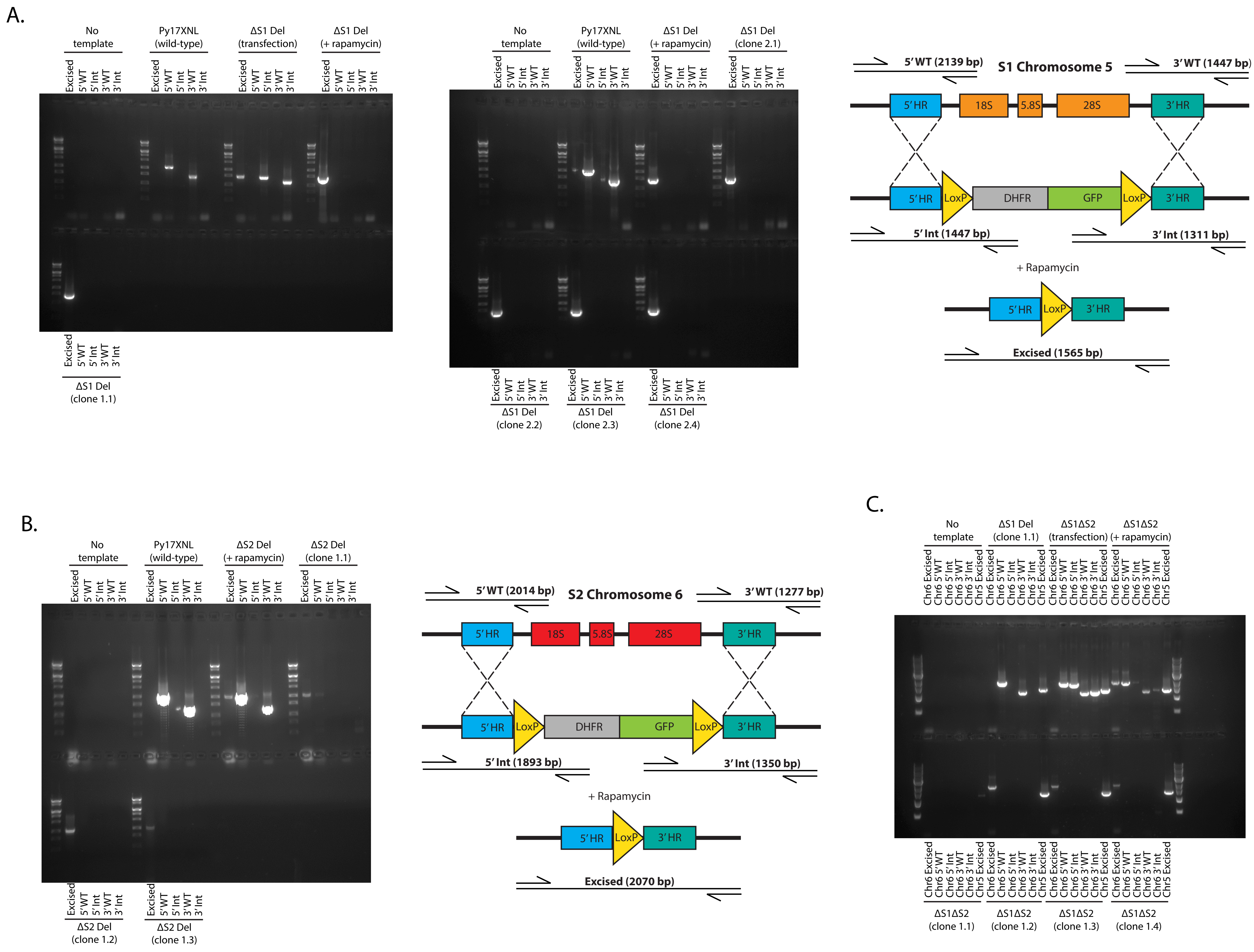


**Figure S1: Genotyping PCRs of PyDiCre S-type rRNA Deletion Lines**

Genomic DNA from transgenic parasites was compared to Py17XNL wild-type parasites or no template controls by genotyping PCR. **A)** Genotyping evidence the PyΔS1 parasite line was generated by targeting the Chromosome 5 S1 rDNA locus with a schematic of the wild type, transgenic, and excised rDNA locus-of-interest provided to the right of the gel image. **B)** Genotyping evidence the PyΔS2 parasite line was generated by targeting the Chromosome 6 S2 rDNA locus with a schematic of the wild type, transgenic, and excised rDNA locus-of-interest provided to the right of the gel image. **C)** Genotyping evidence the PyΔS1ΔS2 parasite line was generated by targeting the Chromosome 6 S2 rDNA locus in PyΔS1 clone 1.1. Multiple clones of each parasite type were isolated, and PyΔS1 clones 1.1 and 2.1, PyΔS2 clones 1.2 and 1.3, and PyΔS1ΔS2 clones 1.2 and 1.3 were used for further study. The PSU 1kb DNA ladder flanks all experimental lanes.
