## Supplementary material for "Specialized S-type ribosomes of *Plasmodium yoelii* enhance host-to-vector malaria transmission": Figure S2

McGee *et al.* Figure S2

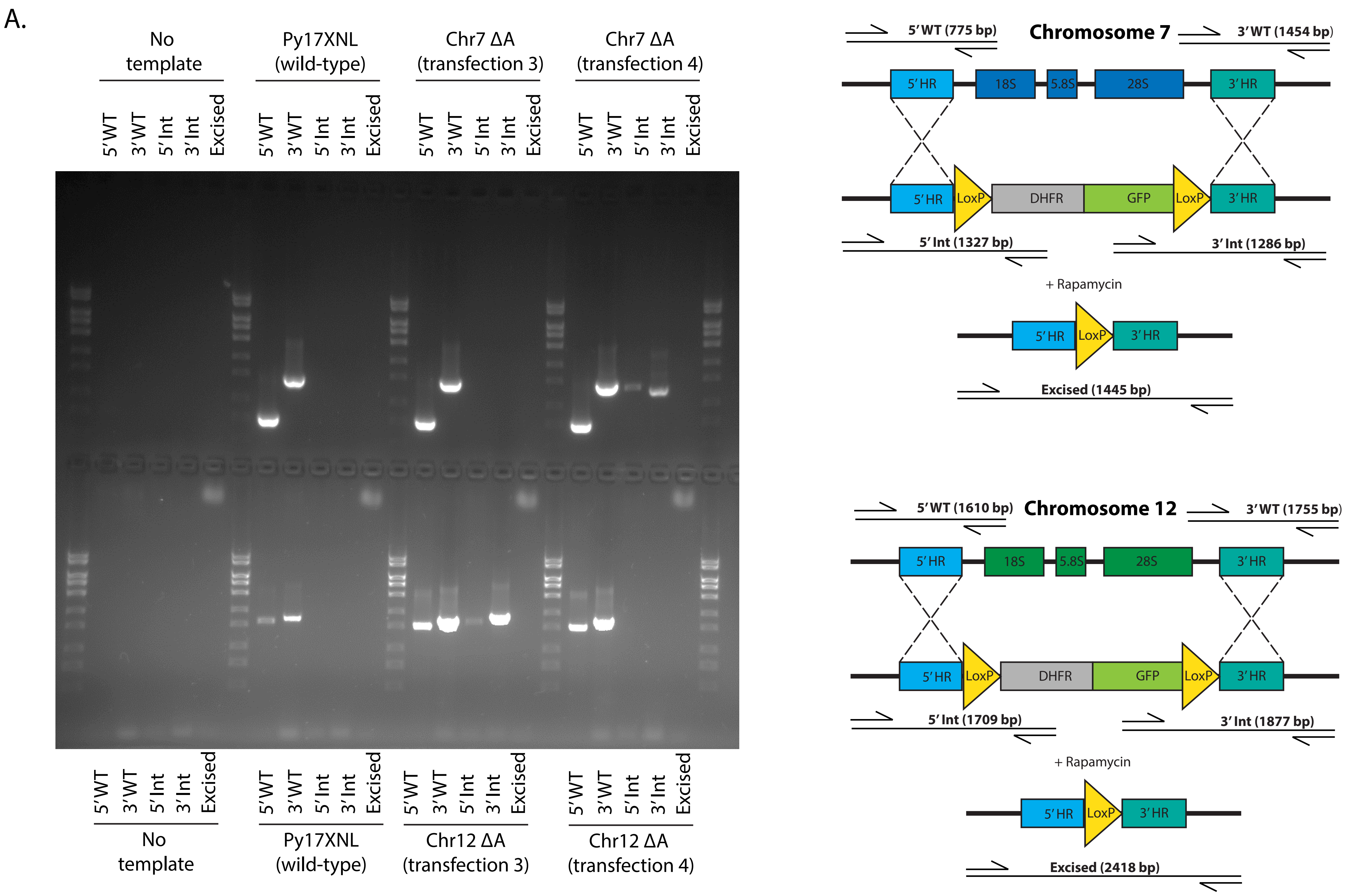


**Figure S2: Genotyping PCRs of PyDiCre A-type rRNA Deletion Lines**
Genomic DNA from transgenic parasites was compared to Py17XNL wild-type parasites or no template controls by genotyping PCR. **A)** Genotyping evidence that either A-type rDNA locus on Chromosome 7 (top row) or Chromosome 12 (bottom row) can be individually targeted and deleted. A schematic of the wild type, transgenic, and excised rDNA locus-of-interest is provided next to the corresponding row of the gel image. The PSU 1kb DNA ladder flanks all experimental lanes.
