## Supplementary material for "Specialized S-type ribosomes of *Plasmodium yoelii* enhance host-to-vector malaria transmission": Figure S3

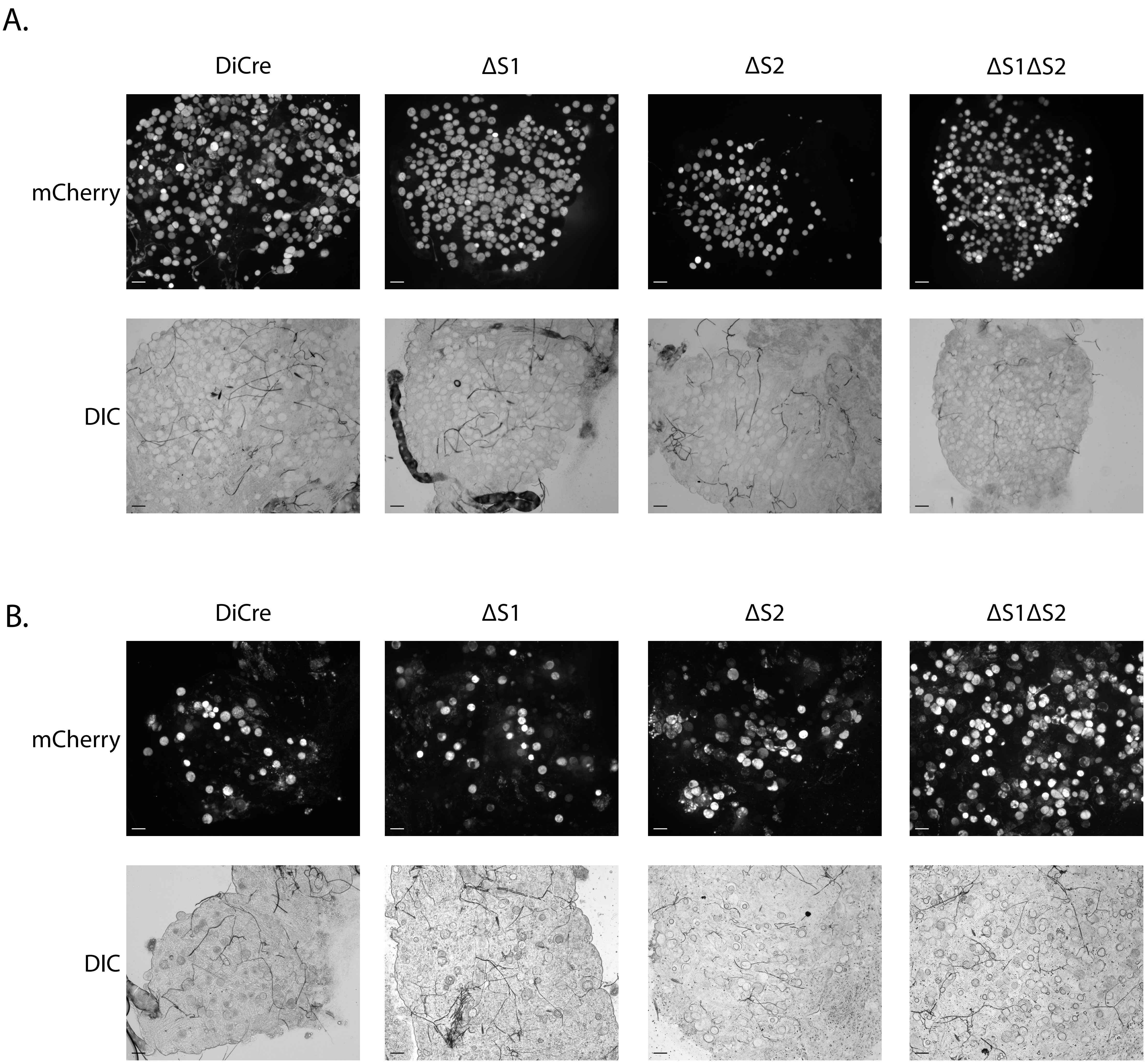


**Figure S3: Representative images of midguts infected with S-type deletion lines**
DIC and live fluorescence microscopy images using a 10x objective of dissected mosquito midguts infected with S-type deletion lines or control on **A)** day 7 (related to Fig 1A and B) and **B)** day 14 after blood meal. Scale bars = 50µm.
