## Supplementary material for "Specialized S-type ribosomes of *Plasmodium yoelii* enhance host-to-vector malaria transmission": Figure S4

A.

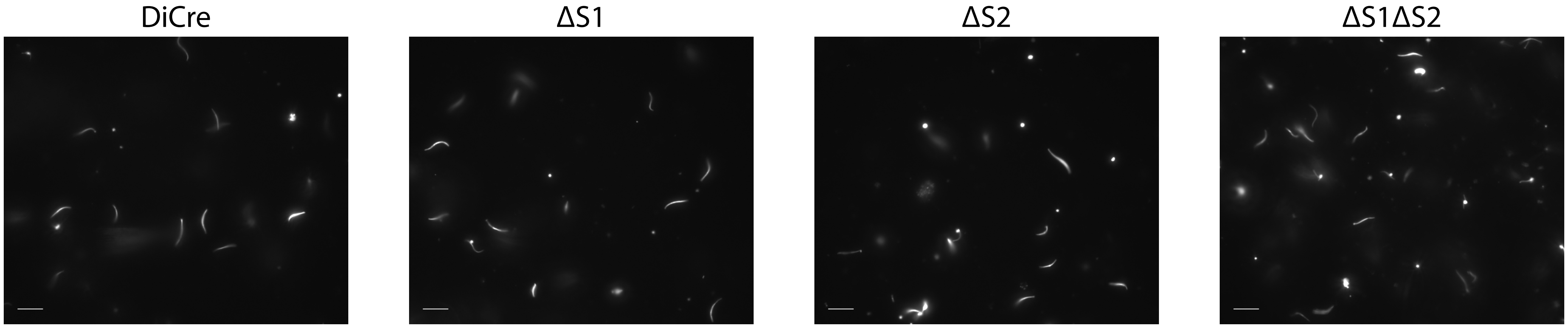


**Figure S4: Representative images of sporozoites from control and S-type deletion lines**
**A)** Representative live fluorescent images of sporozoites from dissected salivary glands of infected mosquitoes 14 days post blood meal using a 63x objective are shown (scale bar = 10μm).
