## Supplementary material for "Specialized S-type ribosomes of *Plasmodium yoelii* enhance host-to-vector malaria transmission": Figure S5

McGee *et al.* Figure S5

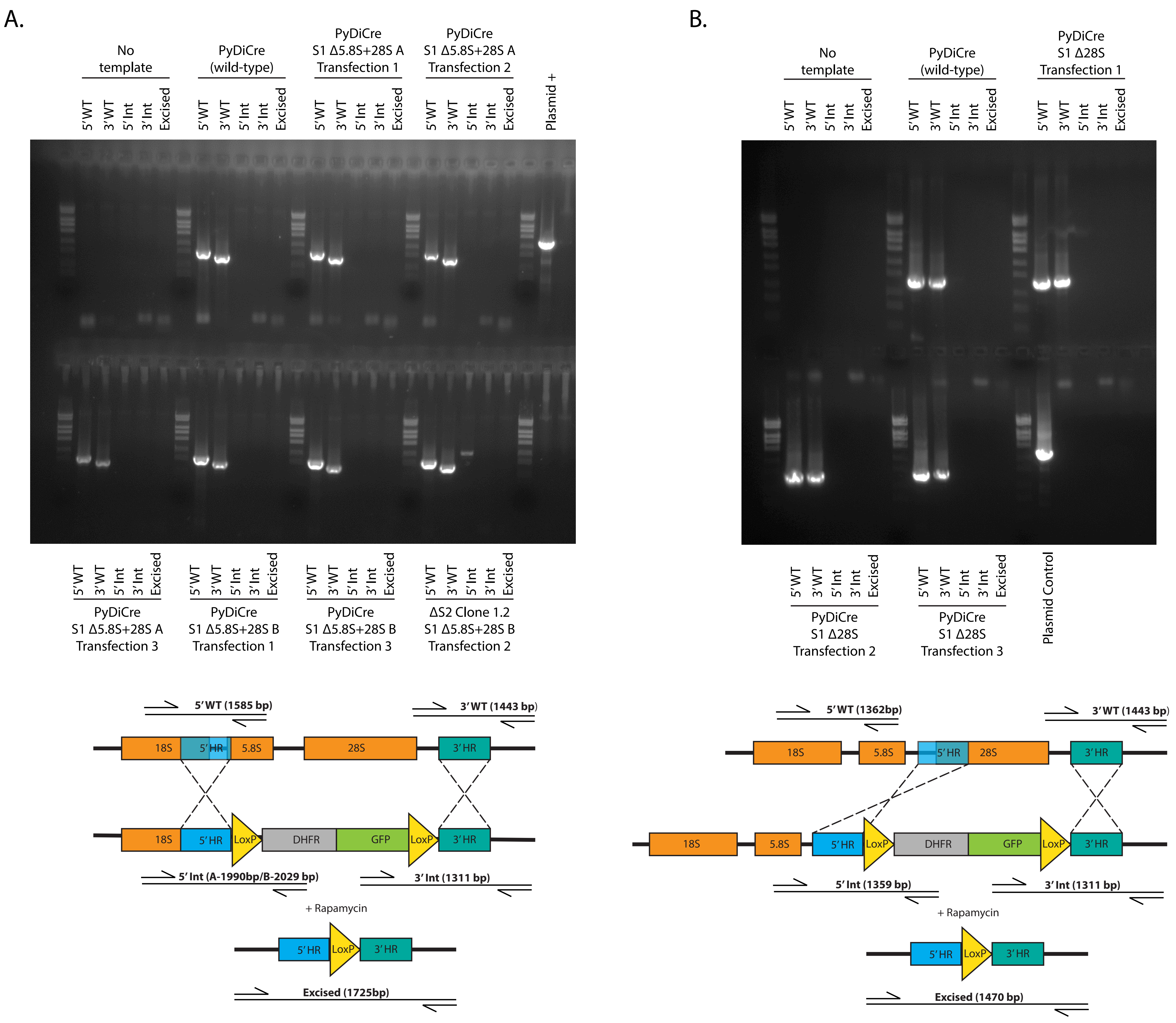


**Figure S5: Genotyping PCRs of PyDiCre S1 LSU sequence deletion attempts**
Genomic DNA from transgenic parasites was compared to that of PyDiCre parental control (wild-type) parasites or no template controls by genotyping PCR. **A)** Two constructs (designated A and B that differed in their 5'HR length) were designed for S1 Δ5.8S+28S and transfected into either PyDiCre or ΔS2 clone 1.2. No evidence of transgenic parasites was observed when the 5.8S and 28S rDNA of S1 on chromosome 5 was targeted for deletion. **B)** No evidence of transgenic parasites was observed when the 28S rDNA of S1 on chromosome 5 was targeted for deletion. A schematic of the wild type, transgenic, and excised rDNA locus-of-interest is provided below each gel image. The PSU 1kb DNA ladder flanks all experimental lanes.
