## Supplementary material for "Specialized S-type ribosomes of *Plasmodium yoelii* enhance host-to-vector malaria transmission": Figure S6

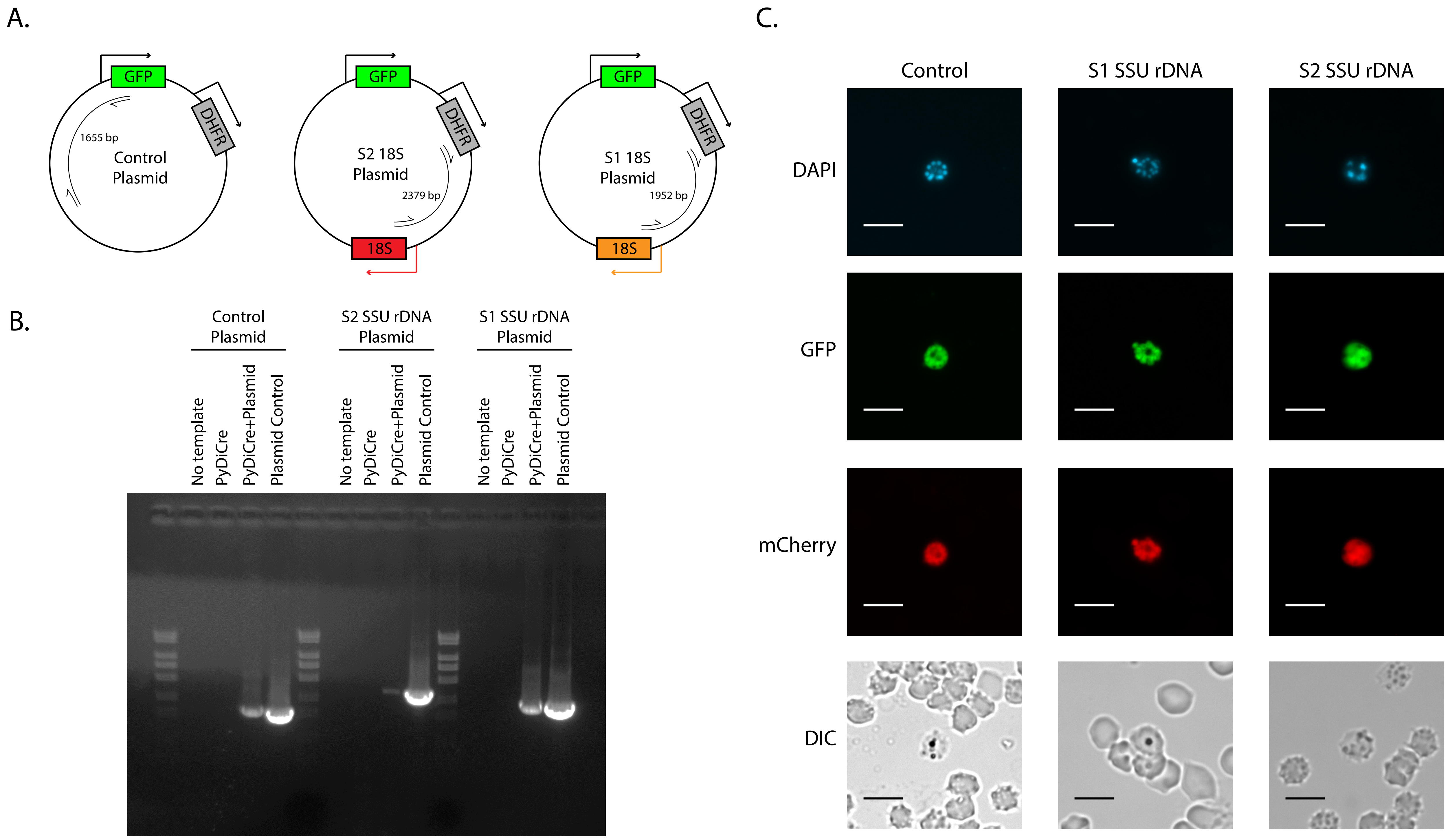


**Figure S6: Genotyping PCRs and live fluorescence GFP expression confirm the presence and retention of control or S-type SSU rDNA plasmids**
**A)** Schematic of the control, S2 SSU rDNA, and S1 SSU rDNA plasmids. **B)** Genomic DNA from parasites transfected with control or S-type SSU rDNA plasmids was compared to that of PyDiCre parental, no template control, or plasmid positive control. The PSU 1kb DNA ladder flanks all experimental lanes. **C)** DIC and live fluorescence microscopy images using a 63x objective of parasites containing either the control, S1 SSU rDNA, or S2 SSU rDNA plasmid. Nuclear DNA was stained with DAPI (scale bars = 5μm).
