## Supplementary material for "Specialized S-type ribosomes of *Plasmodium yoelii* enhance host-to-vector malaria transmission": Figure S7

A.

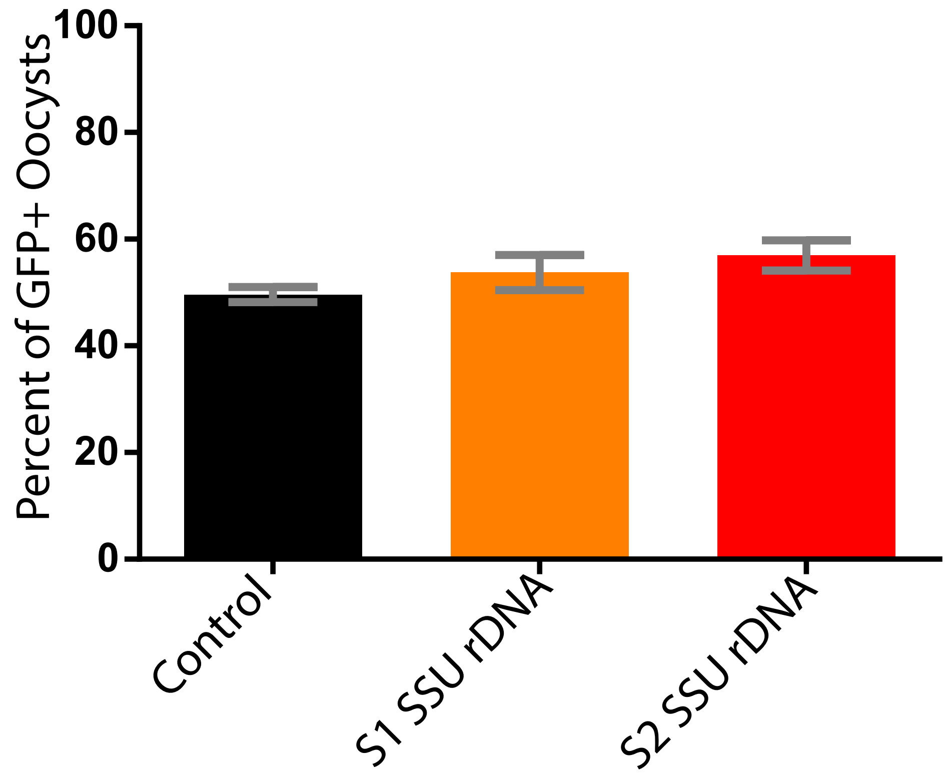


**Figure S7: Percent of oocysts with plasmid derived GFP expression 7 days post-blood meal** Mosquitoes took an infectious blood meal from mice infected with parasites transfected with either the S1 SSU rDNA, S2 SSU rDNA, or control plasmid. **A)** Midguts were dissected 7 days post-blood meal and representative fluorescent microscopy images were used to count the percentage of oocysts that were GFP+ out of the total population of oocysts that were mCherry+ across two biological replicates of transmission (n = 10-18 midguts). Error bars indicate standard error of the mean.
